# Computational design of metalloproteases

**DOI:** 10.1101/2025.11.20.689622

**Authors:** Anqi Chen, Kejia Wu, Hojae Choi, Preetham Venkatesh, Samuel J. Pellock, Nikita Hanikel, Hojeong shin, Declan Evans, Kieran Didi, Lars L. Schaaf, Cristina Díaz-Perlas, Brian Coventry, Donghyo Kim, Seth M. Woodbury, Pengfei Ji, Shingo Honda, Asim K. Bera, Hannah Nguyen, Alex Kang, Yanqing Wang, Xinting Li, Stacey Gerben, Lemuel Chang, Xiao Yan, Benjamí Oller-Salvia, Anthony A. Hyman, Donald Hilvert, David Baker

**Affiliations:** Department of Biochemistry, University of Washington, Seattle, WA 98195, USA; Institute for Protein Design, University of Washington, Seattle, WA 98195, USA; Graduate Program in Biological Physics, Structure and Design, University of Washington, Seattle, WA, USA; Department of Computer Science, University of Oxford, Parks Rd, Oxford OX1 3QD, UK; NVIDIA Corp., Santa Clara, USA; Cavendish Laboratory, Department of Physics, University of Cambridge, Cambridge CB3 0HE, UK; Department of Materials, Imperial College London, Exhibition Road, London SW7 2AZ, U.K.; Institut Químic de Sarrià (IQS), Universitat Ramon Llull, Via Augusta 390, Barcelona 08017, Spain; Howard Hughes Medical Institute, University of Washington, Seattle, WA 98195, USA; Department of Chemistry, Zhejiang University, Hangzhou, China; Max Planck Institute of Molecular Cell Biology and Genetics; Dresden, Saxony, 01307, Germany; Laboratory of Organic Chemistry, ETH Zurich, Zurich, Switzerland

**Author notes:** These authors contributed equally (Anqi Chen, Kejia Wu).

## Abstract

De novo design of enzymes that cleave amide bonds represents a major challenge^1–2^. Here we use RFdiffusion methods^3–4^ to de novo design proteins with water activating zinc based active sites adjacent to a peptide substrate binding pocket. Of 135 designs tested, 36% catalyze cleavage of their cognate peptide substrates at the targeted scissile bond. The best design accelerates peptide-bond hydrolysis by more than 10^8^-fold relative to the uncatalyzed reaction; four substitutions increase this acceleration to over 10^10^-fold by improving active-site preorganization^5^. We use this approach to design metalloproteases that selectively cleave the human disease associated proteins TDP-43, amyloid-β, and serum amyloid A, with rate enhancements of up to 10^9^-fold. We further design caged cytokines and receptor antagonists that are selectively activated by the designed proteases, enabling bioorthogonal control of cell state. Thus, de novo design can yield highly active, programmable proteases with applications in therapeutics, biotechnology, and bioremediation.

## Introduction

Metalloproteases utilize a metal ion as a cofactor to activate a water molecule to hydrolyze peptide bonds in proteins^6–7^. These enzymes play essential roles in biological processes such as tissue remodeling^8–9^, cell migration^10–11^, and signaling^12^. Beyond their roles in biology, proteases have widespread uses in industry and medicine in application areas ranging from food^13^ to therapeutics^14,15^ and cosmetics^16^. This broad utility has driven efforts to engineer native enzymes with tailored activities. De novo design in principle offers a complementary route to novel proteases, but to date computational design of *de novo* metallohydrolases has focused on small molecule ester substrates with activated leaving groups^1,2^. Efficient amide bond hydrolysis poses a considerably greater challenge: amides are less reactive than esters due to stronger resonance stabilization^17^, and peptide cleavage requires departure of an intrinsically poor amine leaving group. The p*K*_a_ for deprotonation of amines to form amine anions is >35^18^, compared with values <8 for typical activated esters^19,20^. Consequently, spontaneous peptide hydrolysis under physiological conditions is exceedingly slow, with half-lives on the order of hundreds of years^5^. Overcoming this kinetic barrier requires active sites that combine precise metal coordination with strategically positioned general acid–base residues to promote proton transfer and transition state stabilization^21^. Moreover, the protease-substrate interface must be designed to ensure both binding and accurate positioning of the target amide bond at the active site.

We reasoned that recent developments in deep learning methods could overcome these challenges and enable the *de novo* design of metalloproteases. Diffusive denoising methods such as RoseTTAFold Diffusion (RFdiffusion) have proven powerful in generating a wide range of new functional proteins starting from random residue distributions and sets of conditions specifying the functional constraints^22–25^. The flow-matching-based RFdiffusion2 (RFD2) method improved on RFdiffusion for enzyme design by eliminating the need for specifying the sequence positions of the catalytic residues and enabling conditioning directly on the positions of the catalytic functional groups in an idealized active site. As protease design requires not only positioning of catalytic groups but also peptide substrate recognition, we chose RFD2 for Molecular Interfaces^3^(RFD2-MI), a version optimized for design of both enzyme active sites and protein-protein interactions. We set out to explore the use of RFD2-MI for building up protein scaffolds around a minimalist metallohydrolase active site and an adjacent peptide binding site, positioning the substrate for cleavage of a defined amide bond.

### Computational design of metalloproteases

We began by constructing an idealized active site for carrying out the metalloprotease reaction. In native metalloproteases such as aminopeptidase N^26^, three residues—two histidines and a glutamate—constitute a Zn(II) binding center. The fourth ligand of Zn(II), a water molecule, is activated by another glutamate to attack the target carbonyl carbon, forming an unstable tetrahedral intermediate. This high-energy oxyanionic species and its flanking transition states are stabilized by a tyrosine hydrogen bond donor. Breakdown of the intermediate and product release are promoted by proton transfer from the protonated glutamic acid to the amine leaving group (Fig. 1a). We focused on active sites containing all five functional groups and used their positions and orientations in a co-crystal structure of aminopeptidase N with a transition state analog (TSA) (PDB 4QHP)^26^ as a geometric template. For substrate placement relative to these catalytic groups, we exploited β-strand hydrogen bonding. As a structural guide, we used AlphaFold3^27^ (AF3) to predict the structure of the enzyme-substrate (ES) complex for native astacin^28^, where the substrate binds in the protease cleft as an antiparallel β-strand. We superimposed the substrate P1-P2’ coordinates (P1 is the residue before and P1’ is the residue after the cleavage site) and the backbone atoms of three adjacent astacin residues on the catalytic residues to generate the full target active site (Fig. 1b; Supplementary Methods and Supplementary Discussion 1).

**Fig. 1:**
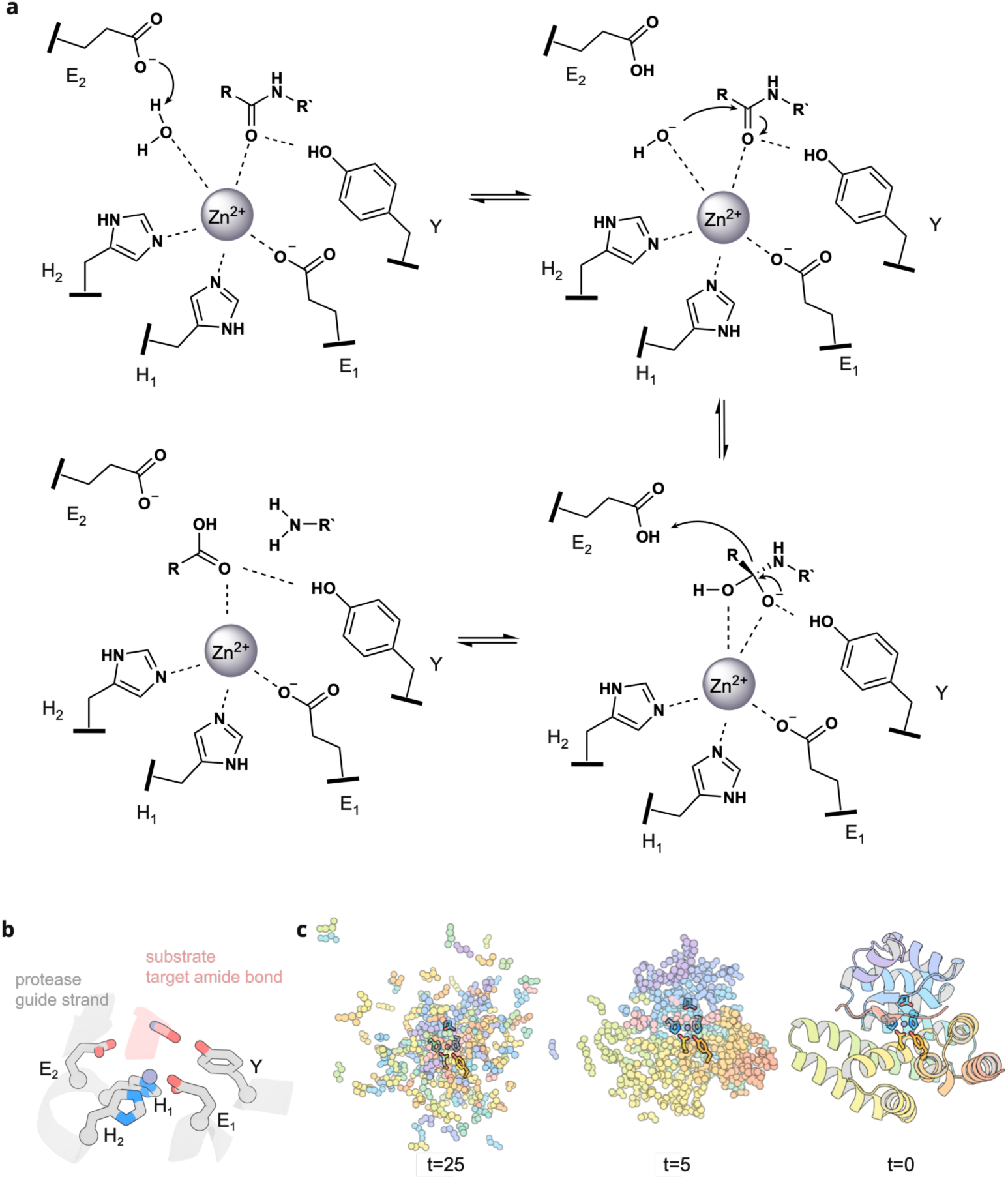
Computational design of metalloproteases. Computational design of metalloproteases. **a,** Reaction mechanism for the hydrolysis of peptide bonds by zinc proteases. **b,** Input motif for RFD2-MI. Catalytic residues were extracted from the crystal structure of aminopeptidase N in complex with a transition state analog inhibitor (pdb: 4qhp). The substrate conformation and strand pairing with the guide strand in the enzyme were extracted from an AF3 prediction of native astacin in complex with a substrate peptide. **c,** Representative RFD2-MI trajectory for backbone generation.

We used RFD2-MI to generate protein scaffolds around the constructed active site (Fig. 1c). During the design calculations, we allowed extensive conformational sampling of both the protease and the peptide substrate structures. For specified lengths of the designed enzyme (160-190AA) and substrate (8-13AA), RFD2-MI generated random residue distributions for the two chains, and progressively denoised them while conditioning on the target catalytic site and substrate binding site geometries. Many RFD2-MI trajectories yielded scaffolds incorporating the desired structural features (Supplementary Methods).

We then used a fine-tuned version of LigandMPNN, EnhancedMPNN^29^, to generate amino acid sequences that encode the designed fold, preorganize the catalytic residues, and correctly position the substrate in the binding cleft. To ensure precise placement of the target amide bond relative to the active site, we performed two-sided design^30^, simultaneously optimizing the protease and substrate sequences. Consequently, each protease design was paired with a unique substrate sequence. Designs for which AF3 predicted high-confidence structures closely matching the design models were fed back to RFD2-MI and MPNN for finer sequence and sampling to further improve structural precision and protein-substrate interface quality (Supplementary Methods).

We modeled each resulting design in three states: 1) the apo protease, to confirm correct folding; 2) the enzyme-substrate (ES) complex, to assess substrate binding and positioning of the target amide bond; and 3) the TSA complex, to evaluate transition state stabilization. For modeling the ES complex, we first used Zn(II) alone (SMILES string: [Zn+2]) as the input for modeling the metal center, a common approach for predicting Zn(II) binding complexes^27^. However, this representation omitted the catalytically essential nucleophilic water molecule, which plays a central role in zinc protease catalysis^26,28,31,32^. In a second approach, we included both Zn(II) and a coordinating water molecule (SMILES string: [Zn+2].O), enabling more direct assessment of the geometry required for nucleophilic attack. To model the transition state, we used a phosphonamidate TSA^33^. Because many crystal structures have been solved for similar TSA complexes^26,28,31,32^, these complexes are better modeled by machine learning software trained on the PDB than the actual transition state. We evaluated designs in all three states according to x prediction confidence, interface quality, and active site geometry. Synthetic genes were obtained encoding 95 designs from the Zn(II)-only modeling strategy and 40 designs from the Zn(II)-water strategy (see Extended Data Fig.1 and Supplementary Methods, Supplementary Discussion 1,2, Table S1 for the full details on the computational pipeline).

### Experimental validation of two-sided designed proteases

As each design has a unique protease-substrate pair, we first screened activity using a fusion construct in which the protease was covalently linked to the substrate so both could be encoded by the same synthetic gene. The construct additionally included an mScarlet tag to facilitate detection of the molecular weight shift upon cleavage, and a Strep-tag for purification (Fig. 2a; Extended Data Fig. 2a). We refer to this format as a “cis” screen because cleavage occurs intramolecularly. All 135 designed enzyme-substrate fusions were overexpressed in *E. coli* and purified by Strep-tag affinity chromatography and size-exclusion chromatography (SEC). Of these, 131 were soluble, and 127 migrated at the expected molecular weight of the full-length construct on SDS-PAGE (Fig. S1).

**Fig. 2:**
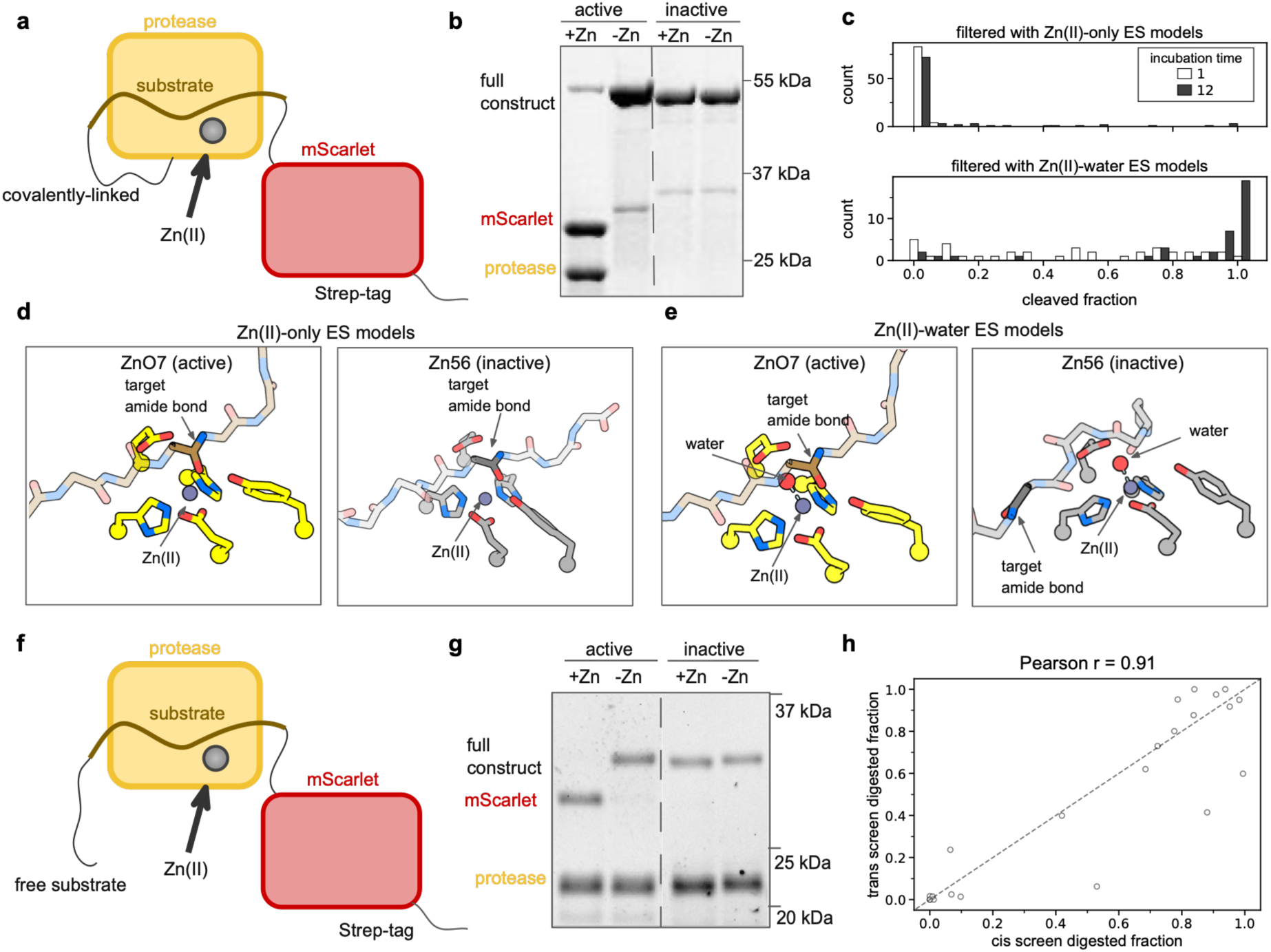
Experimental screening. **a,** Schematic illustration of the fusion construct used in the cis screen. The proteases were expressed as a genetic fusion with their designed substrate peptide sequence, followed by a C-terminal mScarlet. **b,** Representative screening outcomes for an active and an inactive design after 1 h at 37 °C in the cis assay. An active design exhibits Zn-dependent cleavage at the correct site as assessed by mass spectrometry (Table S2). In contrast, an inactive design remained uncleaved. This experiment was performed three times independently with similar results. See Fig. S1 for gel source data **c,** Digested fraction of the substrate measured by SDS-PAGE gel densitometry. The hit rate significantly improved when the nucleophilic water was explicitly modeled in the ES complex. **d,** Zn(II)-only models of the ES complexes did not differentiate between active (ZnO7) and inactive (Zn56) designs. **e,** Zn(II)-water models of the ES complexes clearly differentiated active (ZnO7) from inactive (Zn56) designs, based on the displaced substrate in the latter. **f,** Schematic illustration of the protein constructs used in the trans screen. The proteases and their designed substrates were expressed as separated chains. The substrate sequence was expressed as a genetic fusion with an N-terminal (GS)_15_ linker and a C-terminal mScarlet. **g,** Representative screening outcomes for an active and an inactive design in the trans assay after 1 h at 37 °C. An active design exhibited Zn-dependent cleavage at the correct site. In contrast, an inactive design exhibited no cleavage. This experiment was performed three times independently with similar results. See Fig. S1 for gel source data. **h,** Relationship between the digested fractions of the cis and trans assays, both performed at 37 °C for 1 h.

To evaluate metalloprotease activity, the designs were incubated at 37°C in the presence or absence of Zn(II), and cleavage was assessed by SDS-PAGE. Active designs cleaved the constructs into lower-MW N-terminal protease and C-terminal mScarlet fragments, whereas inactive designs remained as the full-length fusion (Fig. 2b). Cleavage was measured after 1 h and 12 h and quantified by densitometric analysis of substrate and product bands (Fig. S2). For designs selected using the Zn(II)-only strategy, 3/89 showed >10% cleavage after 1 h, and 14/89 showed >10% cleavage after 12 h. For designs selected using the Zn(II)-water strategy, 31/38 showed >10% cleavage after 1 h, and 35/38 showed >10% cleavage after 12 h (Fig. 2c). Hereafter, designs from the first class are designated Zn1, Zn2, etc., whereas designs from the second class are designated ZnO1, ZnO2, etc. The overall success rate for all 135 designs, as defined by >10% cleavage after 12 h at 37°C, was 36% (15% for the Zn designs and 88% for the ZnO designs). In every case, mass spectrometry confirmed cleavage at the intended site (Table S2, Fig. S3).

To understand the difference between the Zn and ZnO designs, we compared the ES structure models of the inactive design Zn56 and the active design ZnO7. In models generated with Zn(II) alone, the substrates of both designs were positioned correctly in their respective binding clefts, providing no basis for differentiating active and inactive designs (Fig. 2d). In the Zn(II)-water models, however, the catalytic water in Zn56 sterically prevented productive substrate binding, displacing the target amide bond by several ångströms (Å). In contrast, the substrate in ZnO7 was predicted to bind correctly at the active site, with the scissile amide correctly oriented for nucleophilic attack(Fig. 2e). Thus, Zn(II)-water modeling successfully distinguished the active design ZnO7 and would have eliminated the inactive design Zn56, suggesting that explicitly including mechanistically essential species—such as the catalytic water—can substantially improve design success.

Although the cis screen efficiently identified designs with functional active sites, it did not assess proper independent folding of the apo enzyme or intrinsic substrate binding. We therefore evaluated 22 hits with protease and substrate produced as separate polypeptides (Fig. 2f, Extended Data Fig. 2b-c). The peptide substrate was fused to an N-terminal (GS)_15_ linker and a C-terminal mScarlet. Protease and substrate constructs were expressed separately in *E. coli* and purified by Strep-tag affinity chromatography followed by SEC. For each design, purified protease and substrate were co-incubated for 1 h at 37°C with and without supplemental Zn(II). Successful cleavage removed an ∼3 kDa N-terminal fragment from the substrate, detectable by SDS-PAGE (Fig. 2g; Fig.S4-5). Fourteen designs, representing five unique scaffolds, exhibited >10% trans cleavage after 1 h at 37°C (Table S3). Cleavage activities in the cis and trans formats were strongly correlated with a Pearson’s r value of 0.91 (Fig. 2h), validating the screening workflow (Table S4).

### Validation of designed mechanism

For each scaffold, we chose one design for detailed characterization (Fig. 3a). To determine the functional contributions of the designed catalytic residues, we used site-directed mutagenesis to knock out each zinc-binding residue, the catalytic base, and the oxyanion-stabilizing tyrosine. For all five designs, mutating any of the zinc-binding residues or the catalytic base reduced the activity of the design by at least sevenfold, as quantified by densitometric analysis of SDS-PAGE images (Fig. 3b; Fig. S6-9). In four of the five designs, replacing the oxyanion-stabilizing tyrosine with phenylalanine also reduced activity by at least eightfold. Thus, the catalytic activity of the designs arises from the designed active sites.

**Fig. 3:**
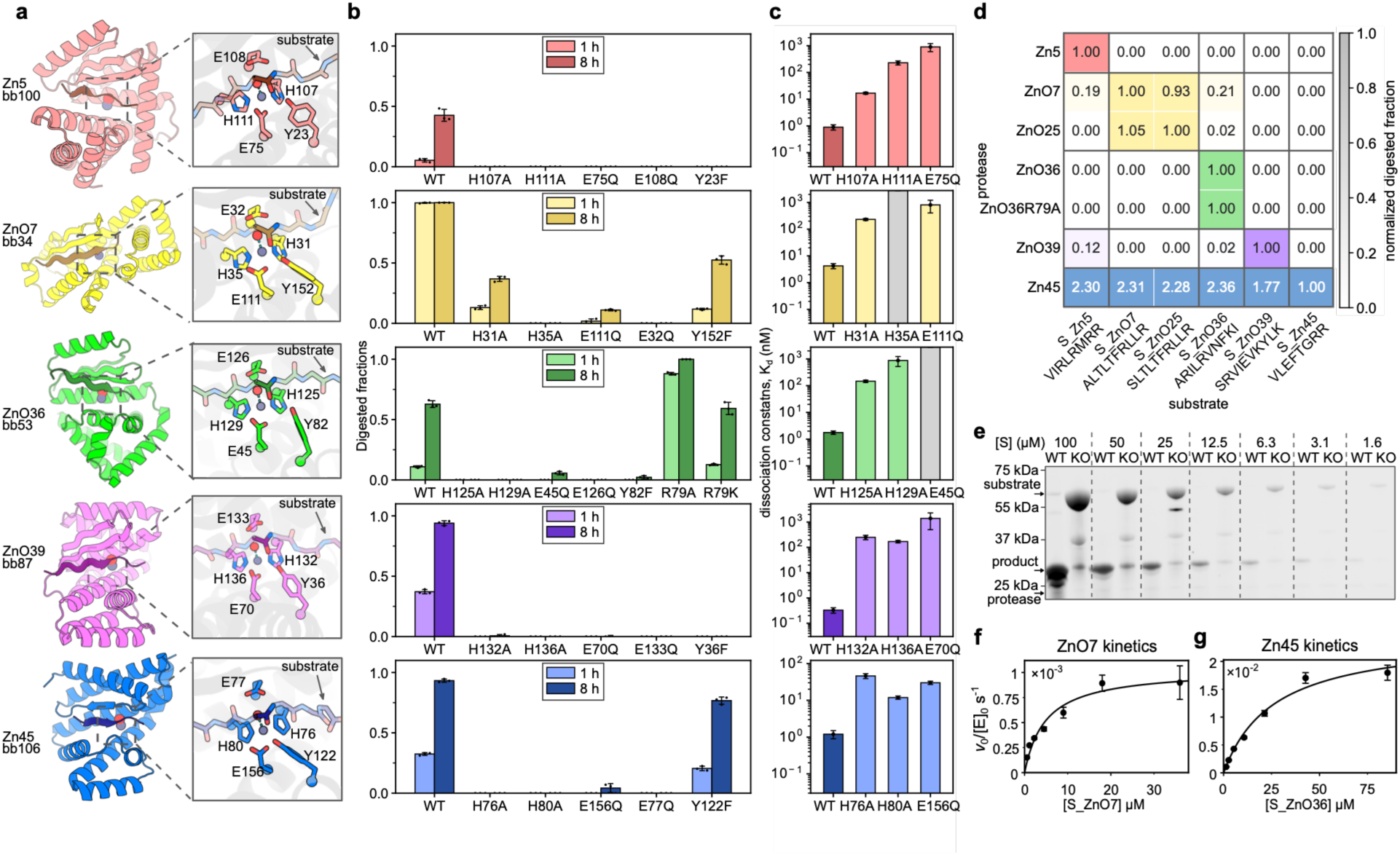
Functional characterization of protease designs. **a,** AF3 models of active protease designs with close-up views of the active sites from five unique scaffolds (TM-score < 0.23 between any pair). **b,** Impact of mutating five catalytic residues on design activity measured by SDS-PAGE gel densitometry. For ZnO36, two additional substitutions, R79A and R79K, were investigated due to the proximity of R79 to the tetrahedral intermediate/transition state oxyanion (Fig. S9). Measurements were made in triplicate. Bars and error bars represent mean ± std. **c,** Assessment of Zn(II) dissociation constant for the designed proteases and the respective knockout variants. Gray bars with heights above the upper boundary of the plots indicate weak Zn(II) binders that are beyond the dynamic range of the assay, suggesting *K*_d_ values above 2 μM. Bars and error bars represent the dissociation constant (*K*_d_ ± std) fitted by binding isotherms across 15 concentrations where each concentration was measured once. The experiment was performed twice with similar results. **d,** Cross-reactivity heatmap of 7 proteases (rows) with 6 designed substrates (columns). Designs from the same backbone were labeled with the same color. ZnO25 is a one-shot design with the same backbone as ZnO7 but differing in sequence at 39 positions. The mean of three measurements is shown in the heatmap. **e,** SDS-PAGE gel analysis of 500 nM Zn45-catalyzed cleavage of the substrate designed for ZnO36 at different substrate concentrations, [S]. This experiment was performed three times independently with similar results. Full gel image in Supplementary Fig. S17. **f,** Kinetics of ZnO7-catalyzed cleavage of its cognate substrate, and **g,** of Zn45-catalyzed cleavage of the substrate designed for ZnO36. *v*_0_: reaction rate in the initial linear region (nM/s). [E_0_]: enzyme concentration, 500 nM. Measurements were made in triplicate. Black dots and error bars represent mean ± std.

Competition experiments with a ratiometric dye revealed that the designed proteases bind Zn(II) with dissociation constants (*K*_d_) ranging from 10^-10^ to 10^-8^ M (Fig. 3c; Fig.S10). ZnO39 had subnanomolar Zn(II) affinity, exceeding that reported for previously designed zinc enzymes^2,34–35^ and approaching the affinities of some native zinc enzymes^36^. The high Zn(II) affinities of these *de novo* proteases could be advantageous in applications where supplementation with exogenous Zn(II) is impractical, for example in therapeutic settings.

### Design selectivity and activity

As we performed two-sided sequence design of both enzyme and substrate, each designed protease has a uniquely designed substrate sequence. We assessed cross-reactivity among proteases from the five scaffold families. Four of the five designs preferentially cleaved their cognate substrate (Fig. 3d; Fig. S11-12). This is notable because substrate selectivity was not explicitly enforced, and designs predicted to cleave alternative substrates were not excluded. Instead, specificity appears to have emerged directly from the two-sided design process, which produced highly complementary protease–substrate interfaces.

The fifth design, Zn45, displayed high activity against all tested substrates (Fig. 3d). This experimentally observed cross-reactivity is consistent with the AF3 metrics used during the design process (Extended Data Fig. 3). To quantify cross-reactivity, we examined cleavage of S_ZnO36, the cognate substrate of ZnO36. Zn45 almost completely digested S_ZnO36 after 12 h at 37°C over substrate concentrations ranging from 1.6 μM to 100 μM (Fig. 3e, Fig. S13). At the highest substrate concentration, each Zn45 molecule turned over ∼200 molecules of S_ZnO36 overnight. An AF3 model of the Zn45 Michaelis complex containing S_ZnO36 placed the main-chain carbonyl of the seventh substrate residue precisely in the Zn45 active site, coinciding with the MS-validated cleavage site (Fig.S14).

We characterized steady-state kinetics for two protease-substrate pairs: ZnO7 on its cognate substrate, which exhibited the highest activity toward its designed substrate in the SDS-PAGE cleavage assays, and Zn45 acting on the ZnO36 substrate, the pair displaying the highest overall activity. To acquire continuous reaction measurements over time, we monitored fluorescence resonance energy transfer (FRET) from substrates fused to N-terminal GFP and C-terminal mScarlet fluorophores (Extended Data Fig. 2d; Fig. S15-17). For ZnO7 cleavage of its cognate substrate, we determined a *k*_cat_ of 0.0010±0.0001 s^-1^, a *K*_M_ of 4.9±1.2 μM, and *k*_cat_/*K*_M_ of 210±60 M^-1^s^-1^ (Fig. 3f). Zn45 cleavage of the ZnO36 substrate was substantially faster, with *k*_cat_ = 0.025±0.002 s^-1^, *K*_M_ = 26±5 μM, and *k*_cat_/*K*_M_ = 900±200 M^-1^s^-1^ (Fig. 3g). The *k*_cat_ for Zn45 corresponds to a >10^8^-fold rate acceleration relative to uncatalyzed peptide bond hydrolysis^5^.

### Structural validation

We determined X-ray crystal structures for Zn5 and ZnO7 (Fig. 4). The Zn5–Zn(II) structure (resolution=3.41 Å, PDB 11CE), and the ZnO7–Zn(II)–substrate complex (resolution=2.37 Å, PDB 11CM) closely matched their design model, with Cα RMSDs of 0.82 Å and 1.35 Å, respectively (Fig.4a-b, Table S5).

**Fig. 4:**
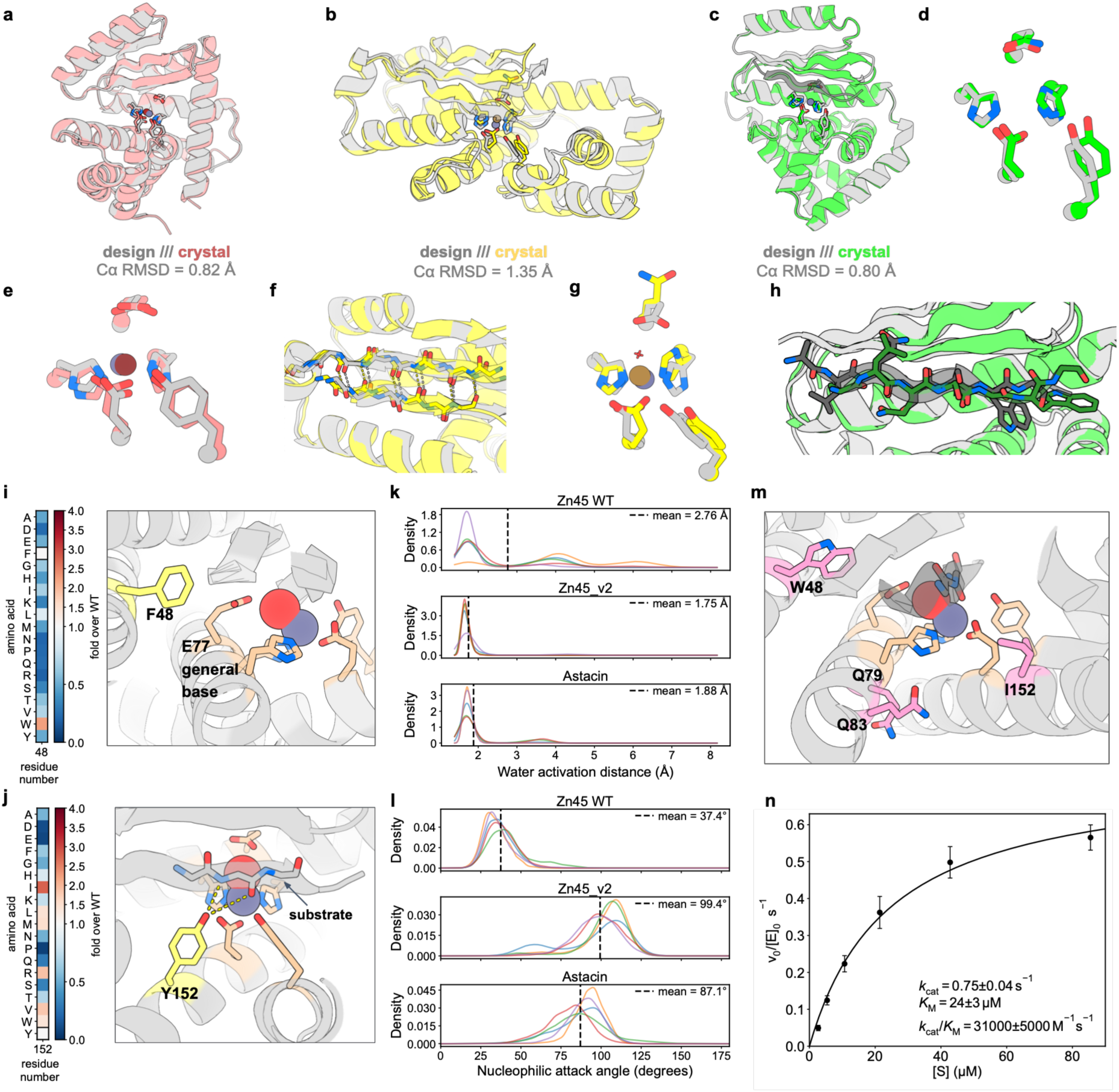
Characterization of structure and active site preorganization. **a-c,** structural superposition of design models (gray) and crystal structures for Zn5 (a, red), ZnO7 (b, yellow) and TDPr3 (c, green). Active sites are shown as sticks. The Zn(II) ions are shown in dark gray in the design models, dark red in the crystal structure of Zn5 and dark yellow in the crystal structure of ZnO7. **d,e,g** Active site overlays of the crystal and the design model for TDPr3 (d), Zn5 (e), ZnO7 (g). The catalytic water in the active site of the ZnO7 crystal in **g** is shown as a red cross. **f,** Zoomed-in view of the substrate-protease β-strand pair in the superposition of the ZnO7 crystal and design model. **h,** Zoomed-in view of substrate registration in TDPr3. The substrate is shifted by one residue between the crystal and the design. **i-j,** activity heatmap of single variants for Zn45, and the positions of mutated residues (yellow) with respect to the catalytic residues (wheat) in the AF3 prediction. **k,** Distribution of the distance between the proton-abstracting oxygen in the general base and the closest water hydrogen, in the simulations of wild-type Zn45 (top), astacin (middle) and Zn45_v2 (bottom). **l,** Distribution of the angle between the water oxygen, the P1 carbonyl carbon, and the P1 carbonyl oxygen, in the simulation of wild-type Zn45 (top), astacin (middle) and Zn45_v2 (bottom). For both k and l, solid curves of different colors represent five replicas. Black dashed lines represent the mean of all five replicas. **m,** Beneficial mutations in Zn45_v2. **i,** Kinetics of Zn45_v2-catalyzed cleavage of the substrate of ZnO36. *v*_0_: reaction rate in the initial linear region (nM/s). [E_0_]: enzyme concentration, 50 nM. Measurements were made in triplicate. Black dots and error bars represent mean±std.

In the Zn5–Zn(II) complex, the five catalytic residues adopt conformations closely matching the design model (RMSD=0.9 Å). The Zn(II) ion is clearly resolved at the intended coordination site at full occupancy (RMSD=0.3 Å; Fig. 4e). An apo structure of Zn5 (at 3.57 Å resolution) revealed a preorganized catalytic configuration in the absence of the Zn(II) cofactor (Extended Data Fig. 4a), validating the accurate design of the catalytic residues and the metal coordination geometry.

To capture the Michaelis complex of ZnO7, we substituted the E32 general base with a glutamine to abolish activity and stabilize the substrate bound in the protease cleft. In the resulting crystal structure, the substrate peptide binds the protease in the designed β strand-pairing conformation (Fig. 4f). A Zn(II) ion is resolved at the designed coordination site with full occupancy (RMSD = 0.3 Å) along with a bound water molecule, corresponding to the nucleophilic water in the Michaelis complex that initiates catalysis (Fig.4g). The substituted Q32 residue adopts a side-chain rotamer distinct from the catalytically competent rotamer predicted in the design model or the crystal structures of other designs, suggesting stabilizing the correct general base rotamer as a direction for improvement for this design (Supplementary Discussion 3.1 and Fig. S18, Extended Data Fig. 4).

Structural similarity of the designs with proteins in the Protein Data Bank (PDB) was assessed using Foldseek^37^. No fold-level matches were identified: the highest TM-scores were below 0.42, whereas scores above 0.5 generally indicate structural similarity (Extended Data Fig. 5). Sequences of active designs were also searched against the NCBI non-redundant database using BLASTP. The best hits had E-values ranging from 0.34 to 33, indicating no sequence similarity to known proteins (E-values below 0.05 indicate sequence similarity; Table S6).

**Fig. 5:**
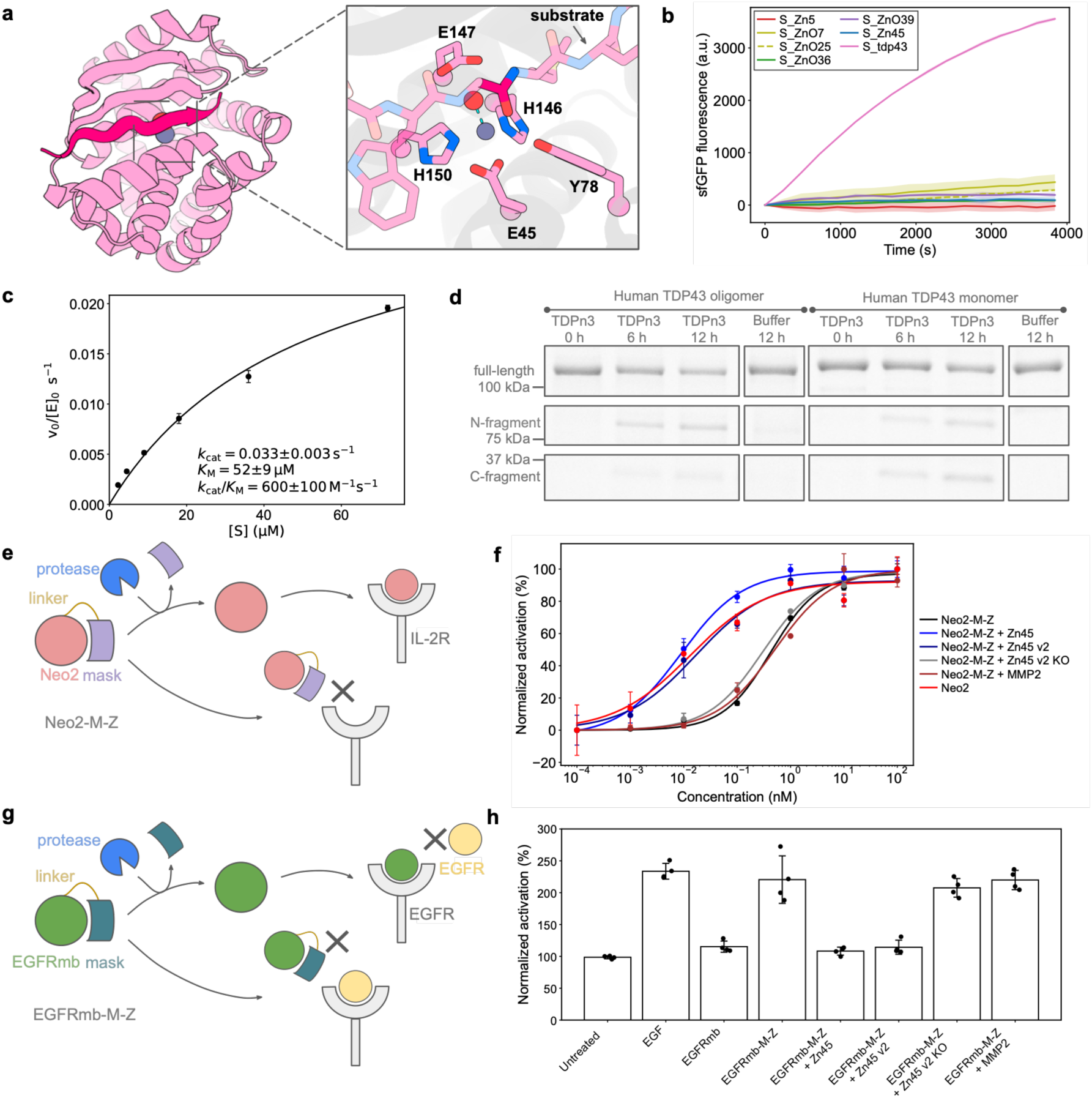
De novo protease for cleavage of human protein and for conditional activation of masked cytokine mimic. **a,** TDPn3 model with zoomed-in view of the active site. **b,** Substrate specificity of TDPr3 against the sfGFP-mScarlet substrates of all designs presented in this work. Measurements were made in triplicate. Solid lines and shaded regions represent the mean ± std. **c,** Kinetics of TDPn3-catalyzed cleavage of the target TDP-43 epitope. *v*_0_: reaction rate in the initial linear region (nM/s). [E_0_]: enzyme concentration, 500 nM. Measurements were made in triplicate. Black dots and error bars represent mean ± std. **d,** Cleavage of full-length human TDP-43 in vitro. This experiment was performed three times independently with similar results. See Fig. S24 for gel source data. **e,** Schematic illustration of the conditional activation of a masked cytokine mimic, Neo2-M-Z, by de novo proteases. **f,** Normalized activation percentage of HEK-Blue IL-2 reporter cells by Neo2-M-Z treated with Zn45, Zn45_v2, Zn45_v2_KO, MMP2 and by Neo2. Measurements were made in triplicate. Data points and error bars represent mean ± std. Solid lines indicate sigmoidal fits to the raw data. **g,** Schematic illustration of the conditional activation of a masked EGFR antagonist, EGFRmb-M-Z, by de novo proteases. **h,** Normalized phospho-ERK percentage of cells treated with Zn45, Zn45_v2, Zn45_v2_KO, MMP2 and by EGFP-mb. Measurements were made in quadruplicate. Data points and error bars represent mean ± std.

### Origin of activity discrepancies between the designs and native metalloproteases

Despite the high structural accuracy of our de novo metalloproteases, their activities are lower than those of the native metalloprotease from which the theozyme was derived^38–39^. Thus, sub-ångström agreement with a design model does not guarantee optimal catalysis: small differences in catalytic-residue geometry and the surrounding chemical environment can have large effects along the reaction coordinate.

To examine these effects, we used machine learning force fields (MLFF)^40^ to calculate enthalpic activation barriers for the likely rate-limiting proton-abstraction and nucleophilic-attack steps^41^ for Zn45, ZnO7, and native astacin (Fig. 1a). Simulating the catalytic residues, substrate, and all residues with at least one atom within 5 Å of the substrate (Supplementary Discussion 4) we calculated the activation barriers for Zn45 and ZnO7 to be 3.6 and 15.1 kcal/mol higher than those of astacin (Fig. S19), indicating less favorable reaction energetics in the designed active sites. Although ZnO7 more closely matched its design model than Zn45, Zn45 had the higher *k*_cat_ (Fig. S18b); the MLFF computed barriers qualitatively tracked the experimental activity trend, despite not quantitatively reproducing the magnitude of the rate differences (Supplementary Discussion 4). Explicit assessment of the catalytic energy landscape could improve activity prediction and guide optimization of one-shot designs.

To identify sub-optimal features of the designs not captured by the MLFF calculations, we systematically mutated 21 positions within 4 Å of the five catalytic residues or the substrate P2, P1, and P1’ residues in the Zn45–S_ZnO36 model, to each amino acid except cysteine (Extended Data Fig. 6). At eleven positions, the designed amino acid was already optimal, whereas substitutions at the remaining ten positions increased activity by at least 30% (Extended Data Fig. 6c), revealing limitations in the original design. Substitution of F48, adjacent to the general base E77 (Fig. 4i), with tryptophan increased *k*_cat_ tenfold (Extended Data Fig. 6d), likely by improving the positioning of E77. Y152, introduced during MPNN sequence design, forms hydrogen bonds with the P1 backbone that compete with interactions in the designed oxyanion hole; replacing Y152 with hydrophobic or bulky residues (I, L, M, V, W, R) increased activity (Fig. 4j), suggesting that Y152 preferentially stabilized the ground state rather than the transition state. A similar effect may account for the beneficial R79A substitution in ZnO36 (Supplementary Discussion 5).

Zn45_v2 which combines F48W and Y152I with A79Q and Y83Q, which may help refine the positioning of the α-helix containing the general base and two histidines (Extended Data Fig. 6e, Extended Data Fig. 7), displayed a *k*_cat_ of 0.75±0.04 s^-1^, a *K*_M_ of 24±3 μM, and a *k*_cat_/*K*_M_ of 31000±5000 M^-1^s^-1^, a 30-fold improvement in both *k*_cat_ and *k*_cat_/*K*_M_ relative to Zn45 (Fig. 4m,n). Zn45_v2 accelerates peptide bond hydrolysis by 1.4×10^10^-fold relative to the uncatalyzed reaction, achieves more than 3500 turnovers per active site (Extended Data Fig. 7d), and has kinetic parameters comparable to some native metalloproteases (Table S7) and other well-characterized proteases^42–44^. Neither AF3 nor MPNN predicted the beneficial effects of the individual or combined mutations, however, highlighting the need for further methodological improvement (Extended Data Fig. 6f-g, Extended Data Fig. 7h, Supplementary Discussion 6).

Molecular dynamics (MD) simulations of Michaelis complexes provided a structural rationale for the improved activity of Zn45_v2. In Zn45, the catalytic glutamate E77 sampled diverse rotamers, producing large fluctuations in the distance between its closest oxygen atom and the water proton, thereby disfavoring proton abstraction. In Zn45_v2 and astacin, by contrast, the general base rotamer was substantially more preorganized, maintaining average oxygen-hydrogen distances of 1.75 Å and 1.88 Å, respectively (Fig. 4k). The scissile amide was also poorly preorganized in Zn45: the angle defined by the water oxygen, target carbonyl carbon, and carbonyl oxygen averaged 37.4°, compared with 99.4° for Zn45_v2 and 87.1° for astacin, which are much closer to the preferred trajectory for nucleophilic attack^45^ (Fig.4l). Collectively, the mutagenesis and MD data implicate incomplete preorganization of both catalytic groups and substrate as a principal source of the residual activity gap between the designed and native enzymes.

Despite these gains, Zn45_v2 remains less active than the best native proteases^38–39^. Site-directed substitutions showed that the Zn(II)-chelating residues and general base are essential, whereas replacement of the putative oxyanion-stabilizing residue Y122 caused only a ∼10-fold loss of activity. Because analogous oxyanion-stabilizing residues typically contribute >50-fold to native protease catalysis^46–48^, this result suggests that the oxyanion stabilizer remains suboptimally positioned; improvement in this region may be feasible, consistent with the beneficial effect of the nearby V121I substitution (Extended Data Fig. 6, Fig. S20). The pH dependence of Zn45_v2 resembles the bell-shaped profiles reported for thermolysin^49^ and stromelysin-1^50^, but the acidic limb is shifted higher by ∼1–2 pH units. Because this transition is commonly associated with activation of the general base–Zn(II)–water system, the shift suggests that the local chemical environment in Zn45_v2 differs from that of native metalloproteases and may still limit efficient nucleophilic attack.

### Design of proteases cleaving peptides in disease-relevant targets

Having established the ability to design proficient proteases, we next explored potential applications in targeting disease-associated proteins and constructing synthetic protease-activated control systems. We selected three human proteins as protease targets whose aberrant aggregation is implicated in disease: TAR DNA-binding protein 43 (TDP-43), serum amyloid A (SAA), and the amyloid-β (Aβ) peptide^51–53^.

For TDP43, we targeted a C-terminal segment (residues 320-350) to disrupt pathological assemblies while preserving essential protein functions^54^ (Extended Data Fig. 8). Initial efforts to alter the substrate specificity of ZnO36 through partial diffusion followed by EnhancedMPNN redesign produced a protease, TDPr3, that cleaved the target sequence but retained a strong preference for the original cognate substrate, likely due to suboptimal interface design (Fig. 4c,d,h, S21,22; Extended data figure 8a-c; Supplementary Methods).

By contrast, direct de novo design of a scaffold for TDP-43 peptide using atom-level RFDiffusion3^4^ successfully accommodated the Zn(II)-water theozyme and introduced substrate-specific interactions (Fig S23, Supplementary Methods). The most active catalyst, TDPn3 (Fig. 5a), had substantially greater activity toward the TDP-43 target sequence than toward the other substrates tested (Fig. 5b). Its steady-state parameters for this substrate–*k*_cat_ = 0.033±0.003 s^-1^, *K*_M_=52±9 μM, and *k*_cat_/*K*_M_ =640±130 M^-1^s^-1^ (Fig. 5c)–were comparable to those of the best two-sided design, but with much greater substrate specificity (Fig. 3d). These results demonstrate that one-shot de novo design can simultaneously achieve appreciable catalytic activity and substrate specificity.

When incubated with full length TDP-43, in both monomeric and oligomeric forms, TDPn3 generated cleavage products of the expected sizes, as confirmed by MS (Supplementary Table S2, S8, Fig. 5d, Fig. S24). No cleavage was detected in buffer-only or base-knockout controls (Extended Data Fig. 8f). Cleavage of full-length TDP-43 was nevertheless slower than cleavage of the synthetic sfGFP-mScarlet substrate, potentially owing to liquid-liquid phase separation or precipitation of TDP-43^55^ (Extended Data Fig. 8e).

We also succeeded in designing metalloproteases cleaving the SAA-derived peptide SSRSFFSFLG and the Aβ-derived KGAIIGLMVG peptide, further demonstrating the generalizability of the one-shot design pipeline (Extended Data Fig. 9). Both sets of designs exhibited their highest activity toward the respective intended substrate relative to other substrates examined in this study.

### Conditional activation of a cytokine mimic and a receptor antagonist by a designed metalloprotease

In addition to targeting disease-associated proteins, de novo protease could enable synthetic, protease-gated therapeutic circuits for precision medicine, analogous to antibody-directed enzyme prodrug therapy (ADEPT)^56^. To evaluate whether designed metalloprotease could conditionally activate therapeutic proteins, we constructed a masked version of Neo2, a previously reported de novo IL2 mimic^57^. A Neo2-binding minibinder generated with RFD2 to mask the IL2R-binding interface was fused to Neo2 through a linker containing the S_ZnO36 cleavage sequence. The resulting masked cytokine, Neo2-M-Z (Table S9), was used in a HEK-Blue IL-2/IL-15 reporter assay, with or without protease, to assess Neo2 activation and signaling through IL2R (Fig. 5e). In the absence of protease, Neo2-M-Z exhibited an EC50 of ∼0.5 nM. In contrast, treatment with Zn45 or Zn45_v2 shifted the EC50 to ∼0.01 nM, comparable to that of unmasked Neo2 and corresponding to a ∼50-fold activation window. In contrast, Neo2-M-Z was not activated by matrix metalloprotease 2 (Fig 5f), a tumor-enriched human protease commonly used as a trigger in prodrug-activation strategies^58^.

We further validated this circuit with a masked de novo EGFR antagonist^59^. Treatment of the masked antagonist, EGFRmb-M-Z (Table S9), with Zn45 or Zn45_v2 activated the antagonist, inhibiting EGFR signaling and downstream ERK phosphorylation. This unmasking effect was abolished by catalytic residue knockout of Zn45_v2 and was not observed following treatment with MMP2 (Fig. 5g,h, Fig. S25). In principle, adoptive cell therapies engineered to display such bioorthogonal de novo proteases on their surface could enable localized activation of masked biologics at disease sites.

## Discussion

Our results show that *de novo* enzyme design has advanced well beyond model reactions on activated substrates. Using a Zn(II) ion to activate a water in active sites containing three Zn(II)-coordinating residues and additional water-activating and oxyanion-stabilizing residues, we achieved >10^8^-fold rate accelerations for peptide bond cleavage relative to uncatalyzed hydrolysis in aqueous solution. Both codesigning the substrate and protease and designing a protease for a prescribed substrate sequence yielded substantial substrate selectivity without explicit negative design against alternative substrates.

Despite these advances, substantial opportunities exist to improve our design methodology. Although the turnover rates of enzymes straight out of the computer are still 10^3^-10^4^-fold lower than those of the most efficient natural proteases, characterization of our metalloprotease designs highlights strategies for further enhancement: more precise active site placement; incorporation of energetic alongside geometric design criteria; improved preorganization of catalytic residues and substrate; and optimization of the local electrostatic environment. Notably, increasing preorganization with a few mutations narrowed the activity gap to top-performing natural metalloproteases to two orders of magnitude.

The ability to design selective proteases for prescribed epitopes could be transformative for many potential applications. Unlike many native proteases, which recognize short 2-5 residue motifs^60^, our designs bind 8–13-residue substrates, creating an extended recognition interface that can encode substantial sequence and conformational selectivity. Such tunable specificity could enable cleavage of viral polyprotein junctions, selective dismantling of disease-associated assemblies while sparing essential protein pools, rewiring of signaling pathways, and engineering protease-controlled cassettes for synthetic biology applications—greatly reducing off-target risks. Beyond therapeutics, programmable proteases could support safe and more sustainable solutions across agriculture, food, and cosmetics, including the control of pests and pathogens and improvements to product texture and stability. Together, these advances point toward a future in which proteolysis becomes a programmable modality for biotechnology, medicine, and engineered living systems.

## Methods

For completeness and to centralize the methodological details, the computational and experimental methods used in this study are described in the Supplementary Information.

## Data Availability

All data generated or analyzed in this study are available in the main text, the Supplementary Materials, or the Zenodo repository: https://doi.org/10.5281/zenodo.22654831. Crystal structures have been deposited in the Protein Data Bank (PDB; https://www.rcsb.org/) under accession codes 11CE, 11CM, 11CP, 11CR, 11CU, 11CY, and 11DU.

## Code Availability

The code used in this study is available in the Zenodo repository: https://doi.org/10.5281/zenodo.22654831.

## Author Contributions

A.C., K.W., P.V., D.H., and D.B. conceptualized the project. A.C., K.W., H.C., P.V., S.J.P. developed the computational design pipeline. N.H. performed the Zn(II)-dissociation constant experiments. H.S. designed and performed the conditional cytokine mimic activation experiments. B.O.-S. and C.D.-P. designed and performed the masked conditional antagonist experiments. B.C. trained RFD2-MI. D.E. performed the MD simulations. K.D. and L.L.S performed the MLFF calculations. A.K.B, H.N., A.K., Y.W. performed protein crystallography. P.J., D.K., S.M.W., and S.H. contributed ideas and code. A.C., K.W., and L.C. produced and characterized the designs. A.C. and K.W. analyzed the experimental results. X.L. performed intact protein mass spectrometry. S.G. provided training on equipment and advice on experimental protocols. X.Y and A.A.H produced and characterized full-length human TDP-43. K.W., D.H and D.B. supervised the project. A. C., K.W., D.H., D.B. wrote the manuscript. All authors reviewed, commented and edited the manuscript.

## Funding Sources

### This work was funded by the following

Bill and Melinda Gates Foundation INV-043758. Defense Threat Reduction Agency Grant HDTRA1-19-1-0003. Helmsley Charitable Trust Type 1 Diabetes (T1D) Program Grant # 2019PG-T1D026. Howard Hughes Medical Institute. The Audacious Project at the Institute for Protein Design. The Open Philanthropy Project Improving Protein Design Fund. The National Institutes of Health’s National Institute of Allergy and Infectious Disease, grant R0AI160052. The National Institutes of Health’s National Institute on Aging, grant R01AG063845. The Matchmakers team supported by the Cancer Grand Challenges partnership funded by Cancer Research UK (CGCATF-2023/100008) and the National Cancer Institute 1(OT2CA297288-01), and the Mark Foundation for Cancer Research. The Advanced Research Projects Agency for Health APECx Program Award No. 1AY1AX000036. The Gates Foundation grant INV-043758. The National Institute for Allergy and Infectious Diseases, National Institutes for Health grant number 1U19AI181881-01. Wu Tsai Translational Investigator Fund. Grantham Foundation for the Protection of the Environment. Schmidt Family Foundation. MICIU/AEI/10.13039/501100011033 and FEDER, EU (PID2023-151988OB-I00). “la Caixa” Foundation (LCF/BQ/PR25/12110008). The National Natural Science Foundation of China (NSFC) 22377107. Wolfson College, Cambridge through a Junior Research Fellowship. ARIA, DSIT and Pillar VC under the Encode AI for Science Fellowship.

## Acknowledgements

We thank L. Goldschmidt for maintaining the computational, and K. Van Wormer, R. Ticzon, H. Nunez-Ortegaand, E. Weston for maintaining the wet lab resources at the Institute for Protein Design. We thank I.Kalvet, H. Yu, B. Belmont, A. Broerman, B. Liu, L.T. Yu, C. Liu, W. Ahern, F. Xue, C. Wang for helpful discussions during the development of the project. We thank D. Whittington and R. Lundeen for helping with mass spectrometry. We thank I. Kalvet and Y. Liu for providing reagents. We thank M. A. Kennedy for reading and editing drafts of the manuscript. We thank L. Stewart for managing the Institute for Protein Design and securing funding resources.

This work is based upon research conducted at the Northeastern Collaborative Access Team beamlines, which are funded by the National Institute of General Medical Sciences from the National Institutes of Health (P30 GM124165). The Eiger 16M detector on the 24-ID-E beam line is funded by a NIH-ORIP HEI grant (S10OD021527). This research used beamtime awards (DOI: doi.org/10.46936/APS-190993/60014759) from the Advanced Photon Source, a U.S. Department of Energy (DOE) Office of Science User Facility operated for the DOE Office of Science by Argonne National Laboratory under Contract No. DE-AC02-06CH11357. This research used resources 17-ID-1 (AMX)/17-ID-2 (FMX) of the National Synchrotron Light Source II, a U.S. Department of Energy (DOE) Office of Science User Facility operated for the DOE Office of Science by Brookhaven National Laboratory under Contract No. DE-SC0012704. The Center for BioMolecular Structure (CBMS) is primarily supported by the National Institutes of Health, National Institute of General Medical Sciences (NIGMS) through a Center Core P30 Grant (P30GM133893), and by the DOE Office of Biological and Environmental Research (KP1605010).

## Extended data

**Extended Data Figure 1.**
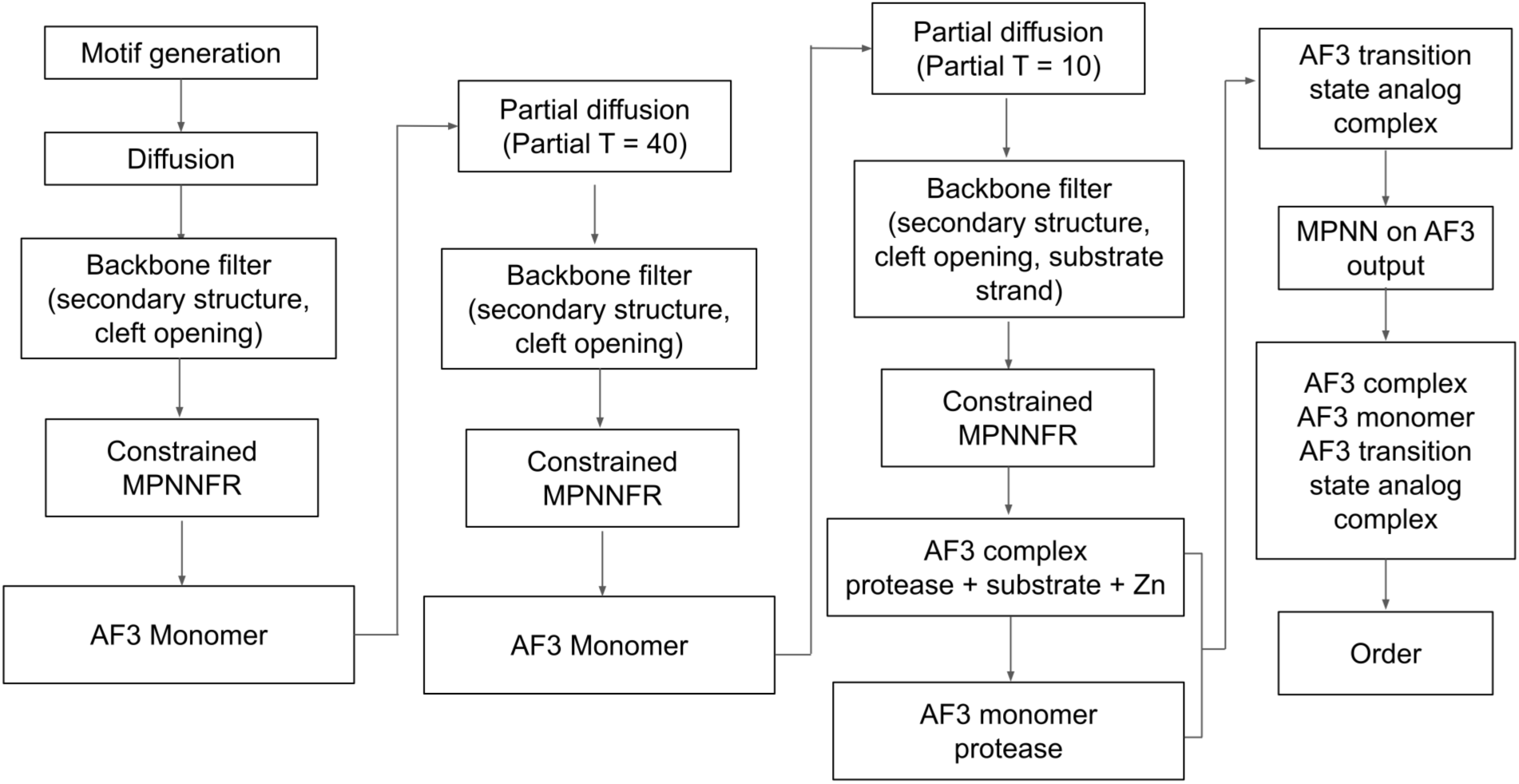
Detailed computational pipeline for the de novo design of metalloproteases. The generation process involves one round of initial motif-scaffolded backbone generation, followed by sequence design and AF3 filtering, two rounds of partial diffusion refinement, followed by sequence design and AF3 filtering and one round of iteration between sequence design and AF3 filtering. See Supplementary Methods for all details of the computational pipeline.

**Extended Data Figure 2.**
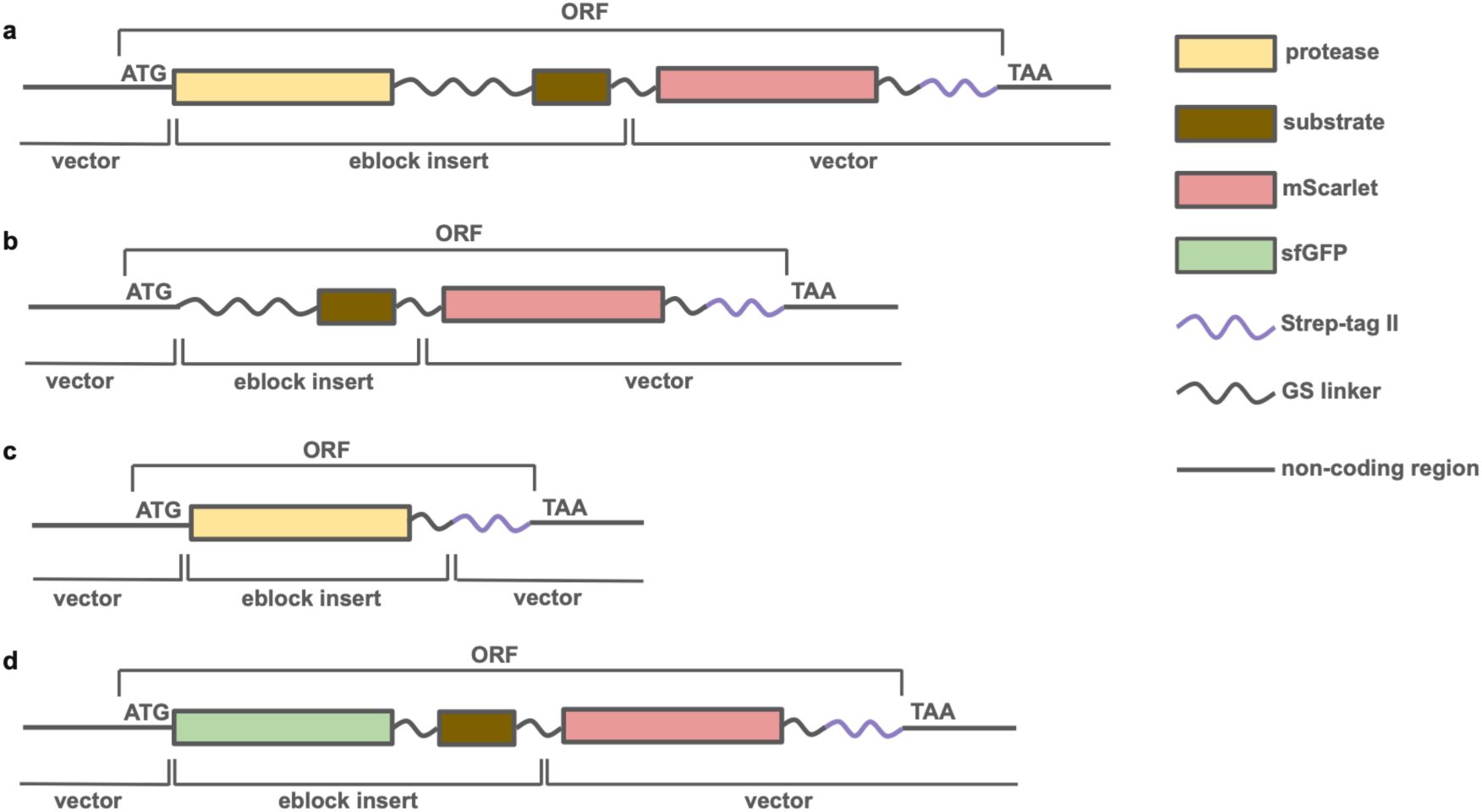
Gene maps of the open reading frames (ORF) of four constructs used in the experimental sections. **a,** The cis construct used for the screen of functional active sites. **b,** The substrate chain of the trans constructs used for the screen of active proteases. **c,** The protease chain of the trans constructs used for the screen of active proteases. **d,** Substrate construct with an N terminal sfGFP and a C terminal mScarlet used in the validation and characterization of active designs.

**Extended Data Figure 3.**
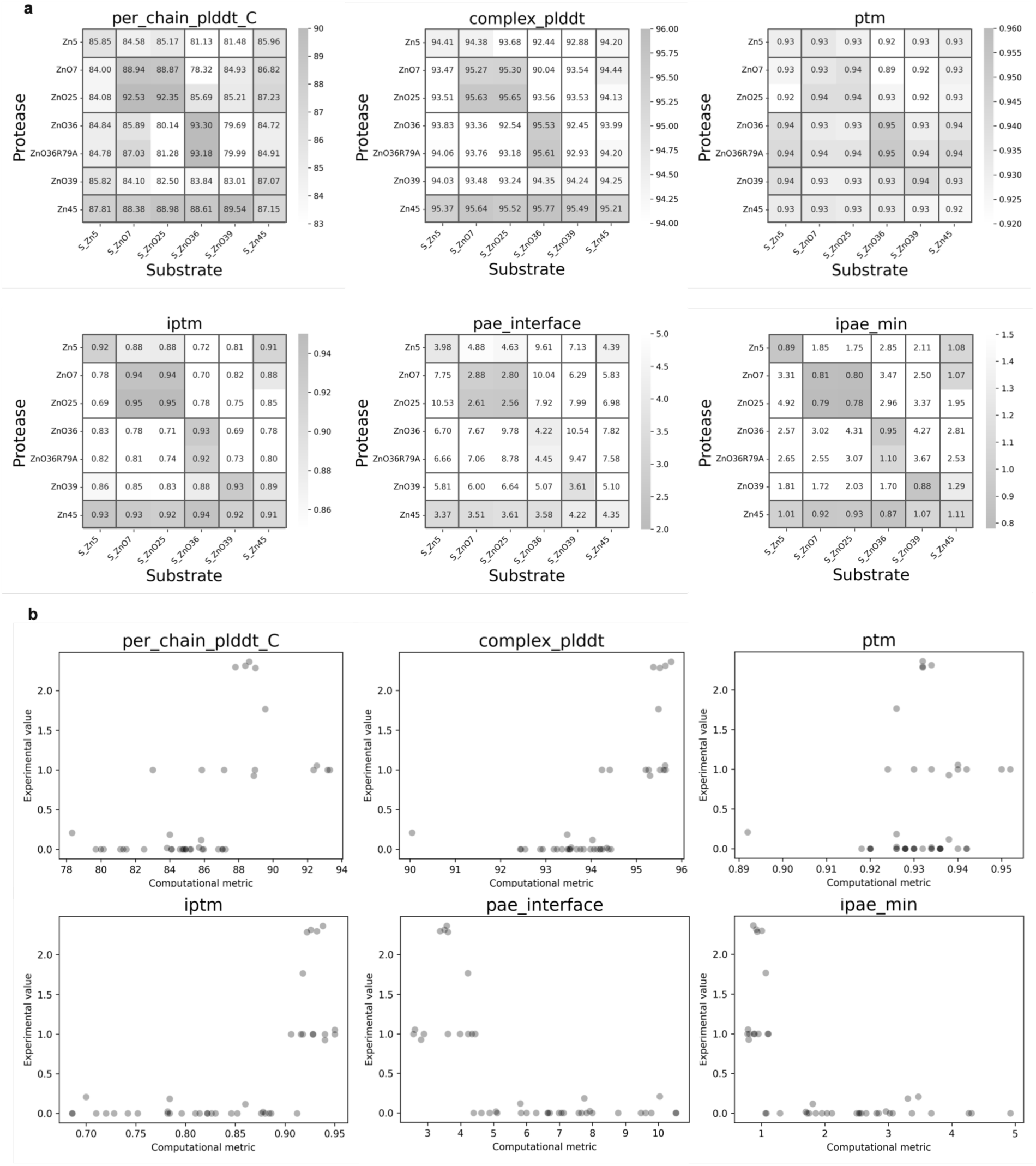
Computational metrics predicative of cross-reactivity. **a,** The heatmap pattern for experimental specificity shown in Fig. 4a, where the diagonal elements and the elements in the last row are highlighted, is observed with multiple computational metrics from AF3 predictions of the [Zn+2].O ES complex, plotted in the same heatmap format as Fig. 4a. **b,** Correlation between experimentally measured normalized digested fraction and computational metrics. High chain C (Zn(II) ion) pLDDT, high complex pLDDT, high iptm, low mean interface PAE and low minimum interface PAE all are predicative of high cross reactivity.

**Extended Data Figure 4.**
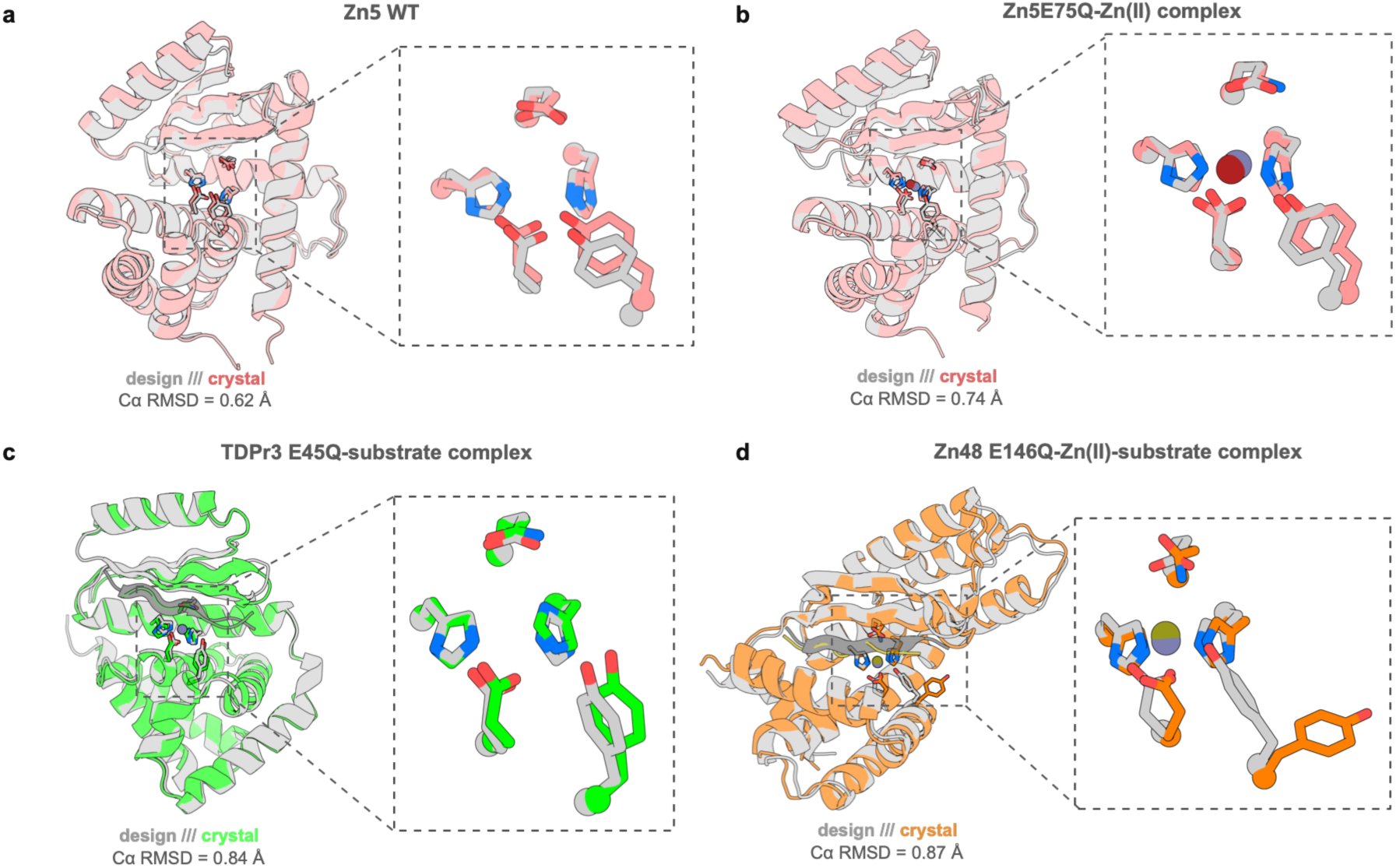
Additional crystal structures for metalloprotease designs. **a,** Apo crystal structure of Zn5 (red) superimposed with the design model (gray) and zoomed in view of the active site. **b,** Crystal structure of Zn5 E75Q-Zn(II) complex (red) superimposed with the design model (gray) and zoomed in view of the active site. **c,** Crystal structure of apo TDPr3 E45Q (green) superimposed with the design model (gray) and zoomed in view of the active site. **d,** Crystal structure of Zn48 E75Q-Zn(II) complex (red) superimposed with the design model (gray) and zoomed in view of the active site.

**Extended Data Figure 5.**
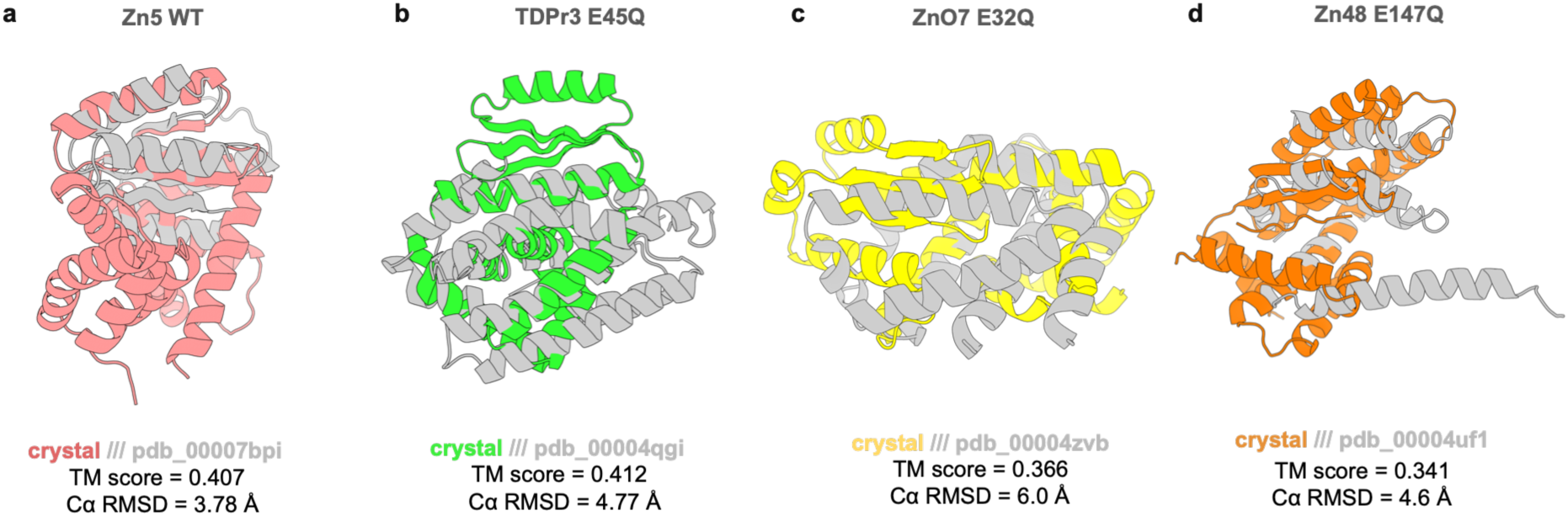
Structural similarity of metalloprotease designs with best matches found in pdb. Searches were performed with: **a**, the Zn5-Zn(II) crystal structure. **b,** the TDPr3 E45Q-Zn(II) crystal structure. **c,** the ZnO7 E32Q-Zn(II)-substrate crystal structure. **d,** the Zn48 E117Q-Zn(II)-substrate crystal structure.

**Extended Data Figure 6.**
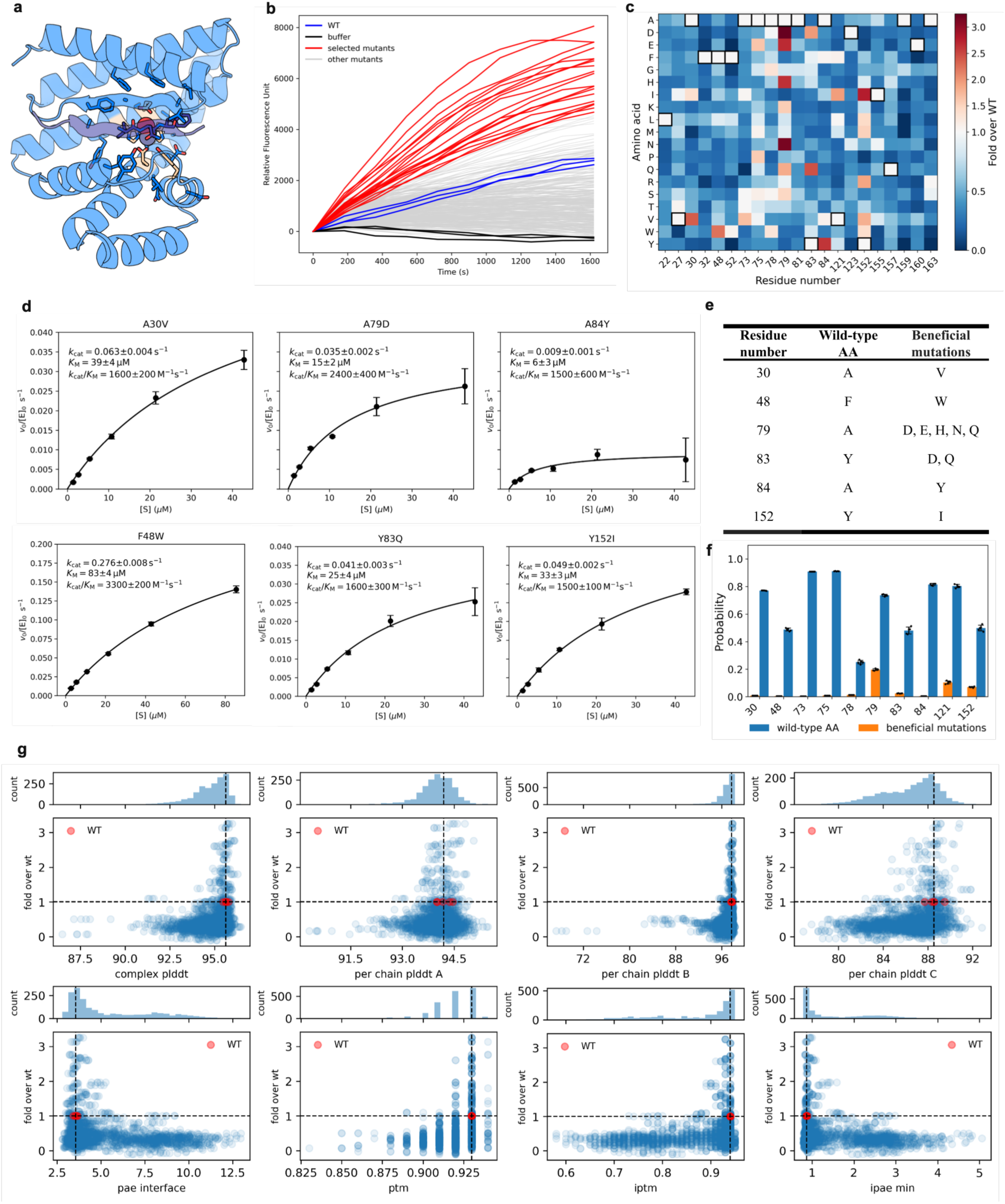
Screening of the active site single site saturation mutagenesis library. **a,** Design model of Zn45 and the substrate of S_ZnO36 showing 21 residue positions chosen for single site saturation in sticks in blue. The five catalytic residues are shown in wheat. **b,** Reaction progress curves for the 378 single variants with WT (blue) and buffer (black) controls. All variants were purified by strep-tag affinity chromatography and concentrations were normalized. Variants in red were further purified using HPLC, tested again to confirm improvements before mutations were recombined (Supplementary Methods 3.19). **c,** Activity heatmap of all single variants **d,** Michaelis-Menten kinetics measurements for six improved single variants, each being the most active variant at the corresponding residue position in the activity screen. *v*_0_: reaction rate in the initial linear region (nM/s). [E_0_]: enzyme concentration, 500 nM. Measurements were made in triplicate. Black dots and error bars represent mean ± std. **e,** Single mutations selected for recombination. **f,** MPNN probabilities for residues where beneficial mutations are discovered. Blue: wild-type AA. Orange: the sum of probabilities for all beneficial substitutions. Individual measurements are represented as black dots and bars and error bars represent mean ± std; n=5. **g,** Correlation between activity fold improvement and AF3 metrics. Many deleterious mutations can be predicted by metrics such as complex pLDDT, ipae min and pae interface.

**Extended Data Figure 7.**
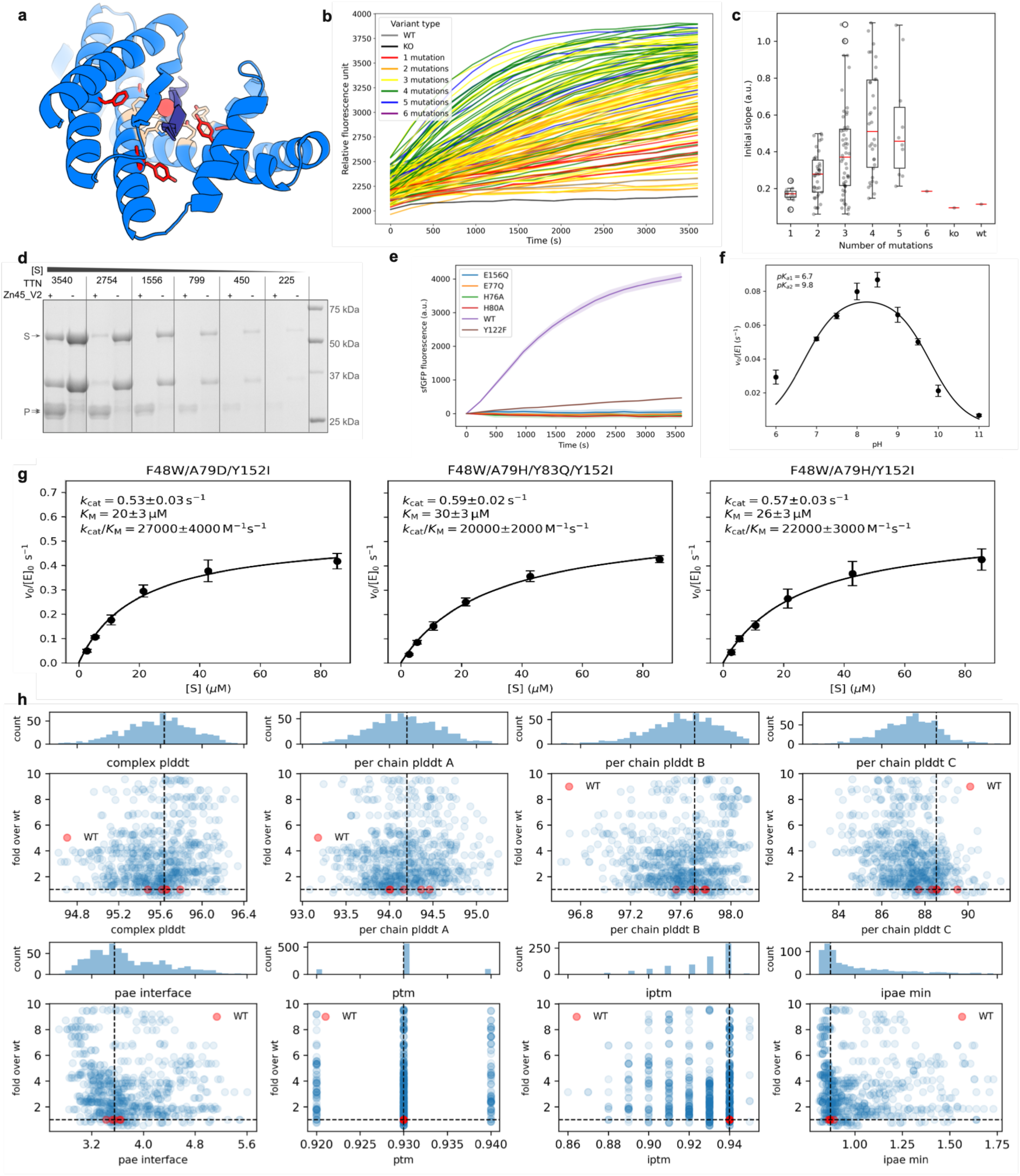
Screening of active site combinatorial library. **a,** Design model of Zn45 and the substrate of S_ZnO36 showing 6 residue positions chosen for combinatorial saturation in sticks in red. The five catalytic residues are shown in wheat. **b,** Reaction progress curves for the 148 combinatorial variants with WT and base KO controls. **c,** The relationship between activity and number of mutations. Box plots show the median (centre line), 25th and 75th percentiles (box limits), and whiskers extending to the most extreme values within 1.5× the interquartile range. Points represent individual observations. The number of variants with mutation numbers 1-6: 11, 38, 52, 34, 10, 1. **d,** Total turnover number (TTN) for Zn45_v2. A series of substrate concentrations of 160 µM, 80 µM, 40 µM, 20 µM, 10 µM, 5 µM, were incubated with or without 20 nM Zn45_v2 at 37 °C for 5 h and the TTN was determined by densitometry. This experiment was performed three times with similar results. Full gel image in Supplementary Fig. S13. **e,** Reaction progress curves of Zn45_v2 and the individual catalytic residue knockout variants. Measurements were made in triplicate. Solid lines and shaded regions represent the mean ± std. **f**, pH dependence of the activity of Zn45_v2 in the low substrate concentration regime (3 µM). Measurements were made in triplicate. Black dots and error bars represent mean ± std. **g,** Michaelis Menten kinetics of three other combinatorial variants. Measurements were made in triplicate. Black dots and error bars represent mean ± std. **h,** Correlation between activity fold improvement and AF3 metrics. Overall the correlation between activity improvement and AF3 metrics is weak.

**Extended Data Figure 8.**
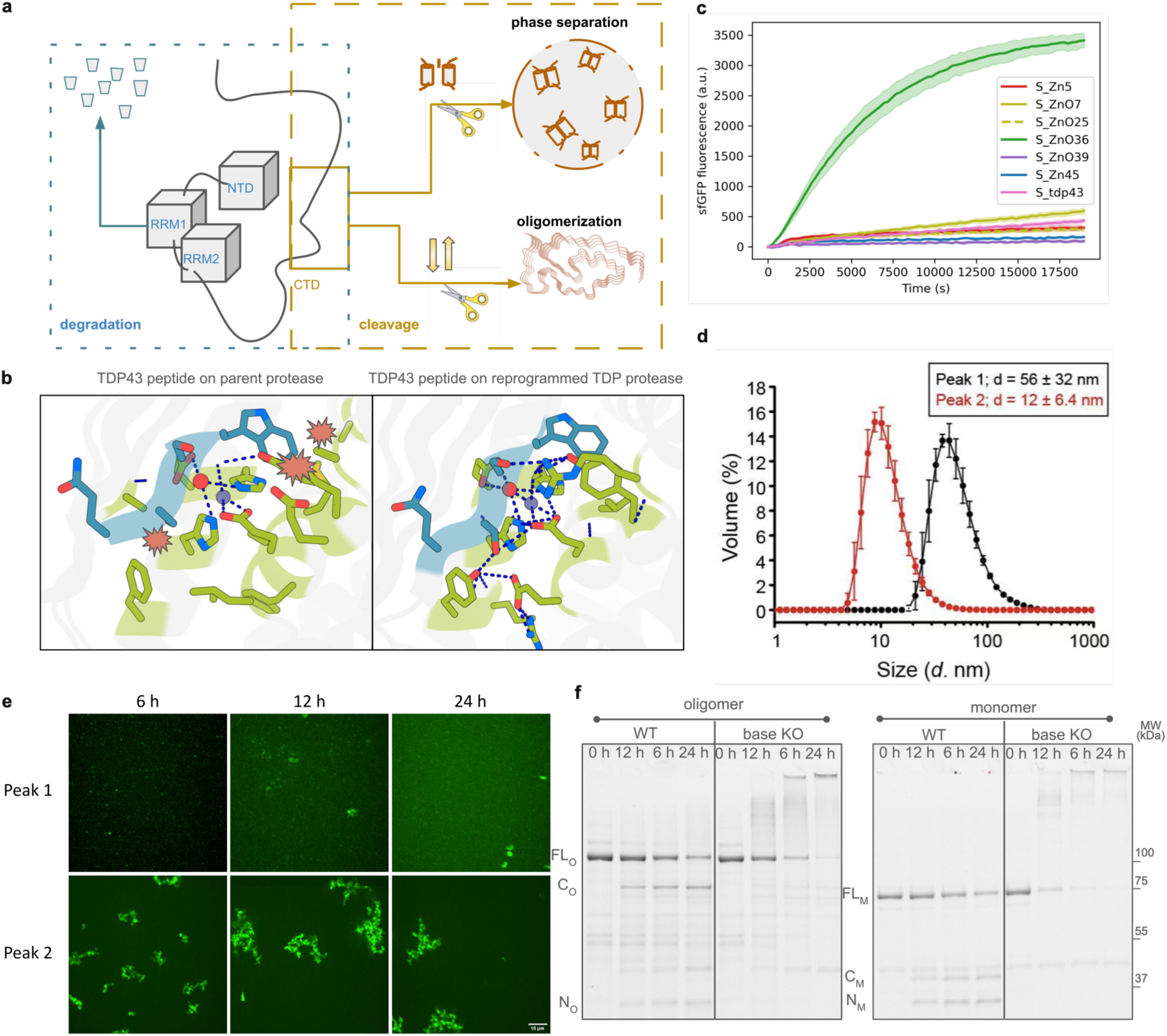
Biological relevance of cleaving human TDP-43 and Characterization of human full length TDP-43 in oligomeric and monomeric formats. **a,** TDP-43 contains structured domains (NTD, RRM1, RRM2) essential for nuclear function and a long disordered C-terminal domain (CTD) that governs liquid–liquid phase separation (LLPS) and oligomerization/aggregation. Full-length degradation risks loss of essential functions, whereas cleavage in the CTD can modulate LLPS and/or oligomerization without ablating function. **b,** Redesigning existing scaffold to target TDP43. **c,** substrate specificity of the TDP43 redesign, TDPr3. Experiments were performed in triplicates. The solid line and the shaded area represent the mean ± std. **d,** dynamic light scattering (DLS) determined the average size of oligomeric TDP-43 (Peak 1) as dozens of molecules solubly clustered together, while monomeric TDP-43 (Peak 2) remains monomeric; Measurements were made in triplicate. Dots and error bars represent mean ± std. **e,** TDP-43 form aggregates after incubation at 37C 6h, 12h, 24 and monomeric TDP-43 at 37C 6h, 12, 24h in buffer 50 mM HEPES, 100 mM NaCl, 50 uM ZnSO4 samples in under microscope, monomers started to aggregate while oligomers remained unchanged. **f,** SDS-PAGE gel analysis after incubating human TDP-43 with TDPn3 or the base knockout (KO). In the WT, the band intensities of the C and N fragments (C_O_, N_O_ for oligomer, C_M_, N_M_ for monomer) gradually increase and the full-length (FL_O_ for oligomer, FL_N_ for monomer) is depleted in time. In base KO, no products are detected. The loss of full-length intensity likely is a result of precipitation as shown by the high molecular weight smears. This experiment was performed three times independently with similar results.

**Extended Data Figure 9.**
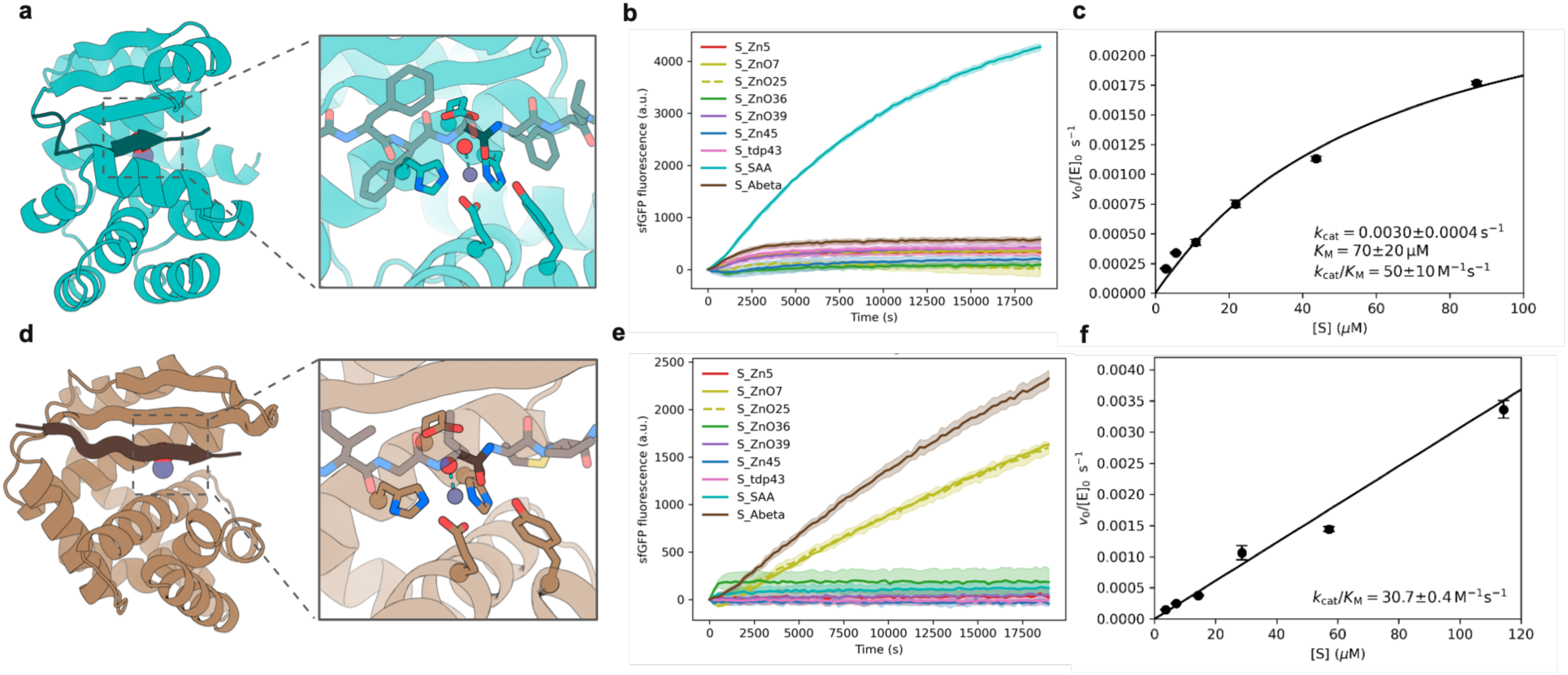
De novo metalloproteases targeting Serum Amyloid A and Amyloid-β peptide. **a,d,** Design model of a de novo zinc protease, SAAc8, against serum amyloid A (SSRSFFSFLG, **a**) and a de novo zinc protease, AbetaF3, against Amyloid-β peptide (KGAIIGLMVG**, d**). **b,e,** Reaction progress curves of SAAc8 and AbetaF3 on their cognate substrates, and on other substrates presented in this work. Measurements were made in triplicate. Solid lines and shaded regions represent the mean ± std. **C,** Kinetics of SAAc8 and AbetaF3 each on their cognate substrates. *v*_0_: reaction rate in the initial linear region (nM/s). [E_0_]: enzyme concentration, 500 nM. Measurements were made in triplicate. Black dots and error bars represent mean ± std.

