## Supplementary Information for "Computational design of metalloproteases"

Anqi Chen^1,2,13^, Kejia Wu^1,2,13*^, Hojae Choi^1,2,3^, Preetham Venkatesh^1,2,3^, Samuel J. Pellock^1,2^, [Nikita Hanikel](https://my.ipd.uw.edu/users/2360)^1,2^, Hojeong shin^1,2^, Declan Evans^1,2^, Kieran Didi^2,4,5^, Lars L. Schaaf^6,7^, Cristina Díaz-Perlas^8^, Brian Coventry^1,2,9^, Donghyo Kim^1,2^, Seth M. Woodbury^1,2,3^, [Pengfei Ji](https://my.ipd.uw.edu/users/2703)^1,2,10^, Shingo Honda^1,2^, Asim K. Bera^1,2^, Hannah Nguyen^1,2^, Alex Kang^1,2^, Yanqing Wang^1,2^, [Xinting Li](https://my.ipd.uw.edu/users/1913)^1,2^, [Stacey Gerben](https://my.ipd.uw.edu/users/1849)^1,2^, [Lemuel Chang](https://my.ipd.uw.edu/users/2695)^1,2^, Xiao Yan^11^, Benjamí Oller-Salvia^8^, Anthony A. Hyman^1^, Donald Hilvert^12*^, David Baker^1,2,9*^

^1^Department of Biochemistry, University of Washington, Seattle, WA 98195, USA

^2^Institute for Protein Design, University of Washington, Seattle, WA 98195, USA

^3^Graduate Program in Biological Physics, Structure and Design, University of Washington, Seattle, WA, USA

^4^Department of Computer Science, University of Oxford, Parks Rd, Oxford OX1 3QD, UK

^5^NVIDIA Corp., Santa Clara, USA

^6^Cavendish Laboratory, Department of Physics, University of Cambridge, Cambridge CB3 0HE, UK

^7^Department of Materials, Imperial College London, Exhibition Road, London SW7 2AZ, U.K.

^8^Institut Químic de Sarrià (IQS), Universitat Ramon Llull, Via Augusta 390, Barcelona 08017, Spain

^9^Howard Hughes Medical Institute, University of Washington, Seattle, WA 98195, USA

^10^Department of Chemistry, Zhejiang University, Hangzhou, China

^11^Max Planck Institute of Molecular Cell Biology and Genetics; Dresden, Saxony, 01307, Germany

^12^Laboratory of Organic Chemistry, ETH Zurich, Zurich, Switzerland

^13^These authors contributed equally

### Methods

#### 1. Computational pipeline for de novo metalloproteases

A comprehensive schematic of the computational pipeline for the de novo design of zinc proteases is shown in Fig. S1.

##### 1.1. Generation of theozyme

To create a minimal catalytic motif, we combined the catalytic residue coordinates of aminopeptidase N and the substrate conformation of native astacin. For the catalytic residues, we extracted the heavy atom coordinates of H293, E293, H297, E316, Y377 from the crystal structure of aminopeptidase N in complex with Zn(II) and a phosphinic transition state analog (TSA; PDB code: 4qhp). For the substrate conformation, we extracted the protease and substrate sequences from ref (PDB: 1qji), and used Alphafold 3 (AF3) to predict the enzyme-substrate (ES) structure of astacin in complex with Zn(II) and this substrate sequence. Superposition was performed using the H293 NE2, E294 OE1 OE2, H297 NE2, E316 OE2, Y377 OH, ZN atoms in the aminopeptidase N TSA complex and the H92 NE2, E93 OE1 OE2, H96 NE2, H102 NE2, Y149 OH, ZN atoms in the astacin ES complex. After superposition, the backbone heavy atom coordinates of the P1-P2’ residues in the substrate chain, the backbone heavy atoms coordinates of the C64, W65, S66 residues in the protease chain, as well as the coordinates of the Zn(II) atom in the astacin ES structure are extracted and combined with the aminopeptidase N catalytic residues. This combined motif is used as the input motif for RoseTTAFold Diffusion 2 for Molecular Interfaces (RFD2-MI) to generate de novo metalloproteases.

##### 1.2. RFD2-MI

We used RoseTTAFold Diffusion 2 for Molecular Interfaces (RFD2-MI) to generate backbones to host the theozyme described in 1.1. RFD2-MI is run using a command such as:

/path/to/rfd2mi/run_inference.py --config-name=aa_ppi	inference.ckpt_path=/path/to/modelcheckpoint/ck.pt	contigmap.length=180-220	inference.num_designs=125	inference.ligand='"ZN"'	inference.input_pdb=/path/to/theozyme.pdb	contigmap.contigs=["'15-100,A64-66,10-100,A293-297,25-100,A315-317,25-100,A376-378,15-100,0_3-5,B3-5,2-2'"]	diffuser.T=50	ppi.hotspot_res='"ZN:ZN"'	inference.output_prefix=/path/to/output

Backbones generated by RFD2-MI are filtered based on low loop contents, ideal C_alpha_-C_beta_ distances, no chain breaks, and clefts large enough for easy access of the substrates.

##### 1.3. Iterated EnhancedMPNN-constrained relax sequence design

To generate sequences that can fold into the scaffolds designed by RFD2-MI, we used EnhancedMPNN for sequence generation as well as side chain packing. We reasoned that simultaneously designing the sequences of both the protease chain (chain A) and the substrate chain (chain B) can enable more flexibility in the sequence space, and thereby increase the chance for the resulting sequences to fold into a structure to scaffold the theozyme precisely. Therefore we carried out two-sided sequence design using EnhancedMPNN with a command such as:

/path/to/mpnn/run.py --model_type ligand_mpnn --enhance plddt_16_20240910-b65a33eb --pdb_path /path/to/scaffolds.pdb --out_folder /path/to/pdb --batch_size 1 --number_of_batches 1 --temperature 0.1 --omit_AA C --chains_to_design A,B --pack_side_chains 1

To ensure preservation of the theozyme geometry, we performed constrained relaxation of the structure after sequence design. We iterated this process 10 times with an implementation analogous to Rosetta FastDesign (Maguire 2021). We refer to this sequence design method as EnhancedMPNN-FR in the supplementary materials.

##### 1.4. AF3 modeling for apo protease

To enable the generation of diverse backbones and to ensure the generation of sequences that can precisely fold into the theozyme, in both the backbone generation stage and the sequence design stage, we simultaneously designed the protease chain and the substrate chain. However, in experimental validation and practical applications, the protease must fold into the desired structure in the absence of a Zn(II) ion or the substrate chain. To assess apo protease folding of the protease chains, we use AF3 to predict the structure of the protease chains.

An example command to run AF3 for the ES complex in shown below:

/path/to/af3/run_alphafold_custom.py --json_path=/path/to/json/input --output_dir=/path/to/output

The JSON input contains only one entity: the protease chain represented as a protein polymer in the format of an amino acid sequence.

From the AF3 models, sequences were selected based on the confidence of prediction with pLDDT> 82 and Cα RMSD < 2 Å.

##### 1.5. Partial diffusion

To ensure precise scaffolding of the catalytic residues and positioning of the target amide bond, we use partial diffusion (Vázquez Torres 2024) to perform finer local sampling of the backbones selected from the previous section. During partial diffusion, the heavy atom coordinates of the five catalytic residues in the protease chain and the heavy atom coordinates of the dipeptide segment containing the target amide bond are fixed. The rest of the backbone atom coordinates are successively noised and denoised using a command such as:

```
/path/to/rfd2mi/run_inference.py --config-name=aa_ppi inference.ckpt_path=/path/to/modelcheckpoint/ck.pt inference.num_designs=25 inference.ligand='ZN' diffuser.partial_T=40
ppi.hotspot_res='"ZN:ZN"' contigmap.contigs=["'18-18,A19-19,60-60,A80-80,97-97,A178-179,2-2,A182-182,28-28,0_4-4,B247-248,2-2'"] contigmap.contig_atoms="\"{'C460':'ZN'}\"" inference.input_pdb=/path/to/backbone.pdb  inference.output_prefix=/path/to/output
```

We used EnhancedMPNN-FR to perform two-sided sequence design of the protease chains and the substrate chains. The protease chain sequences are assessed by AF3 and selected with thresholds of pLDDT > 85 and Cα RMSD < 1.8 Å to ensure apo protease folding into the desired structure. In addition, we reasoned that sidechain preorganization is essential for catalysis (Lauko 2025). Therefore, we further selected scaffolds that satisfy catalytic residue sidechain RMSD < 1.8 Å.

For the scaffolds that pass the above criteria, we performed one more round of partial diffusion (partial T = 10) and EnhancedMPNN-FR for finer sampling.

##### 1.6. AF3 modeling of the protease-Zn(II)-substrate complex

To assess Zn(II) binding, substrate binding, as well as the positioning of the target amide bond with respect to the active site in the nucleophilic attack step, we used AF3 to model the structure of the protease-Zn(II)-substrate complex, which is referred to interchangeably as the ES complex in the supplementary materials.

The JSON input for the AF3 prediction of ES complex contains three entities: the protease chain represented as a protein polymer in the format of an amino acid sequence, the substrate chain represented as a protein polymer in the format of an amino acid sequence, and the Zn(II) atom represented as a ligand in the format of a SMILES string. The Zn(II) is either modeled as an ion alone using the SMILES string ”[Zn+2]”, or modeled in complex with a water oxygen atom using the SMILES string [Zn+2].O.

##### 1.7. AF3 modeling of the protease-Zn(II)-transition state analog (TSA) complex

To assess Zn(II) binding, transition state binding, as well as the stabilization of the transition state by key interactions with the protease catalytic residues, we used AF3 to model the structure of the protease-Zn(II)-transition state analog complex, which is referred to interchangeably as the TSA complex in the supplementary materials.

The JSON input for the AF3 prediction of TSA complex contains three entities: the protease chain represented as a protein polymer in the format of an amino acid sequence, the substrate chain represented as a protein polymer in the format of an amino acid sequence with a phosphoaminate modification in the target dipeptide segment, and the Zn(II) atom represented as a ligand in the format of a SMILES string ”[Zn+2]”.

##### 1.8. Filtering

Designs were selected based on AF3 models of three states: the apo protease state, the ES complex state and the TSA complex state. With the apo protease models, we filtered for AF3 confidence metric pLDDT > 86, catalytic residue sidechain RMSD < 1.5 Å and all-residue Cα RMSD < 1.8 Å. With the ES complex models, we filtered for prediction confidence pLDDT > 92, interface quality metric mean PAE interface (pae interface) < 5, minimum PAE interface (ipae min) < 1.2 catalytic residue sidechain RMSD < 1.5 Å and all-residue Cα RMSD < 1.8 Å. With the TSA complex models, we filtered for prediction confidence pLDDT > 92, interface quality metric mean PAE interface < 5, catalytic residue sidechain RMSD < 1.5 Å and all-residue Cα RMSD < 1.8 Å.

In addition, we requested the average distance between Zn(II) and the three chelating atoms (NE2 of the histidines, the closer of the glutamate OE1 and OE2) to be less than 0.5 Å different from the Aminopepidase N crystal structure (PDB: 4qhp). We also requested the distance between Zn(II) and the target amide carbonyl carbon to be less than 0.5 Å different from the AF3 prediction of the ES structure of astacin.

To further refine designs, the AF3 models of the TSA complexes were fed back to EnhancedMPNN for one last round of sequence design and AF3 prediction in all three states. Designs that passed the above filters both before and after the last round of EnhancedMPNN-AF3 iteration were selected and ordered.

#### 2. Computational redesign of de novo metalloprotease to target TDP-43

##### 2.1. Identify starting docks of de novo metalloprotease and substrate window

To create matching starting points for de novo metalloproteases reprogramming, we performed the “threading” step as previously described in Logos pipeline (Wu 2025). Six proteases in Fig. 3 were given as templates, TDP-43 CTD residue 320-340 was given as target (due to biological relevance of regulating liquid-liquid phase separation). Hard clashes were filtered out. Zinc coordinate was manually aligned back to the passing docks followed by Fastrelax. Highest ranked docks were given to AF3 and constructs with metrics ranging from 1.0<minPAE<1.5, 4.0<meanPAE<5.5, complex pLDDT>92 were selected as initial docks. Range was determined with the hypothesis that these initial docks should have close-to-good AF3 complex metrics, but not necessarily passing to avoid non-specific reprogramming. Initial docks (i.e., coordinates) out of Logos threading were passed to next steps instead of AF3 predictions.

##### 2.2. Partial diffusion with fixed catalytic motif and polar interface motif

To ensure precise scaffolding of the catalytic residues and positioning of the new substrate sequence, we use partial diffusion to perform finer local sampling of the backbones selected from the previous section. During partial diffusion, the heavy atom coordinates of the five catalytic residues in the protease chain, the heavy atom coordinates of the dipeptide segment containing the target amide bond, and neighboring residues on proteases around the catalytic sites making hydrogen bonds to the new substrate are fixed. The rest of the backbone atom coordinates are successively noised and denoised using a command such as:

```
/path/to/rfd2mi/run_inference.py --config-name=aa_ppi inference.ckpt_path=/path/to/modelcheckpoint/ck.pt inference.num_designs=25 inference.ligand='ZN' diffuser.partial_T=20
ppi.hotspot_res='"ZN:ZN"' contigmap.contigs=["'18-18,A19-19,60-60,A80-80,97-97,A178-179,2-2,A182-182,28-28,0_4-4,B247-248,2-2'"] contigmap.contig_atoms="\"{'C460':'ZN'}\"" inference.input_pdb=/path/to/backbone.pdb  inference.output_prefix=/path/to/output
```

We used EnhancedMPNN-FR to perform two-sided sequence design of the protease chains and the substrate chains. The protease chain sequences are assessed by AF3 and selected with thresholds of pLDDT > 85 and Cα RMSD < 1.8 Å to ensure apo protease folding into the desired structure. In addition, we reasoned that sidechain preorganization is essential for catalysis. Therefore, we further selected scaffolds that satisfy catalytic residue sidechain RMSD < 1.8 Å.

We used a range of diffuser.partial_T=8,12,15,20.

##### 2.3. Prediction and filtering

Designs were selected using the same criterion as described above in the de novo metalloprotease section.

#### 3. Computational design of de novo metalloprotease to target TDP-43

##### 3.1. Motif scaffolding

The same theozyme from the two-sided design campaigns are used as the input of the TDP-43 campaign. The only difference is that the substrate sequence is predetermined in the input PDB whereas the side chain rotamers are allowed to freely diffuse. The only atoms that are fixed in the substrate chain are the back bone atoms of the P1 and P1’ residues. Diffusion backbones are generated using a command such as:

/rfd3_path/inference.py out_dir=/path/to/output

ckpt_path=/path/to/rfd3_latest.ckpt inputs=/input/json.json diffusion_batch_size=4 n_batches=5 skip_existing=True cleanup_virtual_atoms=True print_config=True dump_trajectories=False inference_sampler.s_jitter_origin=1 seed=5475644

##### 3.2. Sequence design, prediction and iterations

Diffusion backbones go through the same procedure of backbone filtering, EnhancedMPNN sequence design, AF3 monomer prediction as the two-sided designs. After the first round of AF3 monomer, designs with complex_plddt scores > 82 and Cα RMSD < 1.2 Å and catalytic side chain RMSD < 1.5 Å were recycled back to partial diffusion and MPNN. The designs were then subject to a similar iterated refinement workflow as the two-sided design campaign with different partial diffusion conditions and filtering thresholds.

For the first iteration, a range of partial diffusion temperatures from 11-16 are used for backbone diversification. At the end of this round, the AF3 complex predictions with the Zn(II)-bound water were assessed, and backbones passing mean pae < 5, ipae_min < 1.5 enzyme_plddt > 90, complex_plddt > 90, catalytic residue side chain RMSD < 1.8 are assessed by AF3 monomer. The designs that further pass monomer plddt > 83 are fed back in the second iteration.

For the second iteration, a range of partial diffusion temperatures from 3-7 are used for backbone refinement. At the end of this round, the AF3 complex predictions with the Zn(II)-bound water were assessed, and backbones passing mean pae < 5, ipae_min < 1.2 enzyme_plddt > 90, complex_plddt > 90, catalytic residue side chain RMSD < 1.5 are assessed by AF3 monomer. The designs that further pass monomer plddt > 85 are fed back to the last round of interation

For the last round of iteration, we did not further partially diffuse the designs. Instead, we used the complex AF3 predictions of the input of MPNN and repeated only sequence design. After MPNN, the sequences were predicted in the monomer state, canonical complex state, and the phospho TSA complex state. For the complex state, designs were requested to pass mean pae < 5, ipae_min < 0.9 enzyme_plddt > 92, complex_plddt > 92, catalytic residue side chain RMSD < 1.5 Å. We also requested the distance between Zn(II) and the target amide carbonyl oxygen to be less than 3.2 Å. For the monomer state, designs were requested to pass monomer_plddt > 85 and monomer-complex Cα RMSD < 1 Å. For the phosphoTS state, designs were requested to pass mean pae < 5, ipae_min < 0.9 enzyme_plddt > 92, complex_plddt > 92, catalytic residue side chain RMSD < 1.3 Å.

The design of metalloproteases targeting the human serum amyloid A sequence,	SSRSFFSFLG , and the Amyloid beta peptide sequence, in Serum Amyloid A, and KGAIIGLMVG, followed the same computational pipeline as for the TDP43-targeting metalloprotease, except that the input sequence were specified differently in each case.

#### 4. Experimental materials and methods

##### 4.1. Genetic constructs

The amino acid sequences of all designed proteases, substrates or fusion constructs were reverse translated and codon optimized for expression in E.Coli. Overhang sequences “ATACTACGGTCTCAAGGA” at the 5’ end and “GGTTCCCGAGACCGTAATGC ” at the 3’ end are added for Golden Gate Assembly. Synthetic linear DNA fragments are obtained as eBlocks™ Gene Fragments from Integrated DNA Technologies (IDT). The gene fragments are cloned into a custom vector (LM1369, C terminal Strep-tag II, WSHPQFEK), or a modified version of this custom vector (C terminal mScarlet-Strep-tag II) following an established protocol (Wicky 2022).

Four different genetic constructs were used throughout the experimental sections (Extended Data Fig.2):

1. A genetic fusion of the protease, its designed peptide substrate, an mScarlet and a C terminal Strep-tag II. We refer to this fusion as the “cis construct”.
2. A genetic fusion of 15xGS, a designed peptide substrate, an mScarlet and a C terminal Strep-tag. We refer to this as the “substrate chain of the trans construct”.
3. The designed proteases with a C terminal Strep-tag. We refer to this as the “protease chain of the trans construct”, or the “protease chain” in short.
4. A genetic fusion of a sfGFP, a designed peptide substrate, an mScarlet and a C terminal Strep-tag. We refer to this fusion as the “sfGFP-mScarlet substrate”

##### 4.2. Small scale protein expression and purification

To clone the eblock DNA fragments into expression vectors, Golden Gate Assembly (GGA) reaction mixes are prepared using an Echo 525 Acoustic Liquid Handler (Beckman Coulter Life Sciences) with the following components in a 1μlreaction: 1.2 Units of BsaI-HFv2 (New England Biolabs, R3733L), 40 Units of T4 DNA ligase (NEB, M0202L), 4 fmol vector and 8 fmol linear DNA fragment. The cloning mixture was incubated at 37 °C for 20 mins and transformed into BL21 DE3 cells (NEB, C2527I). For transformation, cells are incubated with the GGA mixture on ice for 30 minutes (min), heat-shocked for 15 seconds (s) and recovered on ice for 5 min. After adding 100μlSOC outgrowth medium (NEB, B9020), cells were recovered for 1 hour (h) at 37 °C on a microplate shaker (1050 rpm) before transferring into LB medium for glycerol stock preparation or into autoinduction medium (AIM) for protein expression.

For protein expression, an AIM is prepared by adding 2 mL of 1 M MgSO_4_(Sigma Aldrich, 230391), 2 mL 50 mg/ml kanamycin and 20 mL of 50X 5052 solution (25% glycerol, Sigma Aldrich, G5516; 2.5% D-(+) glucose, Sigma Aldrich, G7021; 10% α-lactose, Sigma Aldrich, L2643) into 1 L TB II medium (3046052, MP Biomedicals). Protein expression for screening is performed in 96 deep well plates (Corning Axygen, P-DW-20-C-S) with round bottoms. Each well is filled with 1 mL AIM and inoculated with 20μl recovered SOC from cloning. The plates are sealed with Breathe-EASIER microplate sealing film (Diversified Biotech, BERM-2000). Cells were incubated on a microplate shaker (1050 rpm, 37 °C) for at least 20 h before lysis and protein purification.

To lyse the cells, overnight cultures are centrifuged at 4000 g for 5 mins and the supernatants were discarded. To lyse the cell pellet, a lysis buffer was prepared by mixing the following components: 50 mL B-PER™ Complete Bacterial Protein Extraction Reagent (Thermo Fisher, 89823), 1 tablet of Pierce™ Protease Inhibitor Tablets, EDTA-free (Thermo Fisher, A32965), 500μl 100 mM PMSF (Roche, 74348021) stock solution in Isopropyl alcohol (VWR, BDH1133-4LP), 5μl100 mg/ml DNase I (Sigma Aldrich, DN25), 5μl 100 mg/ml Lysozyme (Sigma Aldrich, L6876). For each cell pellet, 100μllysis buffer was added and the pellets were agitated on a microplate shaker (1050 rpm, 37 °C) for 10 mins to ensure complete lysis. Lysates were then centrifuged at 4000 g for 15 mins.

Strep-tag affinity chromatography is performed following a published protocol (Schmidt 2007). In a 96 well 25 μm long drip fritted plate (Agilent, 200953-100), 50μlof 50% Strep-Tactin®XT 4Flow® high capacity resin (IBA Lifesciences, 2-5030-025) slurry is dispensed to each well and washed twice with 400μlwash buffer (100 mM Tris, 150 mM NaCl, pH 8). The bottom of the fritted plate was closed with parafilm and the 400μllysates were applied to each well. The fritted plate was sealed with Aluminum Sealing Film (Corning Axygen, PCR-AS-600) and incubated on a microplate shaker (37 °C, 1050 rpm) for 10 mins to allow sufficient binding of the Strep-tagged proteins to the resin. The parafilm seal and aluminum seal were then removed. The strep resin was washed four times with 400 μL wash buffer (100 mM Tris, 150 mM NaCl, pH 8) and eluted through a 0.22 μm filter plate (Agilent 203940-100)with an elution buffer (50 mM biotin, 100 mM Tris, 150 mM NaCl, pH 8).

For the purification of the protein fraction at the desired molecular weight, size-exclusion chromatography was performed using an HPLC (Agilent 1260 Infinity II LC System) installed with a Superdex 75 Increase 5/150 GL column ( 29148722, Cytiva). The column was equilibrated with a running buffer of 50 mM Hepes, 40 mM NaCl, pH 8. For each sample, 100 ul strep elution was injected and the column was run at a flow rate of 0.65 ml/min. Fractions in 200 μL are collected based on A280 peaks into a 384 deep well plate (Greiner, 781270).

##### 4.3. Large scale protein expression and purification

For the preparation of high concentration sfGFP-mScarlet substrates for kinetics measurements, we perform large scale protein expression and purification using 1L cultures. For protein expression, 7 mL overnight culture is used to inoculate 1 L LB medium (1 w/v% Gibco™ Bacto™ Tryptone

, Thermofisher 211701 ; 1 w/v% NaCl, Thermofisher BP358-10; 1 w/v% Yeast Extract 288610). After 1 h outgrowth at 37 °C on a shaker (250 rpm), 1 mM IPTG is added and the culture is grown at 37 °C for 16-18 hours. Cells were pelleted by centrifugation at 4000 g for 5 mins. The pellets were resuspended in 35 mL of wash buffer (100 mM Tris, 300 mM NaCl, 1 mM PMSF, supplemented with 1 tablet of EDTA free protease inhibitor). Sonication is used to lyse cells with a program of 5 s on, 5 s off, 20 mins. Lysates were cleared by centrifugation at 14,000 g for 20 mins. The cleared lysates were loaded on a gravity flow column loaded with 2 mL strep resin. The resin was washed 3 times with 15 mL wash buffer and eluted into 15 mL of elution buffer (wash buffer with 50 mM Biotin). The elution was concentrated using an Amicon® Ultra Centrifugal Filter, 3 kDa MWCO (Sigma, UFC8003) and further purified by SEC.

Size exclusion chromatography purification was performed on an ÄKTApure Fast Protein Liquid Chromatography (FPLC) instrument (Cytiva) installed with a Superdex 75 Increase 10/300 GL column ( 29148721, Cytiva). The column was equilibrated with a running buffer of 50 mM Hepes, 40 mM NaCl, pH 8. For each sample, 1 mL strep elution was injected and the column was run at a flow rate of 0.8 ml/min. Fractions in mL are collected based on A280 peaks.

##### 4.4. Screening of functional active sites using the cis construct

To screen for designs with functional active sites, we use the cis construct (Extended Data Fig.2a). Following purification, 9 μL of the fusion proteins were distributed into 96 well PCR plates (Corning Axygen, PCR-96-FLT-C). Two identical 96 well PCR plates are prepared for the “+Zn” groups and the “-Zn” groups to identify zinc-dependent cleavage. Zinc sulfate heptahydrate (Sigma Aldrich, 221376) was prepared as a 500 μM stock solution in milliQ water. To the “+Zn” plate, 1 μL zinc sulfate stock solution was added to each well and mixed by pipetting (final Zinc sulfate concentration was 50 μM) ; To the “-Zn” plate, 1 μL milliQ water was added to each well and mixed by pipetting. The PCR plates were sealed tightly with aluminum foil and incubated at 37 °C until analysis by gel electrophoresis analysis. Though the proteases were not normalized in this screen, we estimated based on the A280 absorption values detected during SEC that the protease concentrations were < 30 μM. Thus the amount of zinc sulfate supplemented in the screen should be sufficient for the detection of functional active sites.

For the detection of protein cleavage by SDS-PAGE, 10 μL of 2x Laemmli Sample Buffer (Biorad, #1610375) with 2 mM TECP were distributed into each well of two 96 well PCR plates, one for the “+Zn” groups and the other for the “-Zn” groups. The reaction mixtures were added to the sample loading buffer, mixed by pipetting, boiled at 95 °C for 3 mins and cooled down to 4 °C. Denatured samples were loaded to AnykD™ Criterion™ TGX™ Precast Midi Protein Gel (Biorad, 5678125) and run at 200 V for 35 mins in a Criterion™ Cell (Biorad, 1656001) filled with 1x Tris/Glycine/SDS buffer (Biorad, 1610772). Precision Plus Protein™ Unstained Standards (5 μl; Biorad, 1610363) were loaded in the first or last wells as a reference.

After electrophorese, stain-free gels are activated and imaged by a ChemiDoc XRS+ System (Biorad). Gel images were quantified using ImageJ (NIH) with the Gel Analyzer plugin. All gel images taken in this screen are shown in Fig. S3. Gel lane arrangement is shown in Fig. S4.

For each design the digested fraction $F_{D}$ in the cis screen is determined as follows (Fig. S4):

$$F_{D}^{+} = \frac{I_{p1}^{+} + I_{p2}^{+}}{I_{p1}^{+} + I_{p2}^{+} + I_{s}^{+}}$$

Where $F_{D}^{+}$ is the digested fraction of the cis construct in the +Zn lanes, $I_{p1}^{+}$is the total intensity of the product band corresponding to the protease fragment (~20-25 kDa, band 4 in Fig. S4a) in the +Zn lanes, $I_{p2}^{+}$is the total intensity of the product band corresponding to the mScarlet fragment (~27 kDa, band 3 in Fig. S4a) in the +Zn lanes, and $I_{s}^{+}$is the total intensity of the band corresponding to the full cis construct (~ 55 kDa, band 1 in Fig. S4) in the +Zn lanes. Whenever an mScarlet is included in the experimental construct, we routinely observe a band at ~ 19 kDa (band 5 in Fig. S4a) and a complementary band with a size of construct full length - 19 kDa (band 2 in Fig. S4a). These two fragments are likely the byproducts of mScarlet chromophore maturation (Wei 2015). Because they appeared consistently across all designs in both the +Zn and -Zn groups, we did not take them into consideration for quantification of the cleavage products.

For designs that do not exhibit obvious cleavage in the -Zn control groups, the reported digested fraction $F_{D}=F_{D}^{+}$. For designs that exhibit obvious cleavage in both the +Zn and -Zn groups, the cleavage in the -Zn groups was likely due to the activity of trace amounts of native host proteases co-eluted during purification. To calculate the zinc-dependent digestion, which would be cleavage by the designed active sites, we calculated the background digested fractions, $F_{D}^{-}$ in the -Zn control groups as follows:

$$F_{D}^{-} = \frac{I_{p1}^{-} + I_{p2}^{-}}{I_{p1}^{-} + I_{p2}^{-} + I_{s}^{-}}$$

Where $F_{D}^{-}$ is the digested fraction of the cis construct in the -Zn lanes, $I_{p1}^{-}$is the total intensity of the product band corresponding to the protease fragment (~20-25 kDa) in the -Zn lanes, $I_{p2}^{-}$is the total intensity of the product band corresponding to the mScarlet fragment (~27 kDa) in the -Zn lanes, and $I_{s}^{-}$is the total intensity of the band corresponding to the full cis construct (~ 55 kDa) in the -Zn lanes. The final digested fraction, $F_{D}$, was calculated as:

$F_{D}= \frac{F_{D}^{+} {- F}_{D}^{-}}{1 {- F}_{D}^{-}}$

##### 4.5. Screening of active proteases using the trans constructs.

To screen for active proteases, we used the trans constructs, including two separate chains of the protease and the substrate (Extended Data Fig.2b,c). Following SEC purification, 10 μL of the substrate chain and 10 μL of the protease chains were mixed and then distributed into two PCR plates for the +Zn and -Zn groups. To the “+Zn” plate, 1 μL zinc sulfate stock solution was added to each well and mixed by pipetting; To the “-Zn” plate, 1 μL milliQ water was added to each well and mixed by pipetting. The PCR plates were sealed tightly with aluminum foil and incubated at 37 °C until analysis by gel electrophoresis.

Gel electrophoresis and gel imaging and band quantification were performed according to the same workflow. The digested fraction $F_{D}$ in the trans screen were calculated as follows (Fig. S7) :

$$F_{D}^{+} = \frac{I_{p}^{+}}{I_{p}^{+} + I_{s}^{+}}$$

Where $F_{D}^{+}$ is the digested fraction of the substrate construct in the p+s+Zn lanes, $I_{p}^{+}$is the total intensity of the product band corresponding to the product fragment (~27 kDa, band 2 in Fig. S7a) in the p+s+Zn lanes, $I_{s}^{+}$is the total intensity of the substrate band corresponding to the full substrate chain (~30 kDa, band 1 in Fig. S7a) in the p+s+Zn lanes.

For designs that do not exhibit obvious cleavage in the p+s-Zn control groups, the reported digested fraction $F_{D}=F_{D}^{+}$. For designs that exhibit obvious cleavage in both the p+s+Zn and p+s -Zn groups, the cleavage in the -Zn groups was likely due to the activity of trace amounts of native host proteases co-eluted during purification. To calculate the zinc-dependent digestion, which would be cleavage by the designed proteases, we calculated the background digested fractions, $F_{D}^{-}$ in the -Zn control groups as follows:

$$F_{D}^{-} = \frac{I_{p}^{-}}{I_{p}^{-}+ I_{s}^{-}}$$

Where $F_{D}^{-}$ is the digested fraction of the cis construct in the p+s-Zn lanes, $I_{p}^{-}$is the total intensity of the product band corresponding to the protease fragment (~27 kDa) in the -Zn lanes, $I_{s}^{+}$is the total intensity of the substrate band corresponding to the full substrate chain. The final digested fraction, $F_{D}$, was calculated as:

$F_{D}= \frac{F_{D}^{+} {- F}_{D}^{-}}{1 {- F}_{D}^{-}}$

##### 4.6. Functional validation of individual catalytic residues

To investigate the contribution of each designed catalytic residue in the activity of the designs, we used mutagenesis to knock out each designed catalytic residue individually and assessed the impact on the activity using an SDS-PAGE gel cleavage assay. The mScarlet-sfGFP substrate construct (Extended Data Fig.2d) and the separate protease chains (Extended Data Fig.2c) were used for this experiment. Proteases and substrates were purified by Strep-tag affinity chromatography and SEC. Before the experiment, all proteases were normalized to 20 μM and all substrates were normalized to 5 μM. We prepared two separate PCR plates, one for the +Zn group and another for the -Zn group. We mix 5 μL substrate and 5 μL protease in each well. To the “+Zn” plate, 1 μL 500 μM zinc sulfate stock solution was added to each well and mixed by pipetting; To the “-Zn” plate, 1 μL milliQ water was added to each well and mixed by pipetting. The PCR plates were sealed tightly with aluminum foil and incubated at 37 °C until analysis by gel electrophoresis. To account for the different activities of different designs, we take two time points: one at 1 hour and another at 8 hours. Gel electrophoresis, imaging and image analysis were done in the same way as described in section 3.5 for the trans screen. The measurements were carried out in triplicate. The digested fractions at the end of each time point are calculated as in section 3.5.

For designs Zn5, Zn44, Zn45, Zn52, ZnO7, ZnO25, ZnO39, the following residues are mutated: three Zinc chelators (two histidines mutated to alanine; glutamate mutated to glutamine), general base (glutamate, mutated to glutamine), oxyanion stabilizer (tyrosine mutated to phenylalanine). In all measured designs except for Zn44, knocking out one of the designed residues decreased activity. In Zn44, however, the YtoF mutation resulted in a slight increase in protease activity. This is aligned with the AF3 prediction of the TSA complex, where the hydrogen bond between the tyrosine OH atom and the oxyanion in the TSA complex has a non-ideal bond angle of 170°. Whereas in the other four characterized designs, this angle is between 113.9° and 134.7°, closer to the native values of 108.7° in the TSA crystal structure of aminopeptidase N (PDB: 4qhp) and 109.8° in the TSA crystal structure of astacin (PDB: 1qji, Fig. S11).

For design ZnO36, in addition to the five designed catalytic residues, we discovered an arginine residue, R79, adjacent to the transition state oxyanion (Fig. S10). Therefore we tested if this arginine served as an additional oxyanion stabilizer by mutating it into a lysine or an alanine. While the R79K didn’t significantly affect activity, the R79A substitution increased the activity of ZnO36 by 8 times. This result contradicted our hypothesis for the role of R79. However, it demonstrated the potential of de novo proteases to be improved by random mutagenesis and directed evolution.

For simplicity, in the main text and Fig. 3a-c, we only discuss Zn5, ZnO7, ZnO36, ZnO39, Zn45, each representing a unique scaffold.

##### 4.7. Assessment of Zinc binding

The Zn(II) binding of the designed Zn proteases and their single-residue mutants was evaluated using a competition binding assay under deployment of Mag-Fura-2—a ratiometric dye that served as both reporter and competitor. The change of the 323/342 nm absorbance ratio (absorbance at the maximum of the metal-bound state to absorbance at the isosbestic point) was used for the respective analysis. First, the dissociation constant of the dye (*K*_d_(Mag-Fura-2, Zn(II)) = 13 ± 4 nM) was determined in competition with nitrilotriacetic acid (with *K*_d_(NTA, Zn(II)) = 830 pM under the experimental conditions of *I* = 0.05 M, pH 8.0, and 25 °C). Afterwards, the dissociation constants for the proteins were assessed in competition with the ratiometric dye. For each measurement, the proteins, Mag-Fura-2, and/or NTA were each utilized at a 10 µM concentration in a 10 μL buffered solution with varying Zn(II) metal ion concentrations at 0 to 28 µM. The samples were incubated for 3 days in the dark at 25 °C to ensure binding equilibrium.

Additionally, design ZnO39 was subjected to the same competition binding assay under deployment of Fura-2—a ratiometric dye with a lower dissociation constant in respect to Zn(II) under otherwise the same experimental conditions ((*K*_d_(Fura-2, Zn(II)) = 320 ± 60 pM)). Its dissociation constant was determined with help of ethylene glycol-bis(β-aminoethyl ether)-N,N,N′,N′-tetraacetic acid (with *K*_d_(EGTA, Zn(II)) = 58 pM under the experimental conditions of *I* = 0.05 M, pH 8.0, and 25 °C) as competitor. The change of the 340/354 nm absorbance ratio (absorbance at the maximum of the metal-bound state to absorbance at the isosbestic point) was used for the respective analysis.

##### 4.8. Characterization of cross-reactivity

The mScarlet-sfGFP substrate construct (Extended Data Fig.2d) and the separate protease chains (Extended Data Fig.2c) were used for this experiment. Proteases and substrates were purified by Strep-tag affinity chromatography and SEC. Before the experiment, all proteases were normalized to 20 μM and all substrates were normalized to 5 μM. The first histidine to alanine KO was used as the negative control for this experiment. We prepared two separate PCR plates, one for the WT group and another for the KO group. We mix 5 μL substrate and 5 μL protease in each well. To both the WT and the KO plates, 1 μL 500 μM zinc sulfate stock solution was added to each well and mixed by pipetting. The PCR plates were sealed tightly with aluminum foil and incubated at 37 °C until analysis by gel electrophoresis. To account for the different activities of different designs, we take two time points: one at 1 h and another at 8 h. For designs Zn45, ZnO7, ZnO25, ZnO36R79A, ZnO39, the 1 h measurements are used for quantification. For designs Zn5, and ZnO36, the 8 h measurements are used for quantification. Gel electrophoresis, imaging and image analysis were done in the same way as described in section 3.5 for the trans screen. One set of the gel images are shown in Fig. S13. The measurements were carried out in triplicate. The digested fractions at the end of each time point were calculated as in section 3.5.

To calculate the normalized digested fraction, the digested fraction for each protease-substrate pair was divided by the digested fraction of the protease in this pair cleaving its own designed substrate. Values of digested fractions > 1 indicate higher activity of the designed protease on the designed substrate of a different protease than on its own designed substrate.

For simplicity, in the main text and Fig. 4a, we only discuss Zn5, ZnO7, ZnO25, ZnO36, ZnO36R79A, ZnO39, Zn45.

The cross-reactivity shown in Fig. 4a was not completely unexpected. In addition to the structural implications we discussed in the main text (Fig. S15-16), to understand this cross-reactivity, we used AF3 to model the Zn(II)-water ES complex of designs when paired with substrates of other designs. We found multiple computational metrics were predicative of the substrate selectivity patterns shown in Fig. 4a, including ptm, Zn plddt (per chain plddt C), complex plddt, iptm, mean pae interface, and minimum pae interface. The experimental heatmap pattern, where the diagonal elements and the last row are more intense than the rest of the elements, are observed in all of these computational heatmaps (Fig. S17). To quantify the correlation between these computational metrics and the experimentally measured substrate selectivity, we calculate the Pearson’s r between the experimental noralized digested fractions and the computational metrics; Zn plddt (per chain plddt C), complex plddt, iptm, mean pae interface, and minimum pae interface exhibit |r| > 0.58 (Fig. S18), suggesting correlation.

##### 4.9. Kinetics

To acquire continuous reaction measurements in time, we used the sfGFP-mScarlet substrate construct. When this substrate was intact, mScarlet absorbed the fluorescence of sfGFP due to the overlap of the mScarlet excitation and GFP emission spectrums, resulting in Fluorescence Resonance Energy Transfer (FRET) and quenching of sfGFP fluorescence. Upon cleavage of the substrate, the GFP was released, terminating FRET quenching and leading to the increase of fluorescence. This signal can be correlated to the fraction of cleaved substrate over time.

To prepare substrates for kinetics measurements, we prepared 1 L cultures of the sfGFP-mscarlet substrates as described in section 3.3, snap-froze the substrates with liquid nitrogen, and stored in a -80 °C freezer until further use.

To measure the kinetics of ZnO7, a 2x serial dilution of a 40 μM substrate stock was prepared as described in section 3.3. The ZnO7 protease was purified as described in section 3.2, and diluted to 5 μM in Hepes 40 mM, NaCl 50 mM, ZnSO_4_ 25 μM. To equilibrate the substrates to the plate reading conditions, 18 μL serial-diluted substrates were first added to a black 384 well plate (Corning, 3762) and centrifuged to ensure even coverage of the bottom of the plate. To prevent evaporation, the plate was sealed with a TempPlate RT Select Optical Film (USA Scientific, 2921-7800) and loaded into a plate reader (Biotek Synergy Neo2, Agilent) controlled by the Gen 5 software (version 3.14) set to 37 °C and equilibrated for 30 mins. To initiate the reaction, 2 μL of the protease stock or 2 ul of reaction buffer as a control was added to the substrates and mixed by pipetting such that the final protease concentration used was 500 nM. The plate was centrifuged at 1000 rcf for 1 min to remove air bubbles and to ensure even coverage of the bottom of the plate. As the fluorescence signals resulted from the substitution of substrate by product, to calibrate the relative fluorescence readout to actual cleavage of the substrates, we prepared standard curves for both the substrate in 2x serial dilution from 10 μM, and the products (separately expressed sfGFP and mScarlet in equimolar ratio) in 2x serial dilution from 10 μM. Fluorescence intensity measurements were taken with an excitation wavelength of 485 nm and an emission wavelength of 528 nm. The standard curves and the reaction progress curves were taken on the same plate in triplicate (Fig. S19a-f). The linear regions of the reaction progress curves were used to calculate the initial reaction rates. The fitting of Michaelis-Menten kinetics parameters was performed with scipy curve_fit function using non-linear least squares minimization. The conversion between fluorescence intensity and product concentration was performed using the standard curve for the difference between product fluorescence and the substrate fluorescence as a function of concentration (Fig. S19f), after correction for inner filter effect (IFE; Liu 1999, Kubista 1994).

At high substrate concentrations, the emission or excitation light can be absorbed by quenching groups on neighboring substrates or cleaved product molecules so that only a fraction of the fluoresced light impinges upon the detector system of the fluorometer, known as the inner filter effect (IFE; Liu 1999, Kubista 1994). In the high substrate concentration groups of the kinetics measurements, the conversion between the fluorescence readout and the molarity of the cleavage product is not simply linear; thus the fluorescence readout was corrected to the value in the absence of other high concentration absorbing species before this conversion can be performed. The strength of IFE can be described by defining a factor of inner filter effect (*f*_IFE_):

$$f_{IFE}= \frac{F_{fluorophore}}{F_{fluorophore}([S]=0)}$$

In this equation, *F*_fluorophore_ is the fluorescence contribution of the fluorophore in the presence of high concentration substrates (*F*_fluorophore_= *F*_mixture_ - *F*_substrate_ , where *F*_mixture_ is the fluorescence of a mixture of the fluorophore and the substrate, and *F*_substrate_ is the fluorescence of the substrate alone) and *F*_fluorphore_([S]=0) is the fluorescence of the fluorophore in the absence of substrate. As a consequence of substrate absorption, the factor of inner filter effect is a function of [S], *l*, the length of the optical path, and the molar absorption coefficients at the wavelengths of the excitation and emission lights (Palmier 2007). For fixed optical conditions, optical paths and molecular identities, *f*_IFE_ reduces to a function of [S].

To correct for IFE, we prepared the same serial dilutions of the substrates as we used for the measurements of reaction progress curves, supplemented with 3 μM or 0 μM fluorophore sfGFP, measured the fluorescence of both groups, *F*_mixture_([S]), *F*_substrate_([S]) and calculated the difference between the two groups, *F*_sfGFP_([S]), as a function of substrate concentration. The total volume of the correction samples were 20 μl, the same as the kinetics measurements to keep the optical path, $l$, consistent. The optical settings, including the excitation, emission wavelengths and bands were the same as the kinetics measurements to keep extinction coefficients consistent. Thus the *f*_IFE_ was only a function of substrate concentration and was calculated as follows:

$F_{sfGFP}([S])= F_{mixture}([S]) -F_{substrate}([S])$

$$f_{IFE}([S])= \frac{F_{sfGFP}([S])}{F_{sfGFP}([S]=0)}$$

These *f*_IFE_ values (Fig. S19h) for different concentrations were used to correct for the fluorescence readout in the progress curves before using the standard curves to convert to the initial reaction rates.

The kinetics of Zn45 cleaving the designed substrate of ZnO36 was performed in a similar workflow except that higher maximum substrate concentration was used due to the higher K*_M_*. The raw progress curves, standard curves and *f*_IFE_ as a function of concentrations are shown in Fig. S20.

For both kinetics measurements, we confirmed the cleavage detected by the optical readouts by running an end point SDS-PAGE gel after plate reading. The gel images are shown in Fig. S21.

##### 4.10. Screening for TDP-43 proteases

Synthetic DNA was obtained from IDT for 48 redesigns from the ZnO36 scaffold and 113 de novo designs of TDP-43 proteases. Proteins were expressed in 4 ml cultures using BL21 DE3 cells and purified using strep-tag affinity chromatography without concentration normalization.

For the redesigns, screening was first performed using the cis constructs by incubating the purified proteins with 50 μM ZnSO_4_ in 40 mM HEPES 50 mM NaCl. Designs that showed cis cleavage were further tested using the trans constructs by incubating the purified proteins with 50 μM ZnSO_4_ as well as 3 μM substrate in 40 mM HEPES 50 mM NaCl. Cleavage both in the trans and cis screens was detected using SDS-PAGE gel analysis. Screening outcomes are shown in Fig. S17 and Fig. S18.

For the de novo designs, we only performed screening in the trans format. The FRET signal generated by the sfGFP-substrate-mScarlet construct of the TDP-43 target sequence was measured using a plate reader to indicate cleavage. To initiate the reaction, 2 μl of purified proteins were added to 3 μM substrate, 50 μM ZnSO_4_ in 40 mM HEPES 50 mM NaCl. The screening outcome is shown in Figure S19.

##### 4.11. Substrate selectivity of TDPn3 and TDPr3

To test the substrate preference of TDPn3 and TDPr3, the sfGFP-substarte-mScarlet constructs of S_Zn5, S_ZnO7, S_ZnO25, S_ZnO36, S_ZnO39, S_Zn45, S_tdp43 were normalized to 3 μM. For both the protease groups and the buffer only control groups, 18 μl of substrate was added to a 384 well clear bottom plate. The proteases (5 μM for TDPn3 and 60 μM for TDPr3) were mixed with 100 μM ZnSO_4_. To initiate the reaction, 2 μl protease stock or HEPES buffer was added to the 384 well plate and mixed with the substrate. The final enzyme concentration was thus 500 nM for TDPn3 and 6 μM for TDPr3. Each condition is measured in triplicates and the progress curves of the TDPr3 and TPDn3 groups are shown in Fig. 4b,d after subtraction of the buffer only groups.

Screening of the SAA and Aβ proteases were performed in the same manner. Synthetic DNA was obtained from IDT for 34 designs against the KGAIIGLMVG sequence in the Aβ peptide and for 14 designs against the SSRSFFSFLG sequence in the SAA peptide. Only the fastest design was chosen for subsequent selectivity and kinetics characterization, which were performed as described in sections 4.9-4.11.

##### 4.12. Cleavage of full-length human TDP-43

TDP-43 was expressed in Sf9 insect cells (BioTrend, Cat#94-001F) and purified as previously described (PMID: 40412392). This cell line was used as supplied by the commercial vendor and were not independently authenticated in the lab. The lab routinely tests for mycoplasma contamination and the outcome has been negative. Briefly, Sf9 cells expressing MBP-TDP-43-GFP were cultured in 500 mL and harvested for purification. The lysate was applied sequentially to a His-tag affinity column and a Superdex 200 size-exclusion column. Monomeric TDP-43 was obtained by cleaving the MBP tag from TDP-43-GFP using in-house 3C-protease, while oligomeric TDP-43 was purified as MBP-TDP-43-GFP. Protein concentrations were determined using a Nanodrop spectrophotometer at 280 nm.

Purified proteases in trans constructs were prepared as described above. Following SEC purification, 4 μL of the oligomeric TDP-43 and monomeric TDP-43 (normalized to 16 uM in 40mM hepes, 50mM NaCl), 4 μL of the proteases or control of buffer only (normalized to 40 μM in 40 mM HEPES, 50mM NaCl, 50 μM ZnSO_4_) were mixed and then distributed into four groups for different incubation time points. The PCR tubes were sealed tightly with aluminum foil and incubated at 37 °C for 0hr, 6hr, 12hr, 24hr and then immediately transferred to -20 °C. After all incubations are completed, SDS-PAGE analysis was performed to detect the cleavage products

##### 4.13. Intact protein mass spectrometry

To identify the molecular mass of cleaved products, intact mass spectra was obtained via reverse-phase LC/MS on an Agilent G6230B TOF on an AdvanceBio RP-Desalting column, and subsequently deconvoluted by way of Bioconfirm using a total entropy algorithm.

##### 4.14. Protein sample preparation for crystallography

Protein samples for crystallography are expressed and purified as described in Supplementary Method section 3.3.

The Zn5 apo sample was submitted for screening after purification and concentration.

The Zn5-Zn (II) complex of Zn5 was prepared by adding ZnSO_4_ to wild-type Zn5.

The Zn5 E75Q-Zn (II) complex of Zn5 was prepared by adding ZnSO_4_ to a Zn5 E75Q-GSx15-substrate fusion construct.

The Michaelis complex of ZnO7 was prepared by adding ZnSO_4_ and a synthetic peptide to the E32Q variant of ZnO7.

The TDPr3 apo sample was prepared by adding ZnSO_4_ to a E32Q variant of TDPr3.

The TDPr3 E45Q-substrate complex was prepared by adding ZnSO_4_ to a E32Q-15xGS-substrate construct.

The Michaelis complex of Zn48 was expressed as a genetic fusion, where the protease and the substrate were linked with a 15xGS flexible linker, and supplemented with ZnSO_4_.

For the supplementation of Zn(II), after SEC purification, ZnSO₄ was added to the elution at a final molar concentration 1.5× that of the protein. For supplementation with synthetic substrate peptides, the peptide was reconstituted in water and added to the elution at a final molar concentration 1.5× that of the protein. The sample was incubated overnight at 4 C to ensure equilibrium and then concentrated to > 20 mg/ml before the preparation of screening plates.

##### 4.15. X-ray crystallography

All crystallization experiments were conducted using the sitting drop vapor diffusion method.

Crystallization trials were set up in 200 nL drops using the 96-well plate format at 20 ˚C.

Crystallization plates were set up using a Mosquito LCP from SPT Labtech, then imaged using UVEX microscopes and UVEX PS-256 from JAN Scientific. Diffraction quality crystals formed in

21% PEG 350 MME, 0.5 M Magnesium chloride and 0.05 M Tris pH 7.5 for the Zn(II) complex of Zn5; in 0.65 M Imidazole pH 7.0, and 35% v/v Glycerol for the E32Q -Zn(II) - substrate complex of ZnO7 ; in 0.07 M Sodium acetate trihydrate pH 4.6, 5.6% w/v Polyethylene glycol 4,000, and 30% v/v Glycerol for TDPr3 E45Q apo; in 0.05 M Glycine ph 9.0 and 55 % v/v PEG 400 for Zn5 apo; in 60% v/v Tacsimate pH 7.0 for the E146Q-Zn(II)-substrate complex of Zn48; in 0.2 M Ammonium chloride and 40% (v/v) MPD for the E75Q-Zn(II) complex of Zn5

and in 0.3 M Magnesium nitrate hexahydrate, 0.1 M Tris pH 8.0 and 23 % w/v PEG 2000 for the E45Q-substrate complex of TDPr3.

Diffraction data was collected either at the Advanced Photon Source beamline on 24-ID-E (for the Zn(II) complex of Zn5, Zn5 apo, the E75Q-Zn(II) complex of Zn5 and the E45Q-substrate complex of TDPr3or at the National Synchrotron Light Source II [Beamline 17-ID-1 (AMX) or 17-ID-2 (FMX)] (for the E32Q -Zn(II) - substrate complex of ZnO7, TDPr3 E45Q apo, and the E146Q-Zn(II)-substrate complex of Zn48. X-ray intensities and data reduction were evaluated and integrated using XDS (version 20240723/ 20250119, Kabsch 2021) and merged/scaled using Pointless/Aimless in the CCP4 program suite (version 8.0.017, Winn 2011). Structure determination and refinement starting phases were obtained by molecular replacement using Phaser (version 2.8.3, McCoy 2007) using the designed model for the structures. Following molecular replacement, the models were improved using phenix.autobuild (version 2.1/2.2, Adams 2010); efforts were made to reduce model bias by setting rebuild-in-place to false, and using simulated annealing. Structures were refined in Phenix (version 2.1/2.2, Adams 2010). Model building was performed using COOT (version 0.9.8.95, Emsley 2004). The final model was evaluated using MolProbity (version MolProbity v4.02, Williams, 2018). Data collection and refinement statistics are recorded in Table S5. Data deposition, atomic coordinates, and structure factors reported in this paper have been deposited in the Protein Data Bank (PDB), http://www.rcsb.org/ with accession code 11CE, 11CM, 11CP, 11CR, 11CU, 11CY and 11DU.

##### 4.16. Analysis of structural similarity between AF3 models and crystal structures

For the calculations of overall backbone RMSD between crystal structures and AF3 predictions, we used the pymol “align” function with the command:

align af3, crystal

For the calculations of active site all-atom RMSD, we first used the pymol “superimpose” function to align two active sites with the command:

super af3_active_site, crystal_active_site

The all-atom RMSD of the active site was the calculated using the command:

rms_cur af3_active_site, crystal_active_site

##### 4.17. Analysis of structural similarity between crystal structures and existing protein structures in nature

We used the Foldseek search engine (https://search.foldseek.com/search) to search for fold similarity between the crystal structures of the metalloprotease designs and existing protein structures the Protein Data Bank available to Foldseek as of Feb 16, 2026, including:

The TM-align mode was selected and the refined crystal structures were uploaded for the search of similar scaffolds. The outputs were sorted by TM scores in a descending order and the highest TM scores across all databases are reported in the main text. Cα RMSD reported by Foldseek and the corresponding TM scores with the top pdb hits are shown in Extended Data Fig. 7.

##### 4.18. Analysis of sequence similarity between designs and existing protein in nature

We used NCBI BLASTP (https://blast.ncbi.nlm.nih.gov/Blast.cgi?PAGE=Proteins) to search for sequence similarity between designs and existing proteins available in the Clustered non-redundant database as of Feb 20, 2026. A E-value threshold of 100 was chosen in the search setting. The search results were sorted by E-values in ascending order. The lowest E-values and the corresponding Accession numbers were reported

##### 4.19. Screening of the active site single site saturation mutagenesis library and the combinatorial library

The synthetic DNA of 378 single variants of Zn45 were obtained from Twist Biosciences. Individual protein variants were expressed and purified following the procedure described in section 3.2 but without the last HPLC purification. For the screen of activity, single variants are mixed with the sfGFP-mScarlet substrate of ZnO36 and ZnSO_4_ with final concentrations of 500 nM protease, 3 μM substrate and 5 μM Zn^2+^. The slopes of the linear regions of the progress curves are obtained as the activity of these single variants and compared to the wild-type to calculate the fold improvements. Variants with > 1.5 fold improvements were further purified with HPLC and normalized to confirm their activity at two substrate concentrations, 3 μM and 30 μM (Supplementary Figure 29a). The single mutations that lead to > 3 fold were recombined (Supplementary Figure 29b), and predicted by AF3. Out of 288 combinatorial variants, 148 variants passed AF3 thresholds of complex plddt > 93 and ipae_min < 1.5. For each residue position where beneficial mutations were discovered, kinetics were measured for the best single variant as described in section 3.9 and reported in Extended Data Fig. 8.

The synthetic DNA of these 148 variants were obtained from IDT Biotechnology. Individual protein variants were expressed and purified following the procedure described in section 3.2 but without the last HPLC purification. For the screen of activity, combinatorial variants are mixed with the sfGFP-mScarlet substrate of ZnO36 and ZnSO_4_ with final concentrations of 50 nM protease, 3 μM substrate and 5 μM Zn^2+^. The slopes of the linear regions of the progress curves are obtained as the activity of these variants. For three significantly improved variants other than Zn45_v2, kinetics were measured as described in section 3.9 and reported in Extended Data Fig. 9e.

##### 4.20. Conditional activation of cytokine mimic by de novo protease

To activate Neo2-M-Z, we added Zn45_v2 at 150 nM, Zn45_v2 KO at 150 nM, or

Zn45 at 1.5 μM to 10 μM Neo2-M-Z in 40 mM HEPES, 50 mM NaCl, 5 μM

ZnSO4 and incubated the mixture at 37 ˚C for 2 h.

For the MMP2 control, pro-MMP2 (Sino Biological, Cat. No. 10082-HNAH) was

activated with 4-Aminophenylmercuric acetate (APMA, MedChemExpress, Cat.

No. HY-148905) in MMP2 activation buffer containing 50 mM Tris, 10 mM CaCl2,

150 mM NaCl, and 0.05% (w/v) Brij-35 for 1 h at 37 °C. After activation, 150 nM

MMP2 was incubated with 10 μM Neo2-M-Z in the same buffer for 2 h at 37 °C.

Following protease treatment, the reaction mixtures were added directly to

HEK-Blue IL-2 cells (InvivoGen hkb-il2-2) for the activation assay according to

the manufacturer’s instructions.

##### 4.21. EGFR antagonist assay

For de novo proteases, the enzyme was added to 10 μM of miniprotein binder to reach a final concentration of 3000 nM Zn45, 3000 nM Zn45 KO, 300 nM Zn45_v2 and 300 nM Zn45_v2 KO. The cleavage was carried out in 50 mM NaCl, 40 mM Hepes, 5 uM ZnCl_2_, pH 8, at 37 °C for 2h.

For MMP2, cleavage of the mask from the miniprotein binder was carried out in vitro using pro-MMP2 (10082-HNAH, Recombinant Human MMP2 Protein, HPLC-verified, Sino Biological, Beijing, China). The enzyme at 1.2 mM was first activated with 2.5 mM of 4-Aminophenylmercuric acetate (APMA, Sigma Aldrich) in TTC buffer (Tris-triton-calcium: 50 mM Tris-HCl pH 7.5, 1 mM CaCl_2_, 0.05% Triton X-100) for 2h at 37 °C. After activation, the enzyme was added to 10 μM of miniprotein binder to reach a final concentration of 160 nM. The cleavage was carried out in TTC buffer at 37 °C for 2h.

The cleavage percentage was assessed by LC-TOF MS.

U-87 cells expressing EGFR were seeded in 12-well plates until confluency was reached. After that, supplemented DMEM medium was aspirated, and cells were washed twice with PBS. Then, DMEM serum-free medium was added, and U-87 cells were serum-starved for 48 h. Following the incubation period, the starvation medium was aspirated, and cells were treated with 700 nM of the corresponding miniprotein binder for 1 h at 37°C, 5% CO_2_. Right after this, cells were stimulated with EGF 1 nM (PHG0315, Gibco) for 15 min at 37°C, 5% CO_2_. Afterwards, the medium was aspirated, and cells were washed once with cold PBS before the lysis treatment. For total protein isolation, U-87 cells were lysed using Pierce™ RIPA buffer (ThermoFisher Scientific, Rockford, IL, USA) supplemented with 7X cOmplete Mini-EDTA free serine/ cysteine protease inhibitor cocktail (Roche Diagnostics GmbH, Mannheim, Germany). All cell-lysate supernatants were collected in microcentrifuge tubes and quantified using the Pierce ™ BCA Protein Assay kit (ThermoScientific). After this, 4X Laemmli sample buffer (Bio-Rad), supplemented with 10% β-mercaptoethanol, was added to all samples before being heated at 95°C for 5 min. Cell lysates were resolved by SDS-PAGE, transferred to low PVDF membranes and blotted using the Trans-blot Turbo Mini Transfer kit (Bio-Rad). The membranes were blocked in TBS-T (Tris Buffer Saline: 50 mM Tris, 138 mM NaCl, 2.7 mM KCl, pH 8.0 + 0.1% Tween-20) with 5% BSA for 1 h, then, incubated overnight at 4°C, under constant stirring, with the Phospho-p44/p42 MAPK Erk 1/2 (1:1000, Cell Signaling Technologies) and β-tubulin (1:1000, Cell Signaling Technologies) primary antibodies. The next day, membranes were washed three times with TBS-T before being incubated with Goat Anti-Rabbit IgG StarBright Blue 700 (1:2500, Bio-Rad) and Goat Anti-Mouse IgG StarBright Blue 520 (1:2500, Bio-Rad) secondary antibodies, respectively, for 1 h at room temperature. Finally, the membranes were washed five times with TBS-T and visualized using the ChemiDoc MP Imaging System System. Densitometric analysis of membranes was carried out using the Image Lab software 6.1 (Bio-Rad).

#### 5. Molecular dynamics and machine-learning–based quantum mechanical calculations for the retrospective analysis of design activity

##### 5.1. Molecular dynamics simulations of the Michaelis complex

Molecular dynamics simulations were performed using the GPU-accelerated AMBER 24 (Case 2024) software package (pmemd.cuda; Götz 2012, Le Grand 2013). Force field parameters for the zinc metalloprotein active site were generated using the MCPB.py program (version 2024; Li 2016). The zinc ion coordination shell was defined using a tetrahedral metal center coordinated by the active site residues. In the design models, derived from the predicted AlphaFold3 protein-substrate complex, these atoms were the Nε2 atoms of His76 and His80, the Oε1 atom of Glu156, and the oxygen of the coordinating water molecule (WAT190). In the native model, derived from PDB structure 1QJI, these atoms were the Nε2 atoms of His92, His96, and His102, and the oxygen of the crystallographic water molecule (WAT207). Small (zinc-coordinating residues) and large (first- and second-shell) cluster models of the metal site were constructed for quantum mechanical calculations. Geometry optimization and frequency calculations were performed on the small cluster models at the B3LYP (Becke 1988, Becke 1993, Perdew 1996) /6-31+G(d) level of theory using Gaussian16, and the resulting Hessian was used by MCPB.py to derive bonded force field parameters (equilibrium bond lengths and angles, and force constants) for the Zn²⁺ center via the Seminario method (Seminario 1996). RESP charges (Bayly 1993, Cornell 1993) for the metal-coordinating residues were fit to the electrostatic potential computed at the same level of theory for the large cluster model. The protein was described using the ff14SB force field (Maier 2015). The solvated system topology and coordinate files were assembled using the tleap module of AmberTools.

The protein complex was immersed in a pre-equilibrated truncated octahedral box with an 8 Å buffer of TIP3P (Jorgensen 1983) water molecules. Explicit Na⁺ and Cl- ions were added to neutralize the total charge of the system. Two sequential energy minimization steps were performed, each consisting of 2,500 steps of steepest descent followed by 2,500 steps of conjugate gradient, for a total of 5,000 cycles per step. The first minimization was performed with strong harmonic positional restraints (500 kcal mol⁻¹ Å⁻²) applied to all non-hydrogen, non-solvent, and non-ion atoms; the second applied weaker restraints (50 kcal mol⁻¹ Å⁻²) to backbone heavy atoms (C, Cα, N, O) only.

Following minimization, a two-step equilibration protocol was employed. The first equilibration step heated the system from 0 K to 300 K over 300 ps under constant-volume (NVT) conditions with a 1 fs time step and harmonic restraints (5 kcal mol⁻¹ Å⁻²) on backbone heavy atoms (C, Cα, N, O), using Langevin dynamics (γ = 5.0 ps⁻¹) for temperature regulation. A second equilibration was then conducted for 50 ns under constant-pressure (NPT) conditions at 300 K and 1 atm using a 2 fs time step; pressure was regulated using the Monte Carlo barostat (taup = 10.0 ps). Throughout all MD simulations, bonds involving hydrogen were constrained using the SHAKE algorithm, and long-range electrostatics were computed using PME (Essmann 1995) with an 8 Å direct-space cutoff applied to both Lennard-Jones and electrostatic interactions. Production trajectories were run in pentaplicate for 500 ns each. Production runs were performed under the same NPT conditions as the second equilibration, except with a shorter barostat relaxation time constant (taup = 5.0 ps, reduced from 10.0 ps). Coordinates were saved every 200 ps. Trajectories were processed and analyzed using the cpptraj (Roe & Cheatham 2013) module of AmberTools, including stripping of solvent and ions and RMSD-fitting to the minimized structure.

##### 5.2. Machine-learning–based quantum mechanical analysis of the reaction pathway

To elucidate the structural and energetic origins of the activity differences between designed and native enzymes, we calculated activation energy barriers for the catalytic reaction. These barriers serve as exponential predictors of the reaction rates for elementary steps, allowing for a direct comparison of catalytic efficiency. For this analysis, we assumed proton abstraction and nucleophilic attack to be rate-determining for the overall catalytic cycle.

**System Preparation:** Initial structural models of the native and designed enzymes were generated using AlphaFold 3. The structures were protonated at neutral pH using pymol. The protonation states of Zn(II) ion-chelating residues and residues involved in proton transfer are manually adjusted. To reduce computational cost while maintaining chemical accuracy, the systems were truncated to cluster models centered on the active site. These clusters included the substrate and all residues with at least one atom within 5 Å of the substrate.

Machine Learning Potential: Reaction pathways were modeled using a reactive, polarizable electrostatic foundation model based on an extension of the MACE architecture. This force field was trained on the OMOL25 dataset, comprising over 100 million density functional theory (DFT) calculations at the ωB97M-V/def2-TZVPD level of theory, ensuring high-fidelity description of bond-breaking and bond-forming events.

**Transition State Search:** Reaction barriers were calculated using the Nudged Elastic Band (NEB) method implemented in the Atomic Simulation Environment (ASE). Geometry optimizations were performed using a preconditioned L-BFGS optimizer. First, the reactant and product states for each enzyme variant were optimized to a maximum force convergence (Fmax) of 0.025 eV/Å. A reaction path was then interpolated using 11 images. The band was initially relaxed using standard NEB until forces fell below 0.5 eV/Å, at which point the Climbing Image (CI-NEB) scheme was enabled to rigorously identify the transition state. The CI-NEB calculations were continued until a final convergence of Fmax<0.05 eV/Å was achieved.

Barrier Analysis The activation energy (ΔE‡) was defined as the energy difference between the highest energy image (transition state) and the initial reactant minimum. These calculated barriers can be used to rank the catalytic potential of the designs and predict relative enzyme activities.

### Supplementary Discussions

#### 1. Choice of the theozyme

The successful design of catalytic machineries depends significantly on the capability of the prediction tool to assess the precision of the catalytic residues at atomic levels. It is a known trend that deep-learning based structural prediction tools show higher confidence and better accuracy in structured regions than loop and disordered regions ( Tunyasuvunakool 2021). Therefore, we chose the catalytic residues from aminopeptidase N, where all five catalytic residues are on structured domains. However, as Aminopeptidase N is an exopeptidase, the substrate scope is significantly limited by the cleavage preference provided by its native substrate conformation. To relax this constraint, we used the substrate conformation of an endopeptidase, astacin, as the other component of the theozyme.

#### 2. Representation of the water molecule in the prediction of the ES complex using the Zn(II)-water strategy.

To implement the Zn(II)-water strategy of our original design campaign, we modeled the catalytic water molecule and the Zn(II) ion as the same entity using the smiles string [Zn+2].O. Therefore, the water molecule was “constrained” in the Zn(II) ion coordination sphere. However, the catalytic water can also be modeled as a second separate ligand entity. We did not initially pursue this approach because we anticipated that increasing the number of ligands might complicate AF3’s ability to predict the complex structure accurately and only tested the “unconstrained” water strategy retrospectively on a small set of designs. Interestingly, the AF3 predictions with constrained and unconstrained water were very similar in the set of designs we tested, where both active and inactive designs were included (Table S1). As we only retained the sequences of ordered designs, we don’t have very bad designs that didn’t pass some initial in silico screens to test for and show negative results. Nevertheless, using the “unconstrained” strategy may exploit the capability of AF3 in the prediction of Zn(II)-water coordination and thereby may provide additional predictability than the "constrained" strategy we used in the design campaigns reported in the manuscript.

#### 3. Additional discussions of the crystal structures

##### 3.1. The protease-Zn(II)-substrate complex of ZnO7 (PDB: 11CM)

The Q32 side chain in the crystal structure flips away from the catalytic water, forming three hydrogen bonds with the carbonyl oxygen of D62, the side chain oxygen of T68, and a water molecule in the solution (Supplementary Figure S30), stabilizing this Q rotamer with 100% occupancy. With the wild-type E32, at least the first hydrogen bond cannot form, potentially destabilizing it from the rotamer state of Q32 in the crystal structure. Moreover, according to our catalytic residue KO experiment, the E32 general base in ZnO7 contributes significantly to catalysis: a E32Q substitution completely abolishes activity (Figure 3b). Therefore, the E32 rotamer should at least be transiently present in the catalytically competent state as shown in the design model. Despite the potential crystallography artifact, this same Q to E mutation in the TDPr3, and a few other crystal structures we obtained (Extended Data Fig. 6), did not result in complete flipping of the rotamer, potentially due to better stabilization in the catalytically competent conformation. Therefore, the difference in the side chain rotamers between Q32 in the crystal and the E32 in the design model of ZnO7 may suggest further preorganizing the general base as a potential direction for activity enhancement for this design.

Among all the de novo scaffolds reported in this manuscript, ZnO7 is unique in that the catalytic base can reach the nearest protein surface upon adopting the flipped rotamer. On this protein surface, many polar residues such as T68 were introduced by MPNN to ensure solubility. Consequently, the Q32 residue residue reaches toward such nearby polar residues to form favorable polar interactions. In addition, because residue 32 is sufficiently close to solvent in this state, a water-mediated hydrogen bond can further stabilize the alternative conformation. In the other designs, by contrast, the catalytic base is more deeply buried within the hydrophobic core of the proteins. As a result, the surrounding environment is less permissive to such alternative stabilization: nearby polar side chains are less likely to be present, and solvent water cannot readily access the residue. Thus, ensuring the positioning of the catalytic base in the hydrophobic core of the designed protease may present a potential direction for improvement of the design pipeline.

##### 1.2 The E45Q-substrate complex of TDPr3 (PDB: 11DU)

The E45Q-substrate complex of TDPr3 was prepared by supplementing ZnSO_4_. However, the Zn(II) ion was not resolved in the crystal structure, suggesting suboptimal Zn(II) ion binding of this design as compared to Zn5, ZnO7 and Zn48, where the Zn(II) ion coordination was correctly captured. Resolved as a substrate complex, the peptide bond between two serine residues is positioned at the active site, suggesting a potential cleavage site at ALQS/SWGMMGML. However, this is inconsistent with the designed cleavage site (ALQSS/WGMMGML), and is inconsistent with the mass-spec detected cleavage site (ALQSS/WGMMGML and ALQ/SSWGMMGML), potentially suggesting the crystallography process, such as the E45Q mutation, high substrate concentration, and the loss of the Zn(II) ion could have led to a different binding registration.

##### 1.2 The E146Q-Zn(II)-substrate complex of Zn48 (PDB: 11CU)

Zn48 is a very slow hit discovered in the cis screen (digested fraction = 0.12 after 12 h incubation at 37 °C) which did not translate to the trans screen. We obtained this crystal structure of this design to inspect potential reasons for its low activity. Unlike the other more active designs such as Zn5 or ZnO7, where the oxyanion stabilizing tyrosines are resolved with the correct rotamer states, The oxyanion stabilizing residue of Zn48, Y34, adopts a rotamer state that is incompetent to donate a hydrogen bond to the transition state oxyanion, potentially lowering the activity of this design. The substrate binding registration in the crystal structure suggests a cleavage site at PLRFRSGY/GS. However, this is inconsistent with the design cleavage site (PLRFR/SGY) and mass-spec detected cleavage site (PLRFR/SGY), potentially suggesting the crystallography process, such as the E146Q mutation and high substrate concentration, could have led to a different binding registration.

#### 4. Limitations of the machine learning force field-based quantum mechanical calculations of the activation energy barriers.

Based on several QM/MM calculations for the reaction pathways of metalloproteases, nucleophilic attack (Pelmenschikov 2002, Varghese 2023), hydrogen-bond rearrangement (Chen 2021), product release (Vasilevskaya 2015) can all be the rate limiting step. These results show that even in metalloproteases using the similar catalytic residues (His/Glu for Zn(II) coordination, Glu as general base, Zn-bound water as nucleophile) and similar reaction steps, the detailed energy landscape may be very different and result in different rate-limiting steps.

As our theozyme contains components from astacin, we chose to calculate the energy landscape of the nucleophilic attack step, which was suggested to be the speed-limiting step of astacin (Chen 2012). This is sufficient to provide an estimation of the lower bound of the activation energy. More importantly, for structure-based enzyme design methods, it points to contributions to catalytic activity missed in pure geometric assessments – subtle differences in catalytic residue placement, and the surrounding chemical environment of the active site.

However, the quantitative prediction of activity using free energy predictors requires careful considerations of long-range interactions, the full reaction path, and entropic contributions, which are not included in our current setup. For example, among the four residues that differ between Zn45_v2 and Zn45, only one is within the truncated region used for the calculation. Consequently, the MLFF-based NEB simulation of Zn45_v2 yielded an activation energy barrier of 17.1 kcal/mol, very close to the value of Zn45 at 17.3 kcal/mol. Thus the activity difference between Zn45 and Zn45_v2 is not captured by the MLFF-based NEB simulation. Nevertheless, the molecular dynamics simulations we performed nicely captured the effects of substrate and catalytic residue preorganization, which correspond to the factors missing from the NEB simulations.

#### 5. Introduction of substrate-stabilizing residues by MPNN reduces activity

We observed an interesting preference of LigandMPNN for the sequence design of metalloprotease in two of our designs, Zn45 and ZnO36. In both designs, MPNN introduced a hydrogen bond donor extremely close to the substrate carbonyl oxygen (R79 in ZnO36 and Y152 in Zn45). When these two hydrogen bond donors were replaced by other residues incapable of hydrogen bonding, such as R79A in ZnO36 and Y152I in Zn45, activity of the enzymes both increased significantly. LigandMPNN was trained as a general model for sequence design; it is optimized to introduce protein-protein interactions wherever possible. However, for the design of an efficient catalyst, hydrogen bonding with a substrate atom extremely important in the reaction mechanism can be distracting: an ideal hydrogen bond that promotes catalysis should preferential stabilize the transition state over the ground state (Du 2025), yet the assessment of such differential stabilization was not incorporated in our current design pipeline. Careful evaluation of the geometry, or energetic contributions of these important hydrogen bonds across multiple states can thus further improve the activity of one-shot designs.

#### 6. Predictability of AF3 and MPNN in the mutagenesis space of Zn45-S_ZnO36

In the single mutation space of Zn45-S_ZnO36, we noticed that AF3 can predict some deleterious mutations with metrics such as complex pLDDT and minimum ipae. If a filter was set at complex pLDDT > 94 and minimum ipae< 1.5, no significantly improving single variants will be removed. Therefore, AF3 can be used as an effective filter for the in silico prescreen of single-site saturation mutagenesis libraries of de novo metalloproteases. However, above these thresholds for both metrics, no more correlation can be observed between wet-lab activity and these computational metrics, as shown in Extended Data Fig. 6g. We also investigated MPNN probability as another potential predictor for activity. However, when we masked individual positions where beneficial mutations were discovered and redesigned the amino acids for each such position, we only recovered the wild-type amino acids with high probability. The sum of the probabilities of all beneficial mutations at each such position is consistently < 20% and in most cases < 5%, as shown in Extended Data Fig. 6f.

### Supplementary Figures

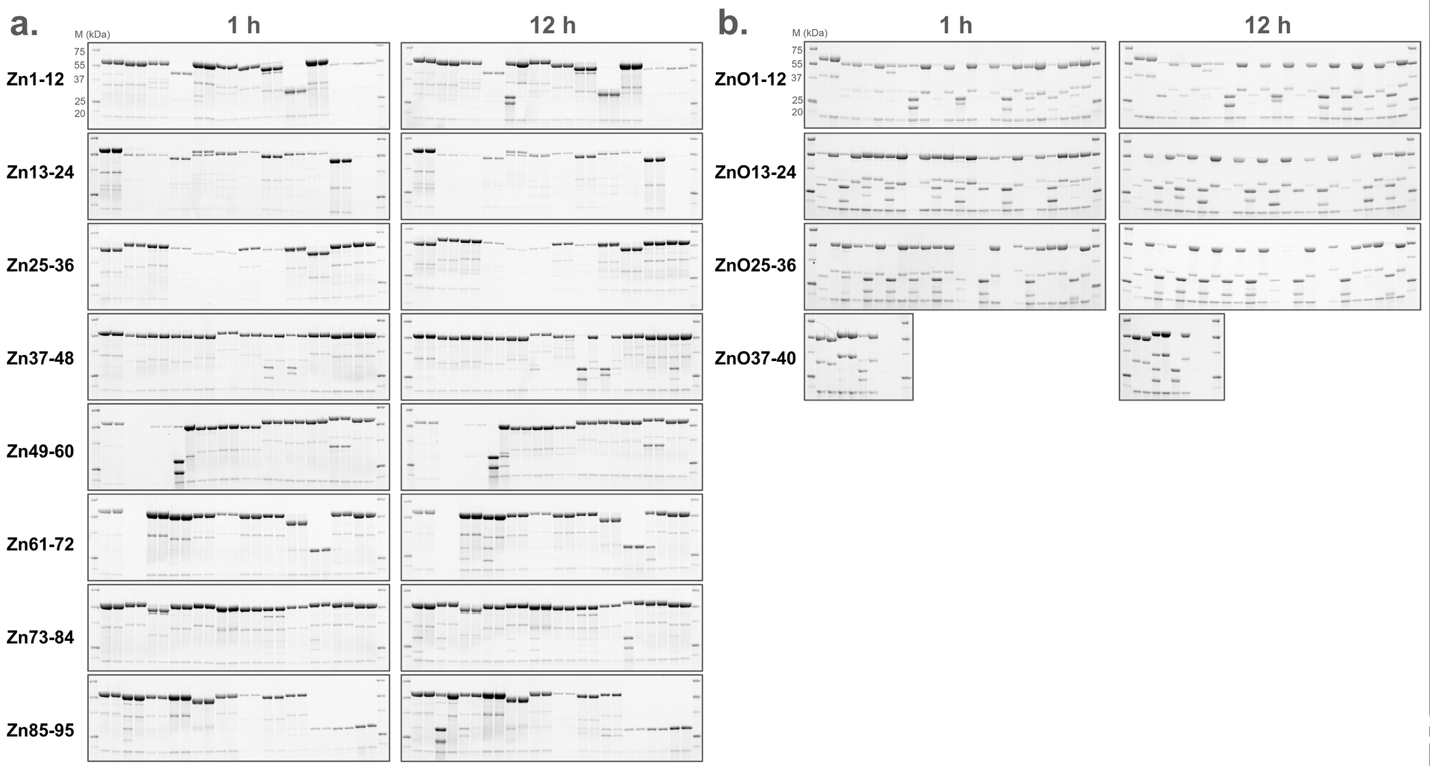

##### Figure S1. Gel images for the cis screen of 135 designs.

a. 1 h and 12 h time points for the 95 Zn designs. b. 1 h and 12 h time points for the 40 ZnO designs. The 96th sample on the same gel with Zn85-95 is a TEV positive control to indicate the expected position of the protease fragment after cleavage. Each design is shown in pairs of the +Zn and -Zn groups. A zoomed-in view for the grouping of each design is shown in Fig. S4.

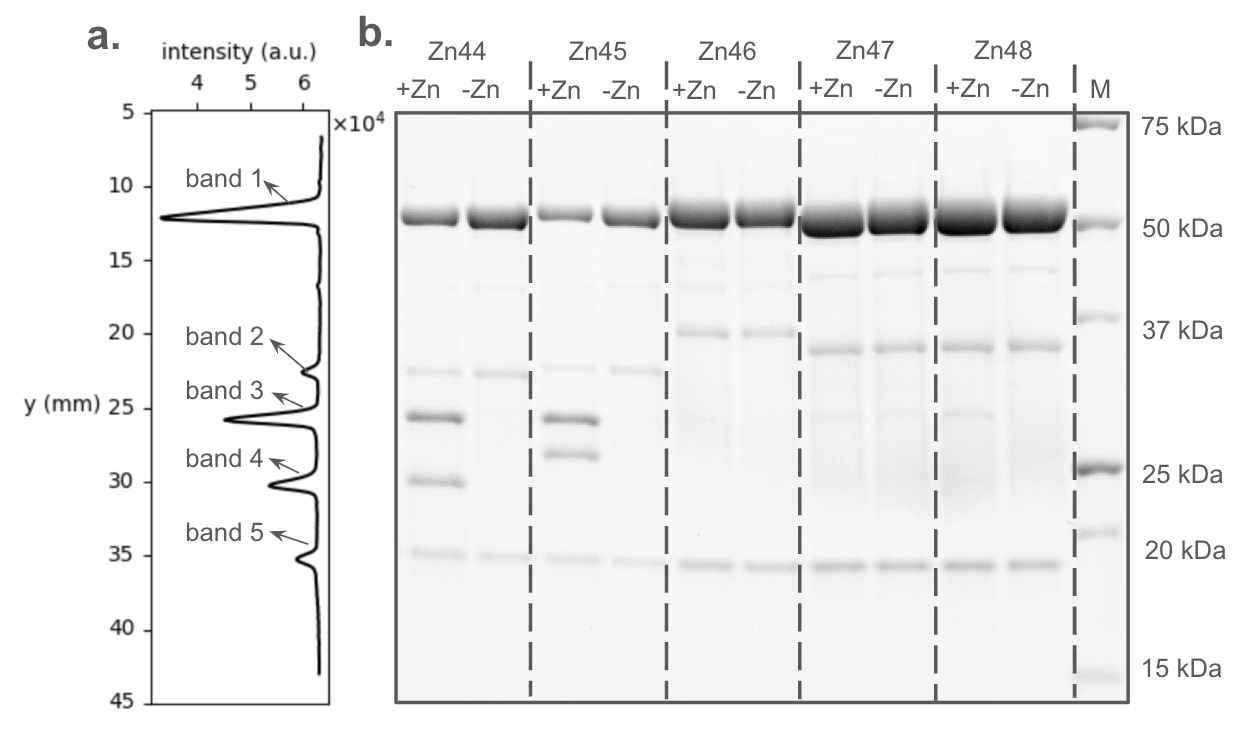

##### Figure S2. Densitometry analysis to determine the digested fraction of Zn44 in the cis screen after incubation at 37 °C for 1 h.

a**.** Intensity value of the Zn44 +Zn lane plotted as a function of y position in pixels. Band intensity gray values are shown in arbitrary units (a.u.) b. Gel image showing the +Zn and -Zn groups for five designs and a marker lane in the end. The digested fraction of Zn44 is calculated as the the sum of the intensity gray values corresponding to band 3 and band 4, divided by the sum of the intensity gray values corresponding to band1, band 3 and band 4. Band 2 and band 5 are likely byproducts of mScarlet chromophore maturation (Wei 2015) and are thereby omitted from the quantification.

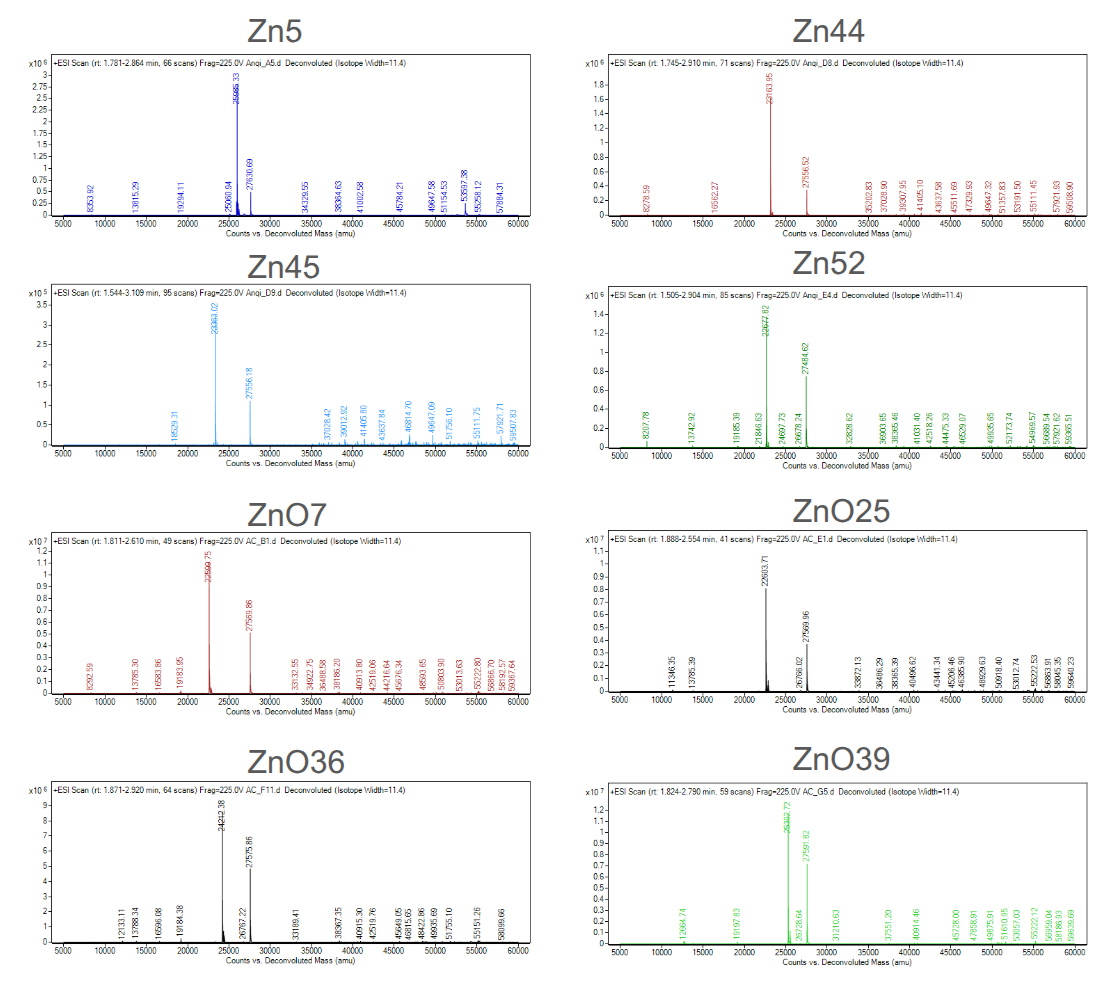

##### Figure S3. Representative deconvoluted mass spectra of selected hits from the cis screen.

In each case the spectrum has two dominating peaks at the expected molecular weights of the N terminal fragment and the C terminal fragment after cleavage.

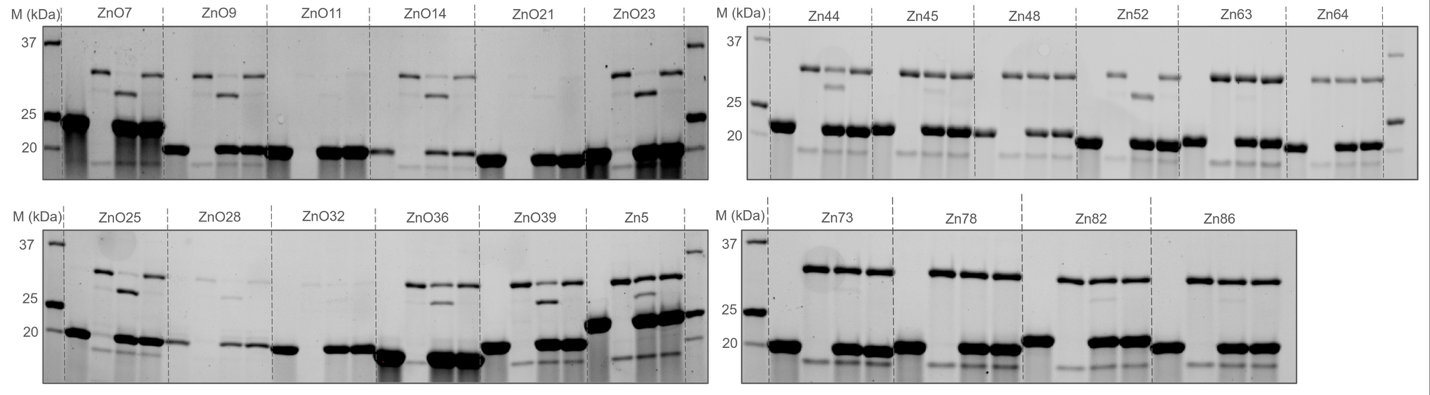

##### Figure S4. SDS-PAGE gel images for the trans screen of 40 designs.

Each design is shown in four groups of protease alone, substrate alone, protease + substrate + Zn and protease + substrate - Zn. A zoomed-in view for the grouping of each design is shown in Fig. S7.

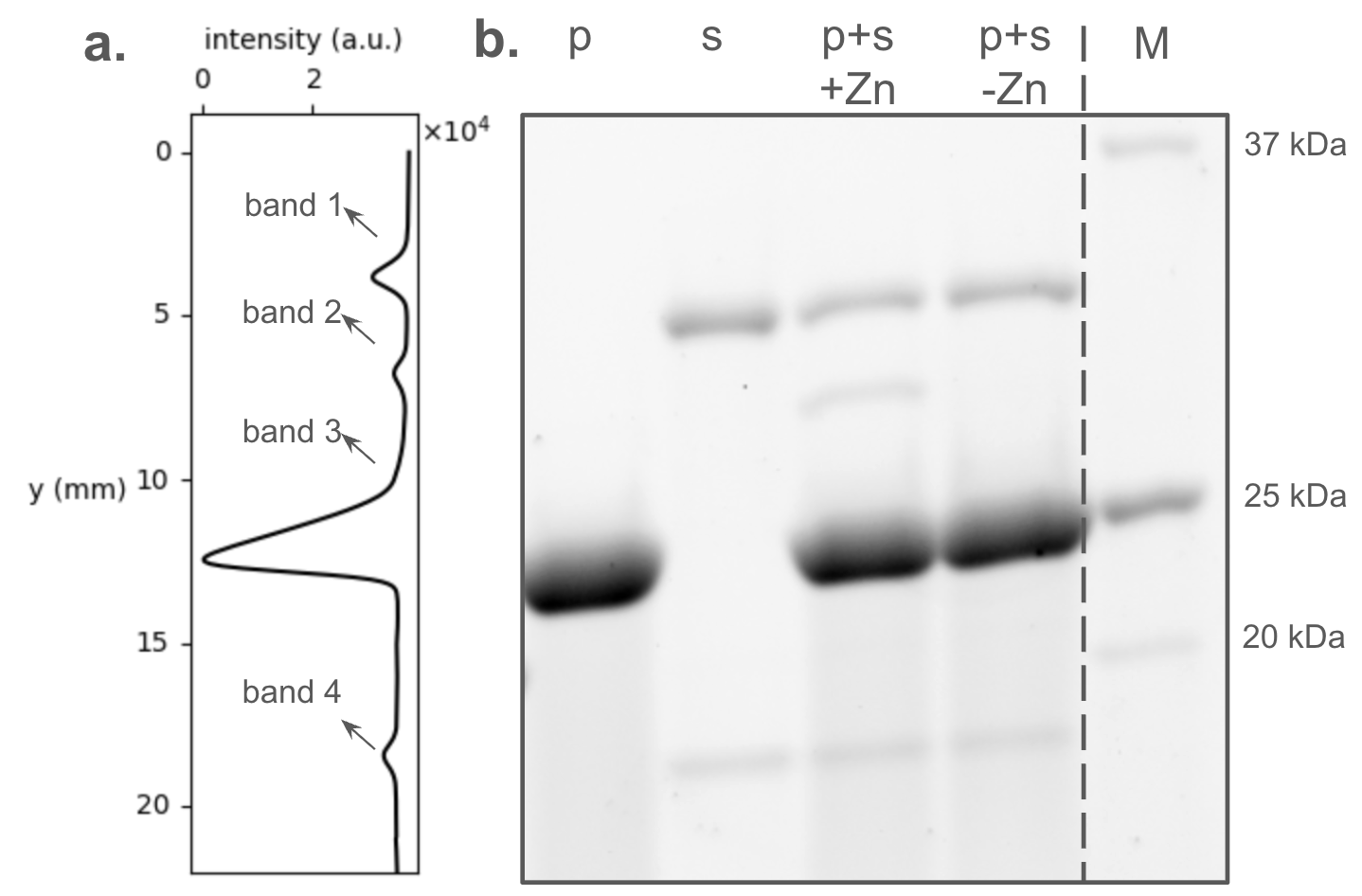

##### Figure S5. Densitometry analysis to determine the digested fraction of Zn5 in the trans construct screen after incubation at 37 °C for 1 h.

a**.** Intensity value of the Zn5 p+s+Zn lane plotted as a function of y position in mm. Band intensity gray values are shown in arbitrary units (a.u.) b. Gel image showing the protease alone (p), the substrate alone (s), protease + substrate+Zn (p+s+Zn) and protease + substrate-Zn (p+s-Zn) groups and a marker lane in the end. The digested fraction of Zn5 is calculated as the intensity gray value corresponding to band 2, divided by the sum of the intensity gray values corresponding to band1, and band 2. Band 3 corresponds to the protease. Bond 4 is likely byproducts of mScarlet chromophore maturation (Wei 2015) and is thereby omitted from the quantification.

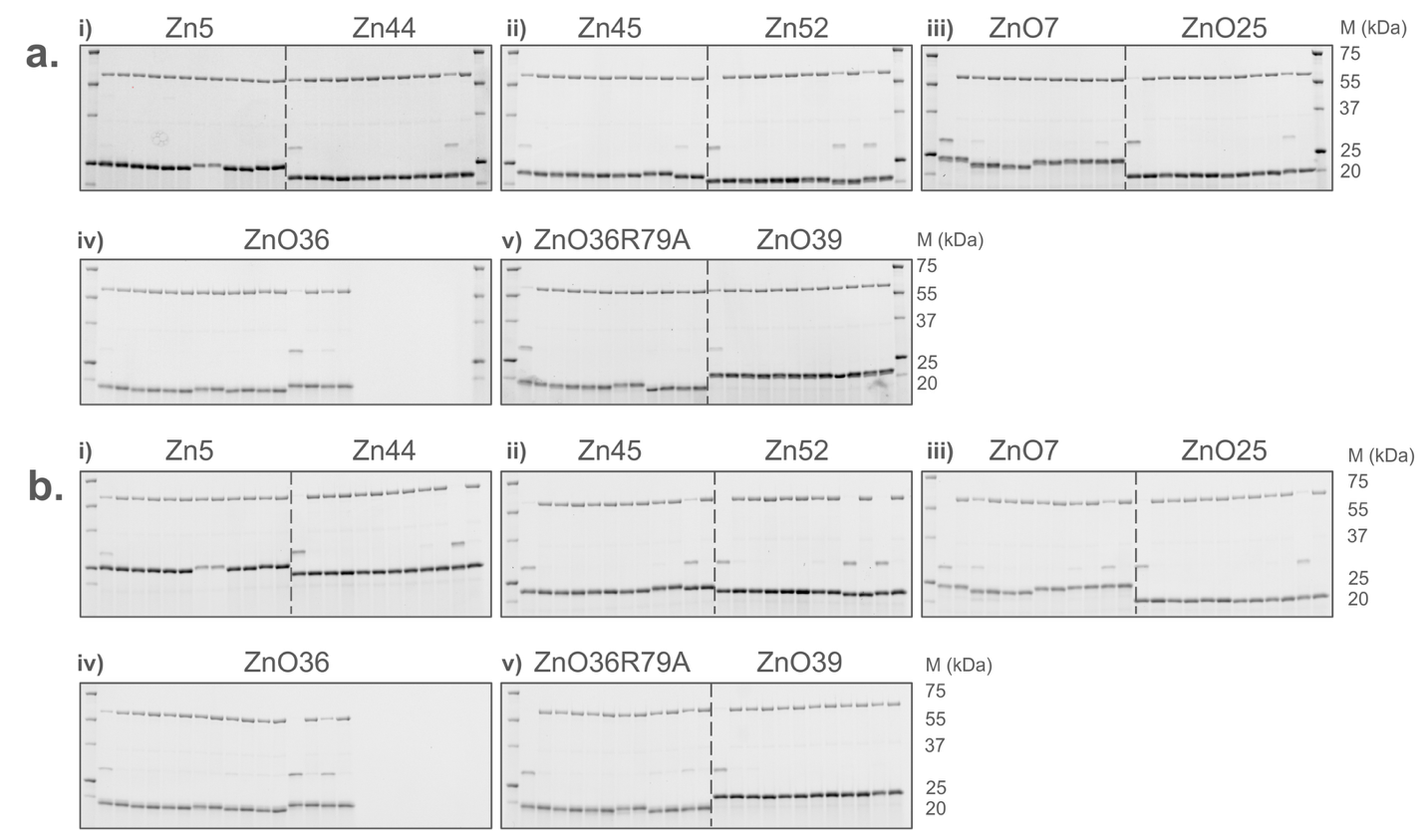

##### Figure S6. Single catalytic residue knockout experiment. SDS-PAGE gel images from one of the three replicates

a. Incubation at 37 °C for 1 hour b. Incubation at 37 °C for 8 hours. Designs on gels (12 lanes on the left, 12 lanes on the right): i) Zn44, Zn52; ii) ZnO7, ZnO25; ii) ZnO39, ZnO36; iv) ZnO36R79; v) Zn5, Zn45. For each design, the mutated catalytic residues (left to right) are: first Zn(II) chelating residue (histidine), second Zn(II) chelating residue (histidine), third Zn(II) chelating residue (glutamate), general base (glutamate) and the oxyanion stabilizer (tyrosine). +Zn and -Zn groups for each variant are paired in neighboring lanes. Zoomed-in gel images with detailed description of grouping is shown in Fig.S9.

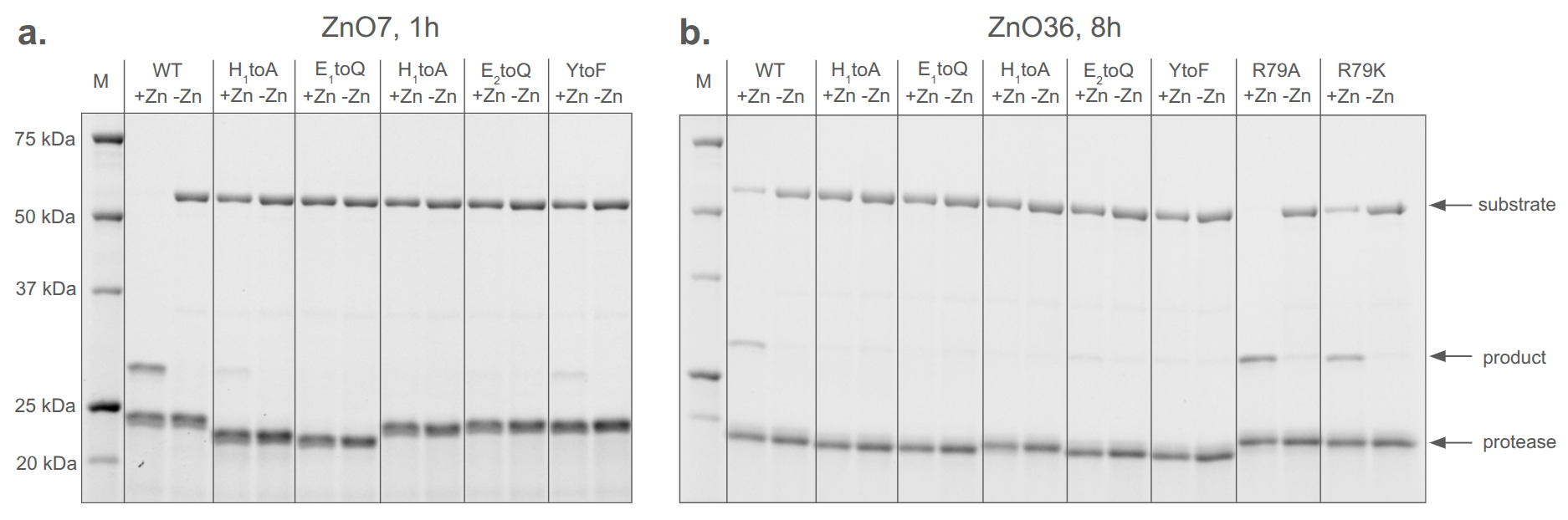

##### Figure.S7 Detailed grouping of gel lanes for the single catalytic residue KO experiment.

a. For ZnO7 at the 1 h time point. The gel lanes for the following designs at both time points are organized in the same manner as ZnO7 at the 1 h time point: Zn5, Zn44, Zn45, Zn52, ZnO36, ZnO36R79K, ZnO39. b. For ZnO36 at the 8 h timepoint.

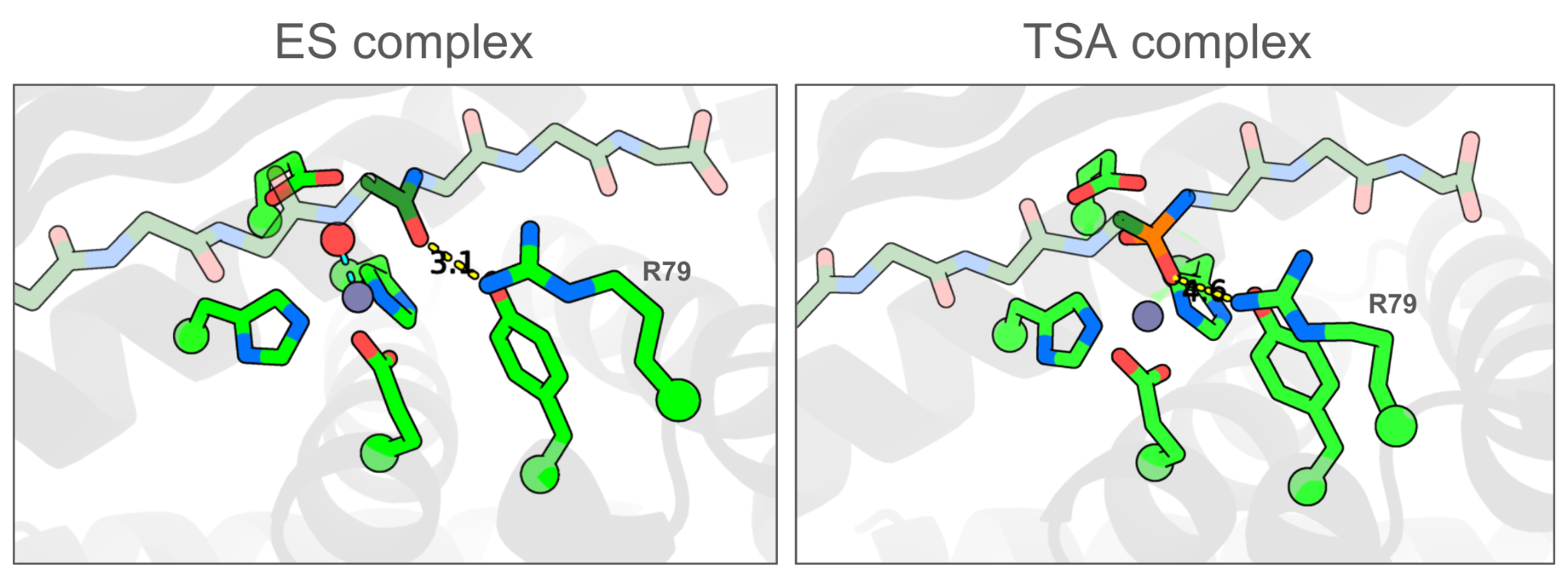

##### Figure S8. Position of R79 on ZnO36. AF3 predictions of the ZnO ES complex and the transition state complex.

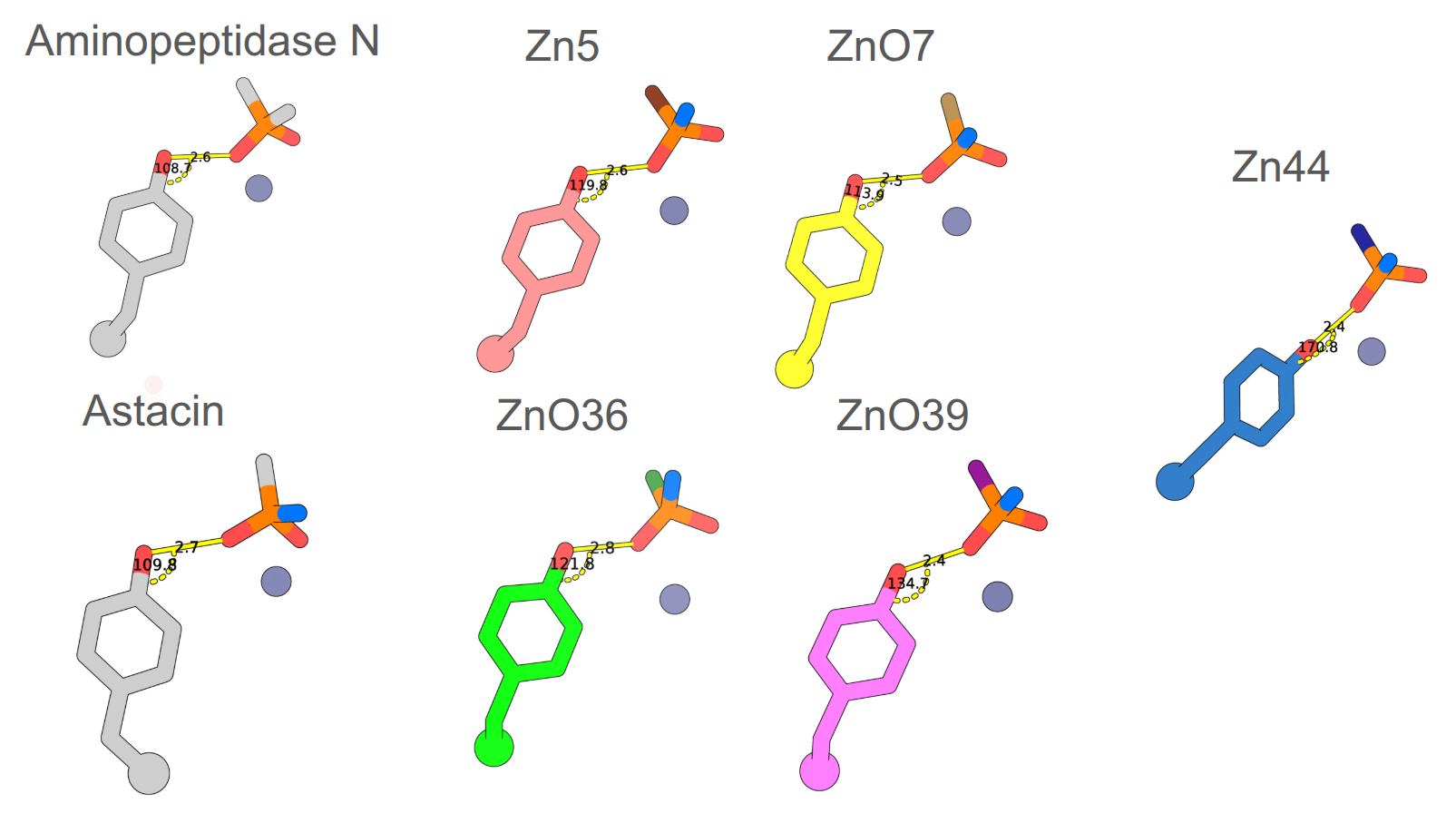

##### Figure S9. Tyrosine hydrogen bond angles in the crystal structures of two native proteases and in the AF3 TSA models of five designs.

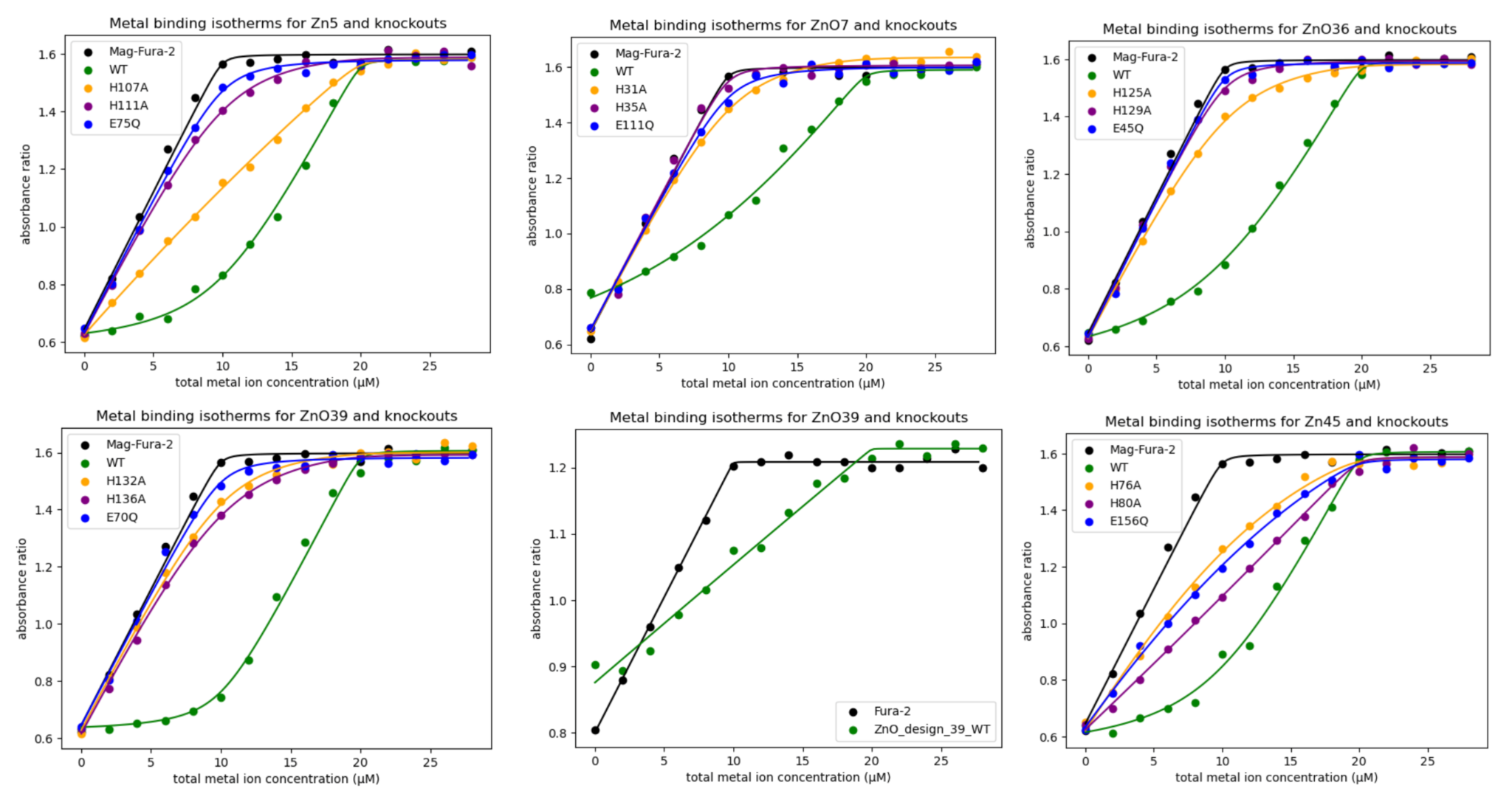

##### Figure S10. Zn(II) binding isotherms for Zn5, ZnO7, ZnO36, ZnO39 and Zn45

For Zn5, ZnO7, ZnO36, Zn45, the binding isotherms were measured using Mag-Fura-2. Because ZnO39 (WT) exhibited strong Zn(II) binding that is at the limit of the sensitivity range of Mag-Fura-2, we assessed it using two ratiometric dyes— a weaker-binding Mag-Fura-2 and a stronger-binding Fura-2. The measured dissociation constants (*K*_d_) are consistent across the two dyes (280 ± 150 pM for Mag-Fura-2 and 330 ± 80 pM for Fura-2), confirming the tight Zn(II) binding of ZnO39.

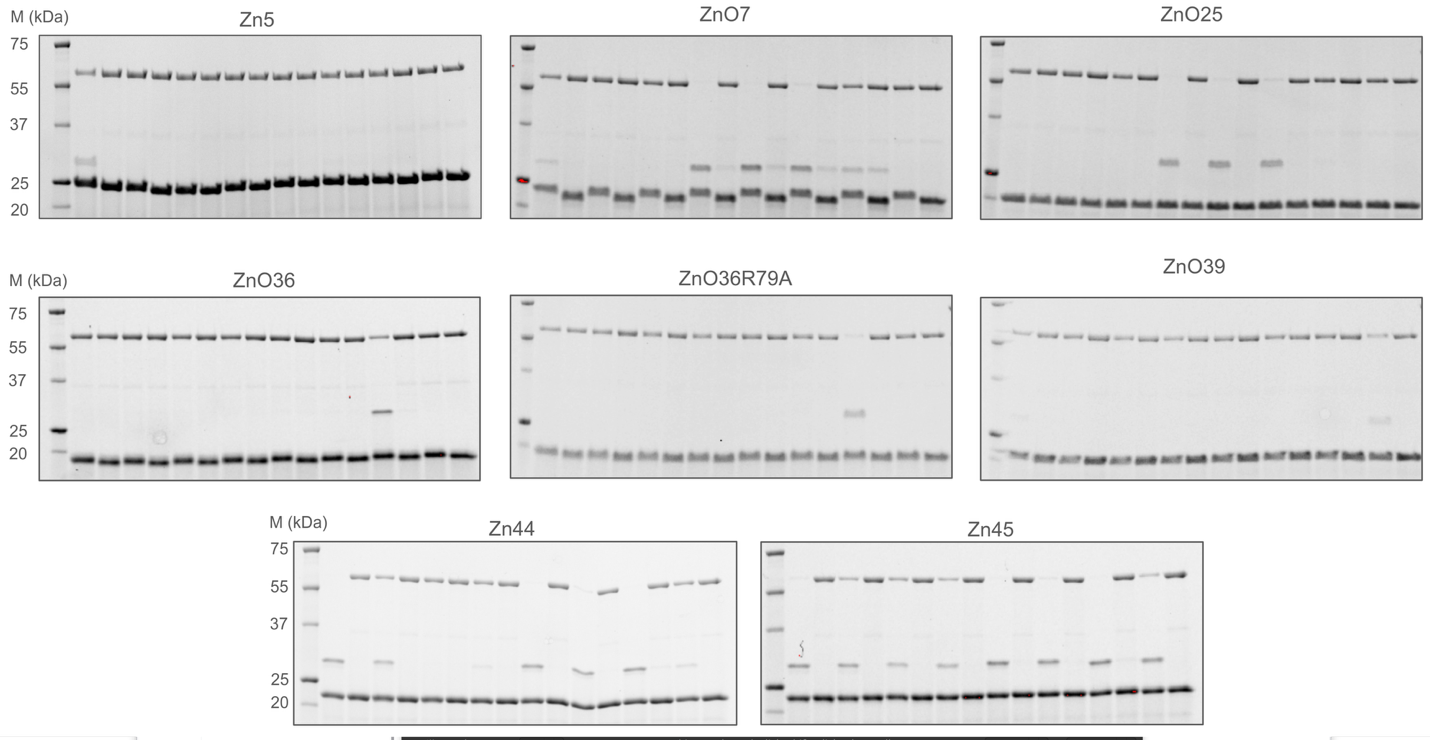

##### Figure S11. cross-reactivity experiment, SDS-PAGE gel images from one of the three replicates.

For each protease, the cleavage of different substrates are on the same gel for direct comparison. The WT proteases and the histidine-to-alanine negative controls are in neighboring lanes for direct comparison. On each gel, the order of substrates are as follows: S_Zn5, S_Zn44, S_Zn45, S_Zn52, S_ZnO7, S_ZnO25, S_ZnO36, S_ZnO39. A zoomed-in gel image with detailed description of grouping is shown in Fig.S11.

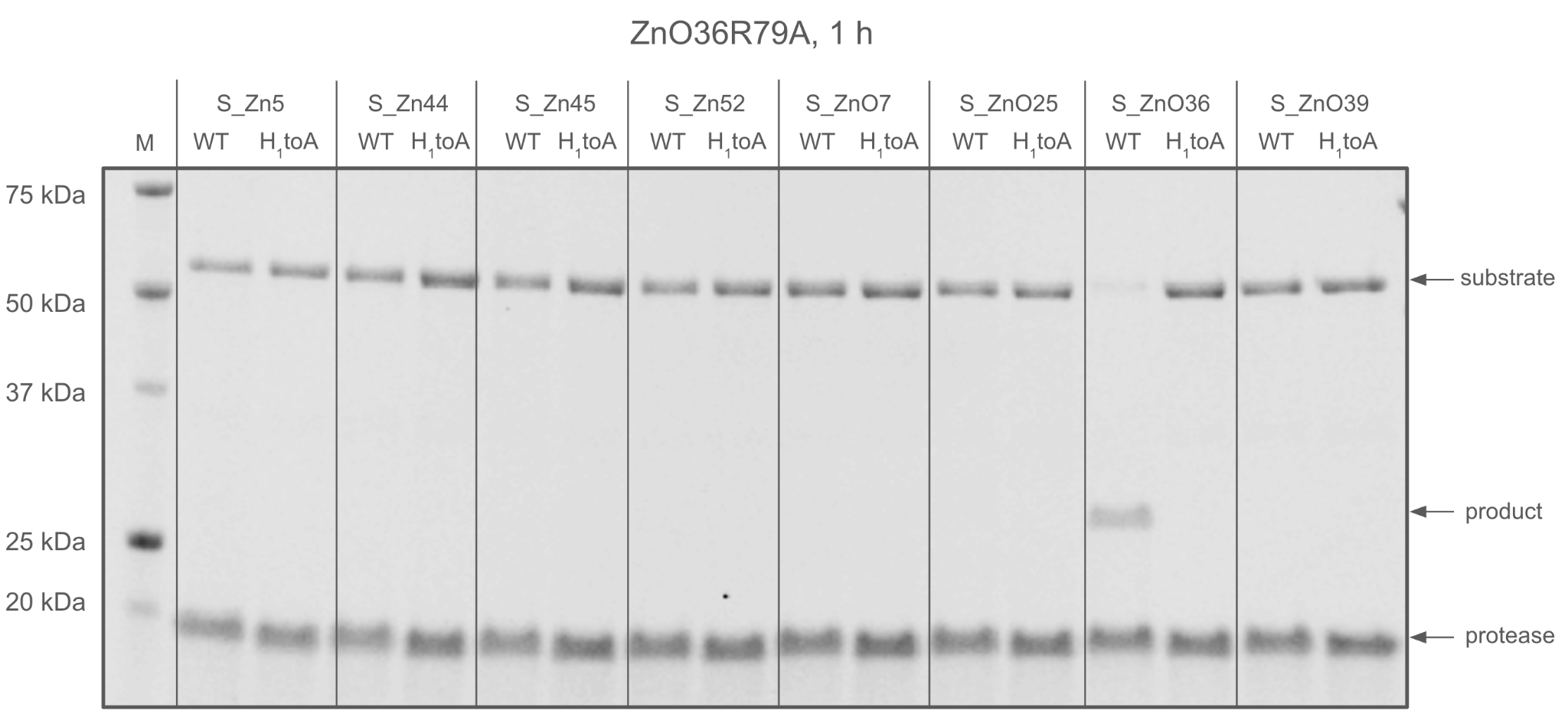

##### Figure S12. Detailed grouping of gel lanes for the cross-reactivity experiment.

The ZnO36R79A protease is used as an example. The gel lane arrangements are the same for all proteases in this experiment.

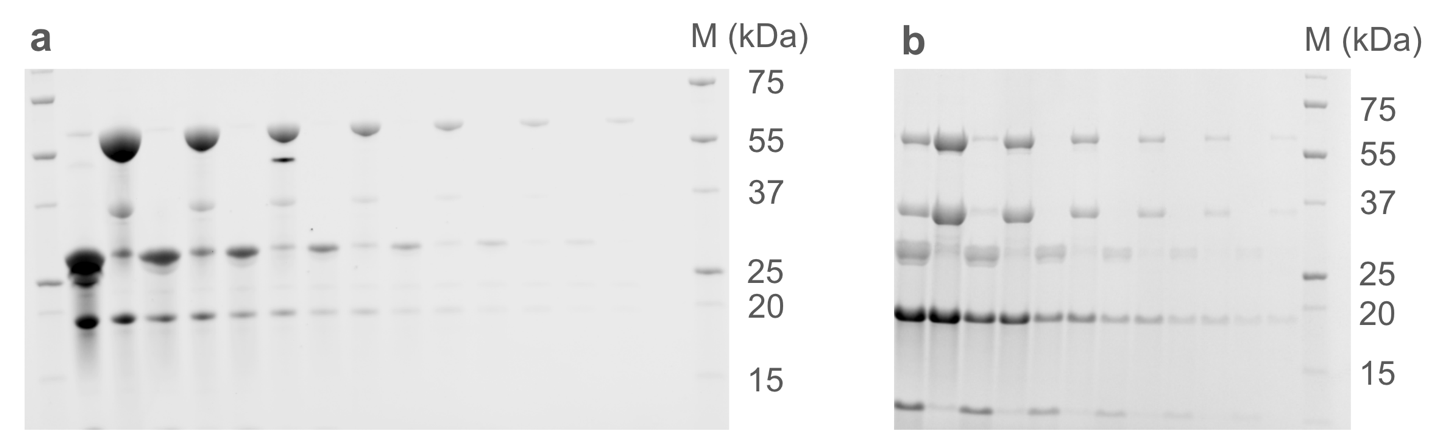

##### Figure S13. Raw SDS-PAGE gel images of the total turnover number experiments for a, Zn45 and b, Zn45_v2, after 12 hours and 8 hours at 37 C, respectively.

For Zn45, the substrate concentrations were 100, 50, 25, 12.5, 6.1, 3.1, 1.6 µM and the protease concentration was 500 nM. For Zn45_v2, the substrate concentrations were 160, 80, 40, 20, 10, 5 µM, and the protease concentration was 20 nM.

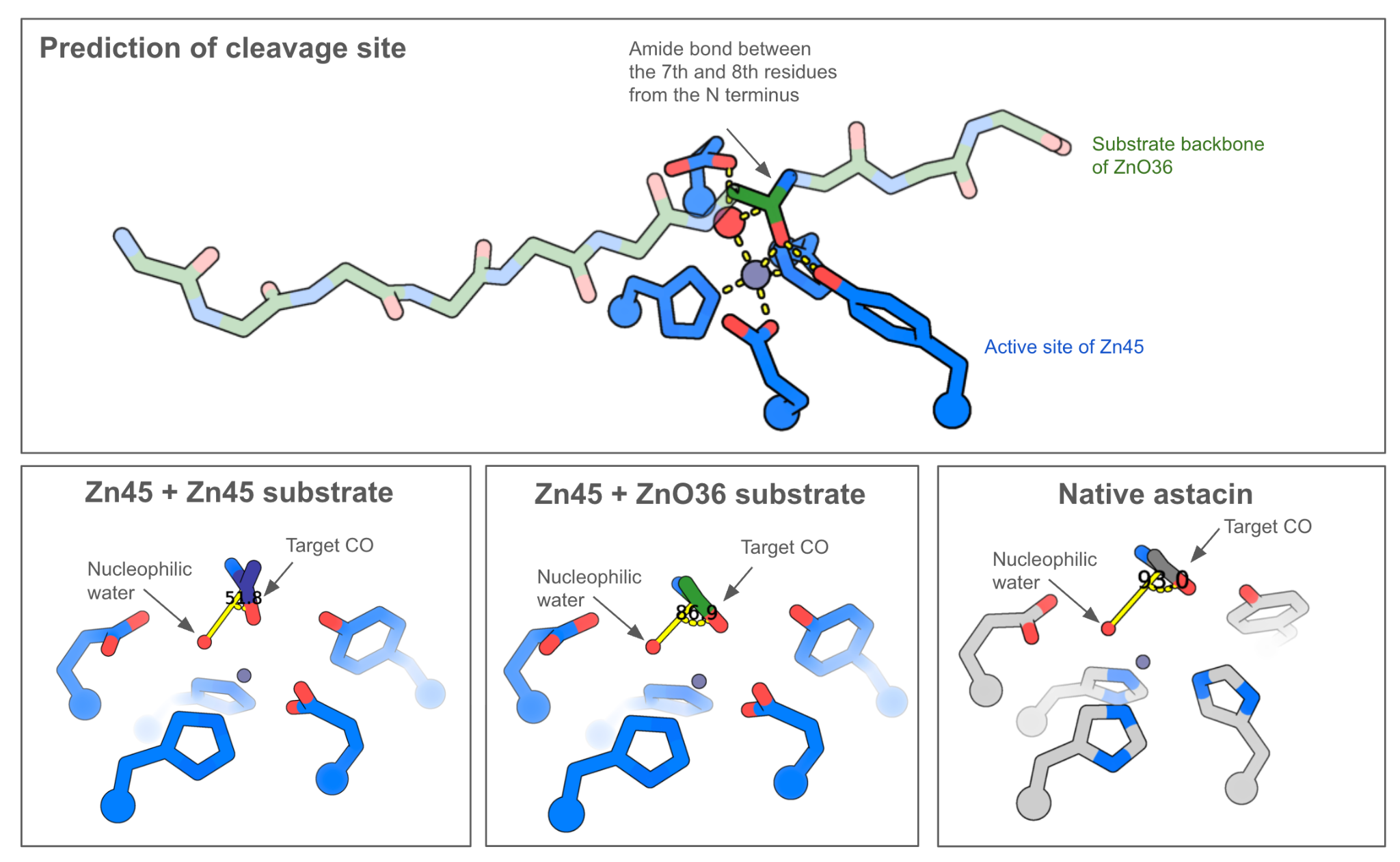

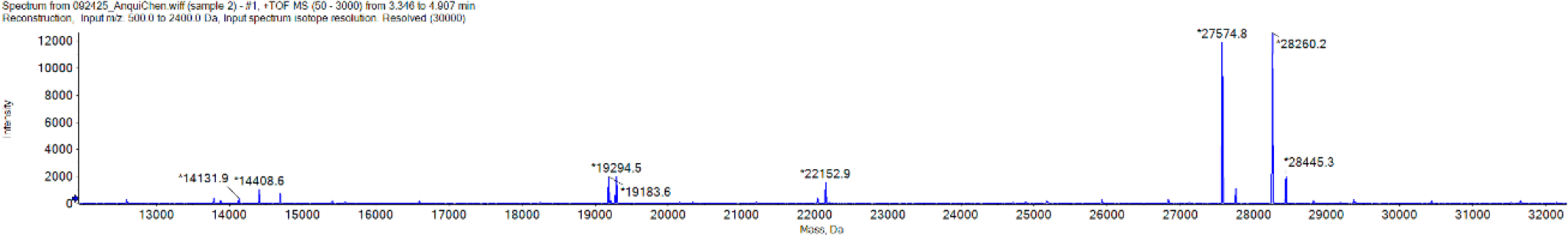

##### Figure S14. The AF3 ES models for the cleavage site of Zn45 cleaving ZnO36 and mass spec.

The tall peaks at 28260.2 and 27574.8 correspond to the expected N-terminal and C-terminal cleavage products from the mScarlet-S_ZnO36-sfGFP substrate, consistent with the expected masses when the cleavage happens between the 7th and 8th residue in the designed substrate sequence ARILRVN/FKI (Table S2).

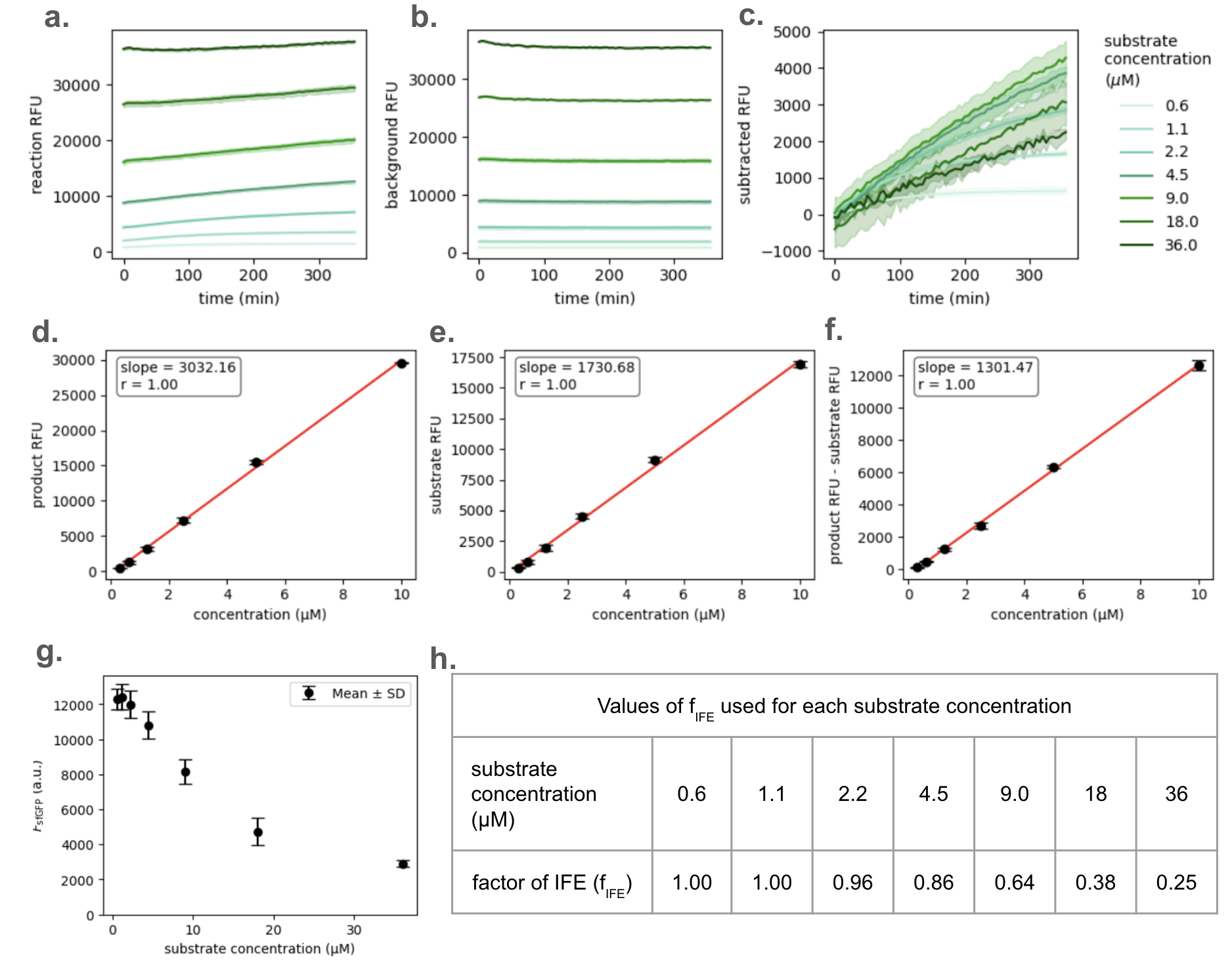

##### Figure S15. Progress curves, standard curves and correction of inner filter effect for the kinetics measurements of ZnO7.

**a**. Reaction progress curves with 500 nM enzyme, **b.** substrate alone control, and **c.** reaction progress curve after subtracting the substrate alone control. **d**. Fluorescence standard curves of the equimolar mixture of sfGFP and mScarlet. **e**. Fluorescence standard curves of the sfGFP-S_ZnO7-mScarlet substrate. **f**. Standard curves for the fluorescence difference between the intact substrate and separated fluorophores. **g.** The contribution of fluorescence by fluorophore (sfGFP) as a function of substrate concentration for the correction of inner filter effect **h.** Factors of inner filter effect used for correction.

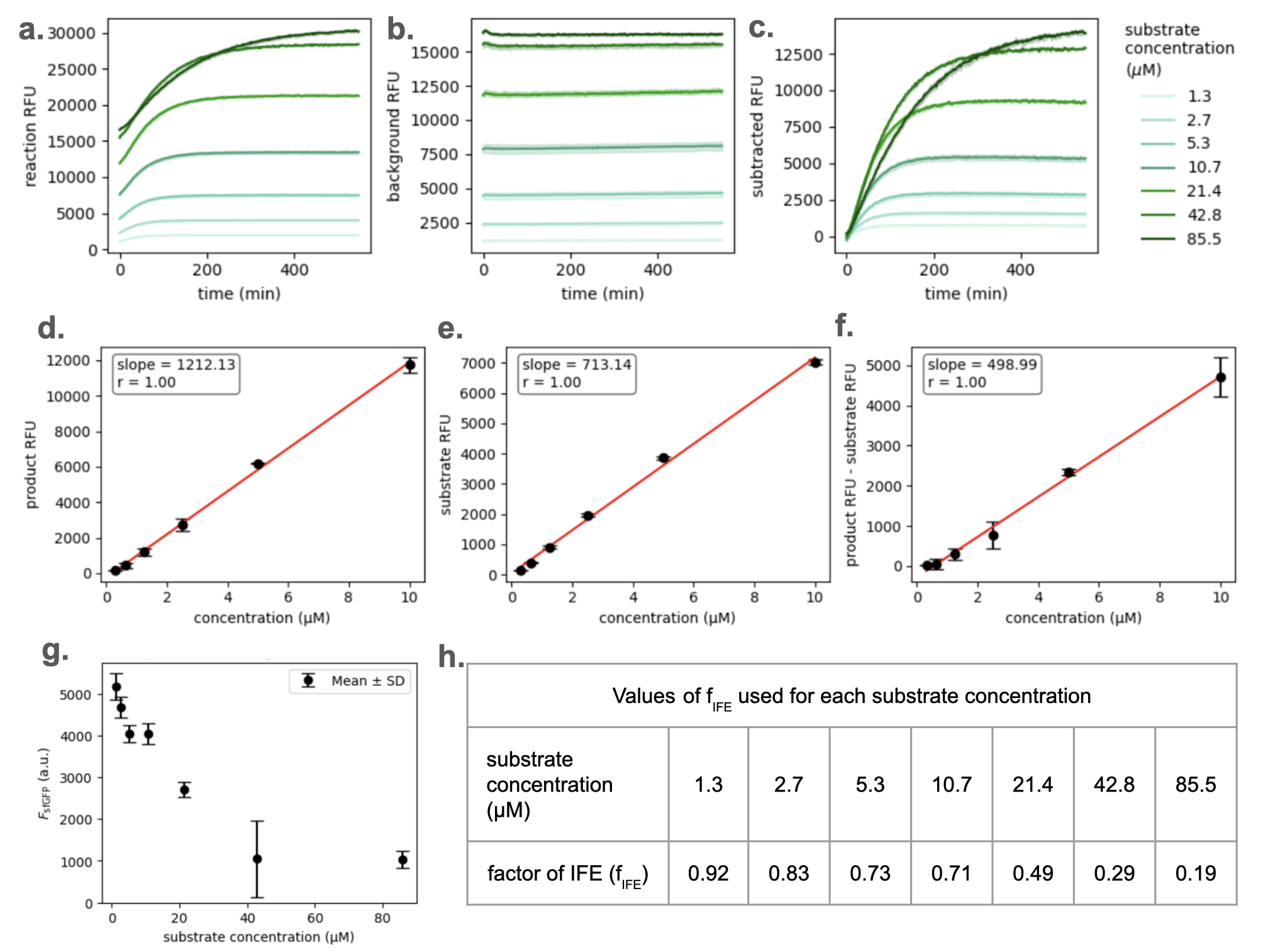

##### Figure S16. Progress curves, standard curves and correction of inner filter effect for the kinetics measurements of Zn45 cleaving the substrate of ZnO36.

**a**. Reaction progress curves with 500 nM enzyme, **b.** substrate alone control, and **c.** reaction progress curve after subtracting the substrate alone control. **d**. Fluorescence standard curves of the equimolar mixture of sfGFP and mScarlet. **e**. Fluorescence standard curves of the sfGFP-S_ZnO36-mScarlet substrate. **f**. Standard curves for the fluorescence difference between the intact substrate and separated fluorophores. **g.** The contribution of fluorescence by fluorophore (sfGFP) as a function of substrate concentration for the correction of inner filter effect **h.** Factors of inner filter effect used for correction.

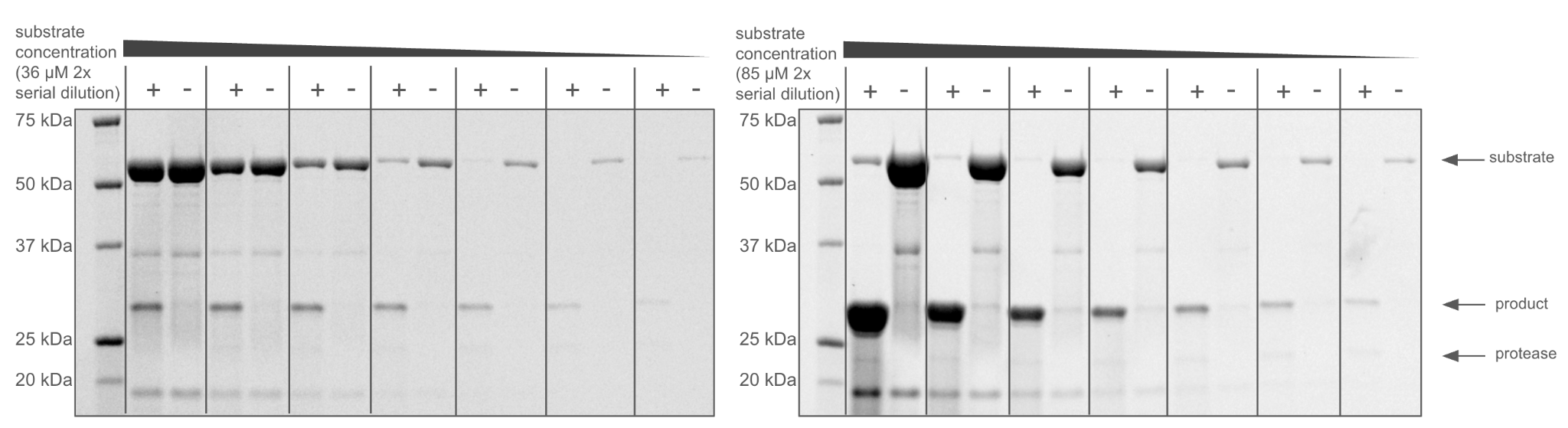

##### Figure S17. End point gels of the kinetics measurements of ZnO7 and Zn45.

Lanes are grouped in pairs of each “with enzyme group” (+) and “no enzyme group” (-) from the same substrate concentration. Because the protease concentration (0.5 μM) in both measurements is less than even the lowest substrate concentration (0.6 μM for ZnO7 and 1.3 μM for Zn45), and that the molar extinction coefficients of the proteases (16960 M^-1^cm^-1^ for ZnO7 and 25330 M^-1^cm^-1^ for Zn45 at 280 nM, the wavelength used for stain-free gel imaging) is lower than the substrate (59820 M^-1^cm^-1^ for both S_ZnO36 and S_Zn45), the protease bands are very faint.

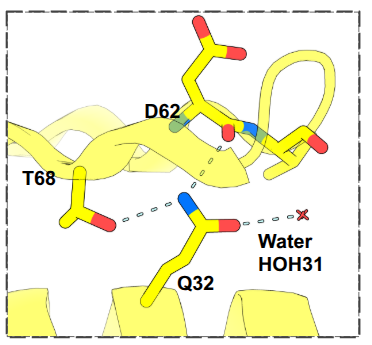

##### Figure S18. Hydrogen bonds around the Q32 residue in the crystal structure of protease-substrate complex of the E32Q variant of ZnO7.

Q32 adopts a rotamer conformation facing the surface of the protein, stabilized by three hydrogen bonds with T68, D62 and a water.

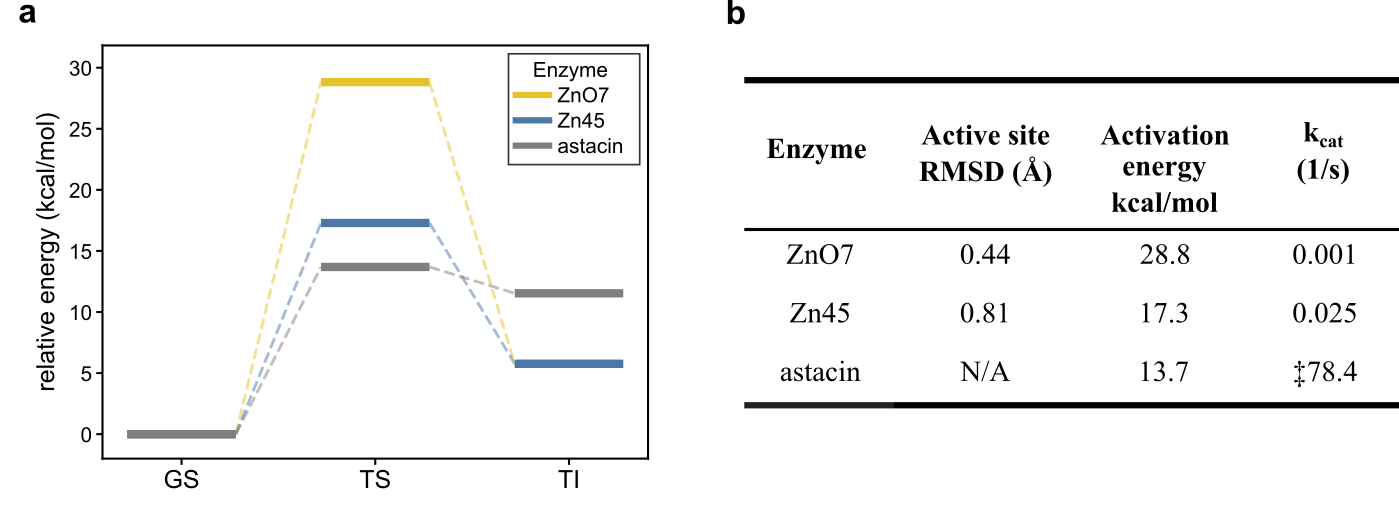

##### Figure S19. Machine learning force field calculations of the activation energy barriers of the nucleophilic attack steps of ZnO7, Zn45 and astacin.

a, relative energies calculated using MLFF with respect to the ground state (GS) of the proton abstraction and nucleophilic attack steps, for the transition state (TS) and the tetrahedral intermediate (TI). b. Active site RMSD between the design model and theozyme, activation energies and the kcat values for the proteases involved in the MLFF calculations. ‡Value reported in ref.40.

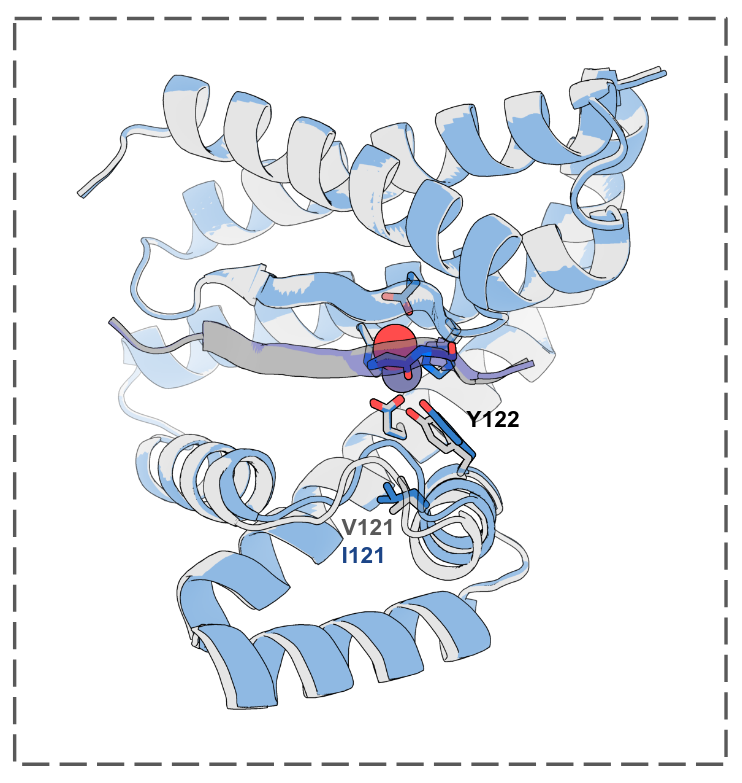

##### Figure S20. Conservative mutation V121I slightly refines the positioning of the oxyanion stabilizing tyrosine Y122.

Gray: AF3 prediction of the WT-Zn45. Blue: AF3 prediction of the V121I single variant of Zn45. Five catalytic residues as well as the V121 or I121 residues are shown in sticks representations.

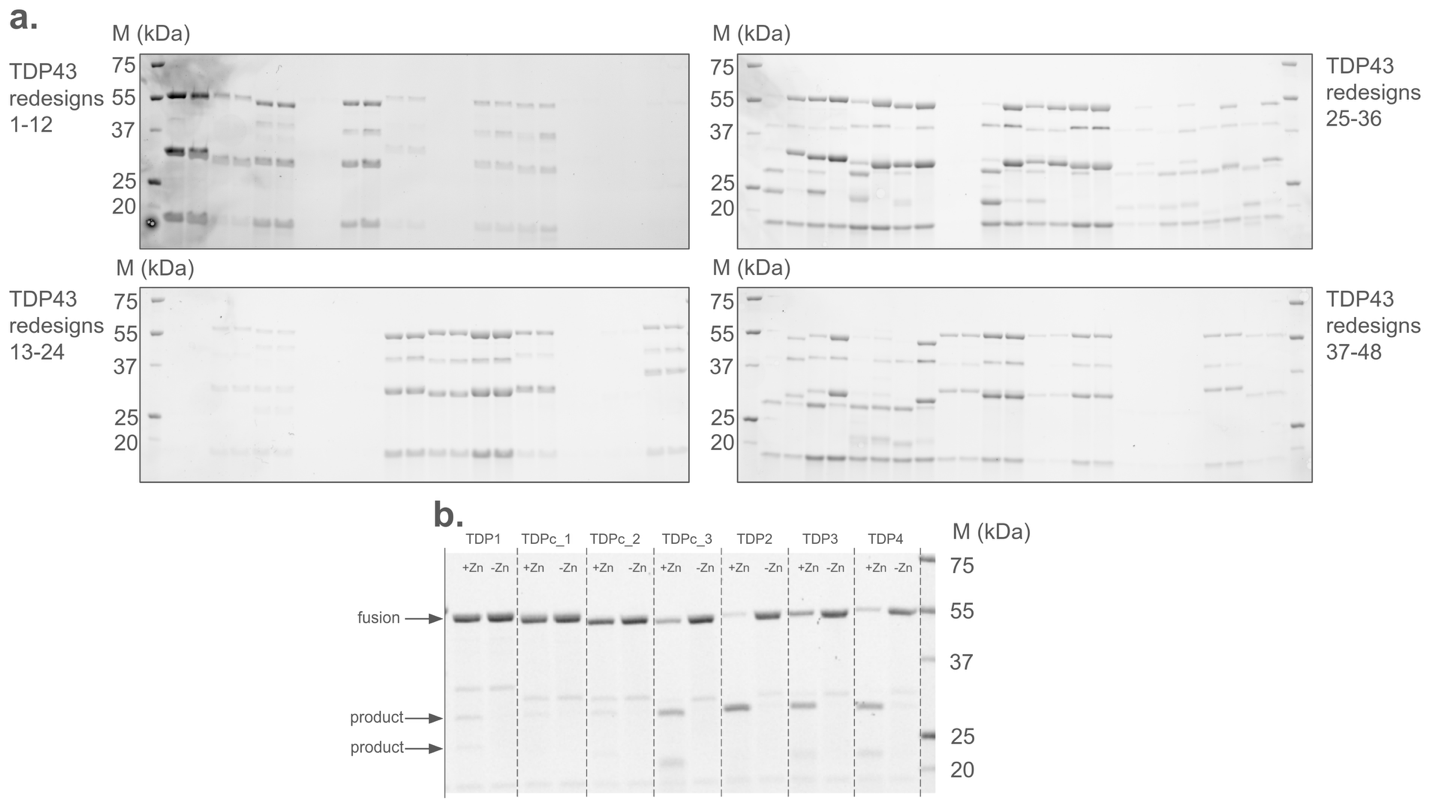

##### Figure S21. Cis screening for active redesigned proteases against human TDP-43.

**a,** 4 h time points for the 48 TDP-43 designs. Each design is shown in pairs of the +Zn and -Zn groups. **b,** 10 min time points for the 7 TDP-43 designs.

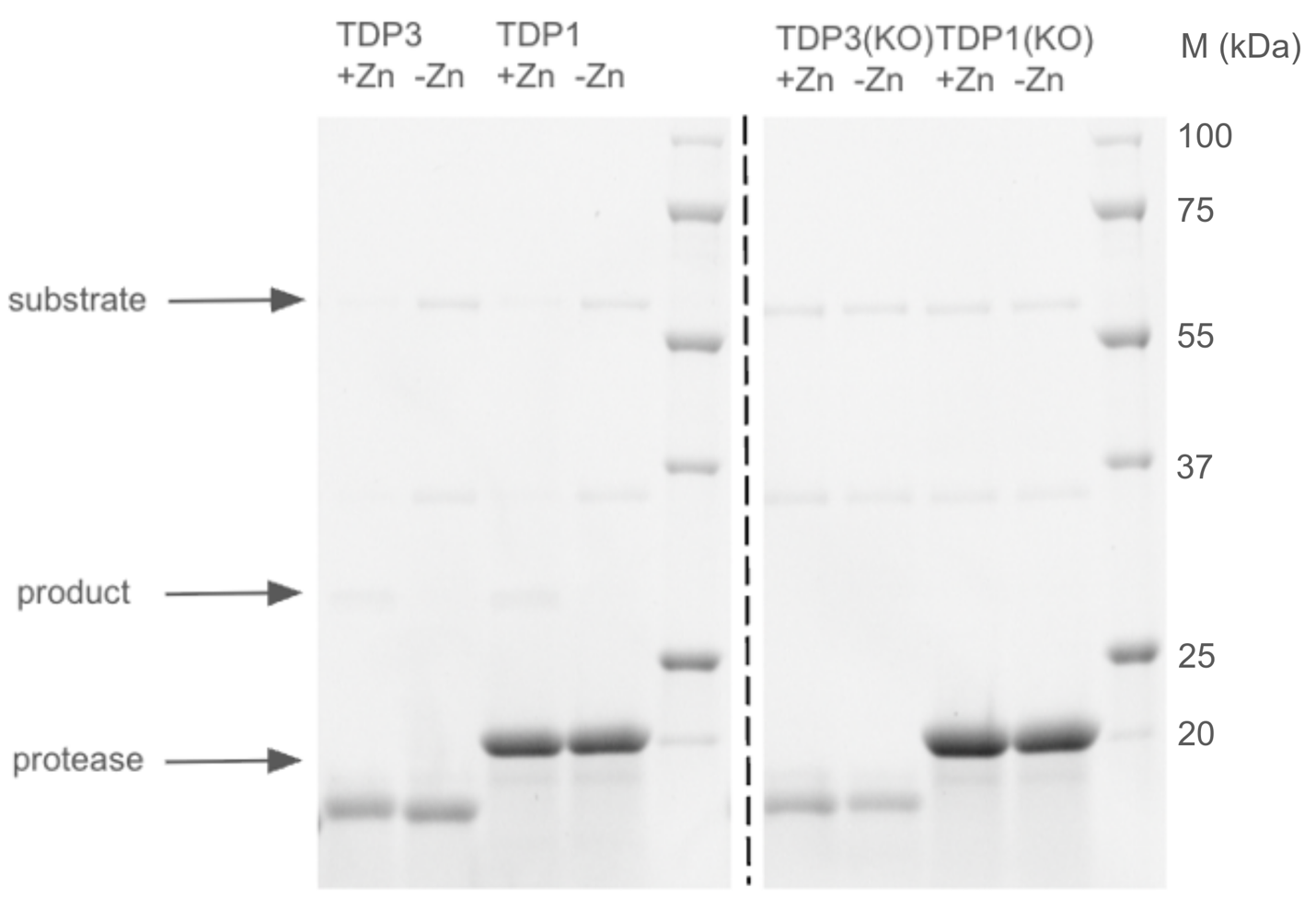

##### Figure S22. Trans screening for active redesigned proteases against human TDP-43.

Gel image showing a pair of protease + substrate + Zn and protease + substrate-Zn groups of TDP4, TDP2, TDP3, TDP1 on the left while a pair of protease + substrate+Zn and protease + substrate-Zn groups of TDP4_KO, TDP2_KO, TDP3_KO, TDP1_KO on the right.

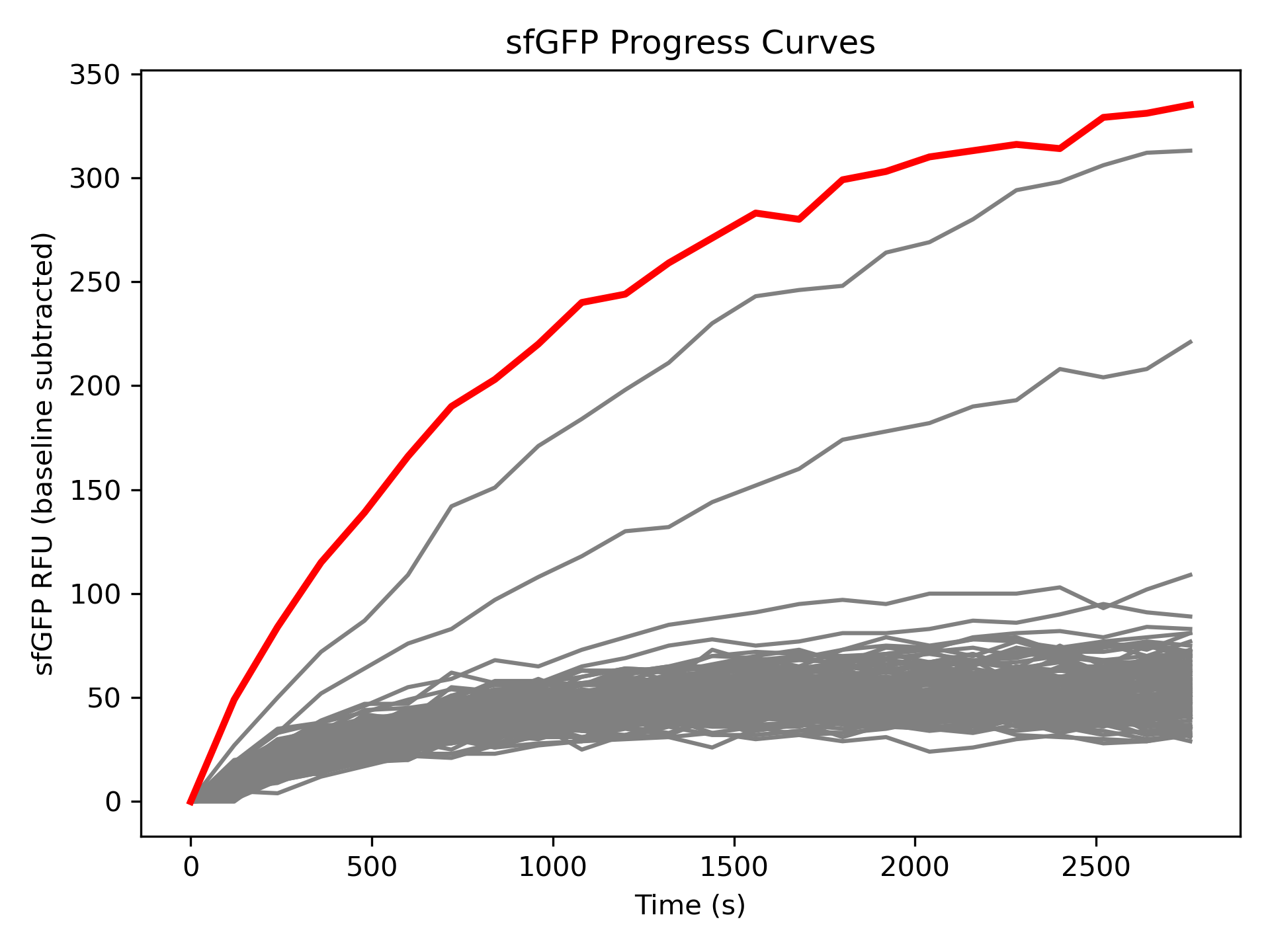

##### Figure S23. Screening for active de novo proteases against human TDP-43.

Reaction progress curves of de novo proteases cleaving the sfGFP-mScarlet substrate at 37 °C. The progress curve of TDPn3 is shown in red and the curves of the other designs are shown in gray.

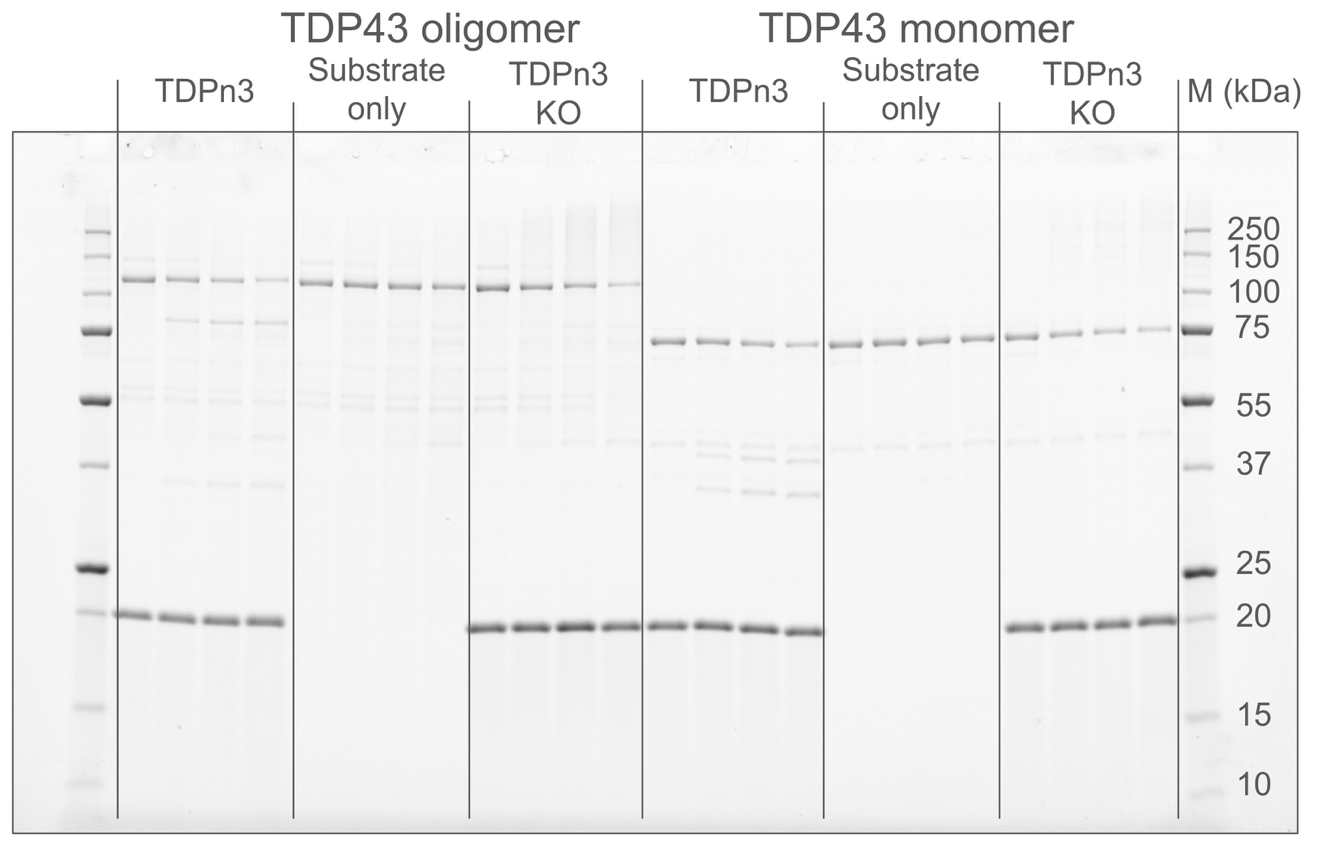

##### Figure S24. Raw SDS-PAGE gel image of TDPn3 cleavning the full-length TDP43 in the oligomer state and the monomer state.

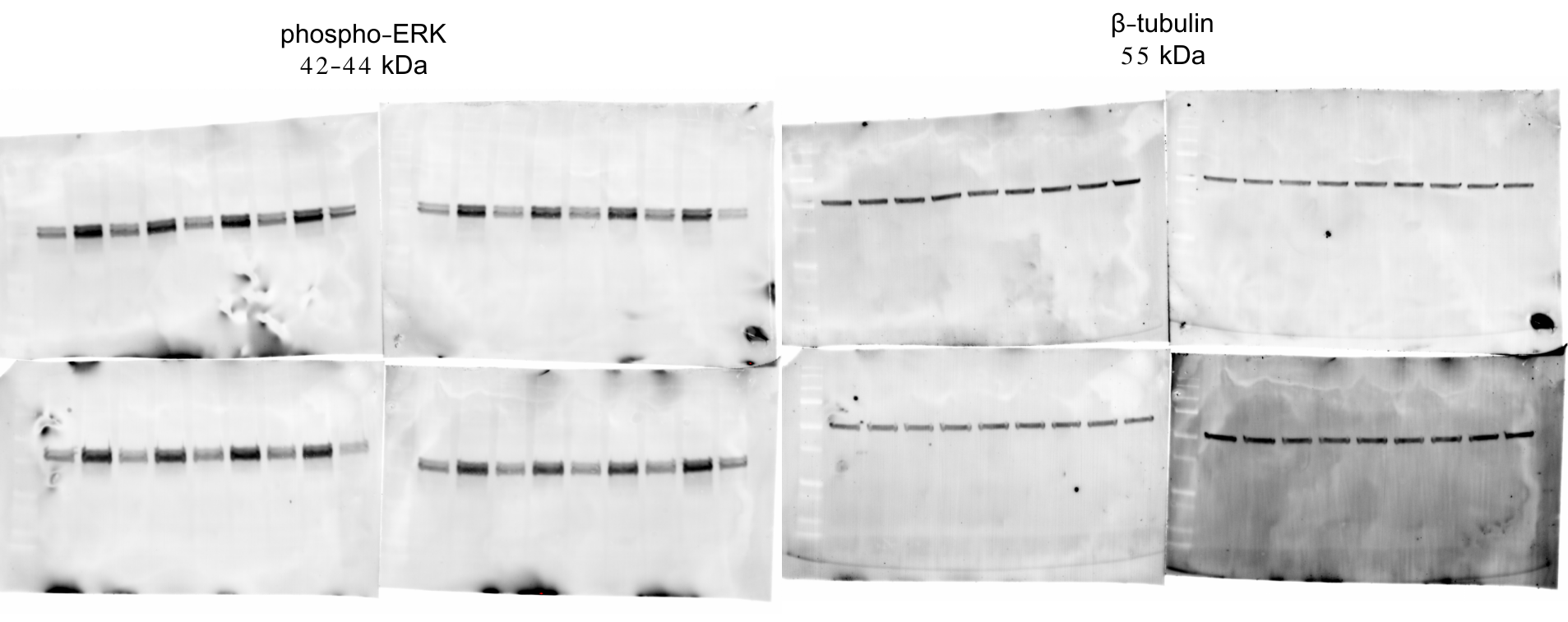

##### Figure S25. Western blot gel images for the masked EGFR antagonist assay.

From left to right: Untreated, EGF, EGFR minibinder, masked EGFR minibinder, masked EGFR minibinder with Zn45, masked EGFR minibinder with MMP2, masked EGFR minibinder with Zn45_v2, masked EGFR minibinder with Zn45_v2 KO.

### Supplementary tables

##### Table S1. Comparison between the positions of water when modeling water and Zn(II) ion as one or two entities in AF3.

The water RMSD is calculated after superposition based on all backbone Cα atoms.

| **Design name** | **Zn1** | **Zn18** | **Zn44** | **Zn45** | **Zn48** | **Zn5** | **Zn52** |
| --- | --- | --- | --- | --- | --- | --- | --- |
| **Water RMSD (Å)** | 0.26 | 0.43 | 0.19 | 0.39 | 0.33 | 0.32 | 0.16 |
| **Active in cis screen** | No | No | Yes | Yes | Yes | Yes | Yes |
| **Design name** | **Zn56** | **Zn93** | **ZnO1** | **ZnO25** | **ZnO36** | **ZnO39** | **ZnO7** |
| **Water RMSD (Å)** | 0.47 | 0.23 | 0.26 | 0.08 | 0.22 | 0.34 | 0.19 |
| **Active in cis screen** | No | Yes | No | Yes | Yes | Yes | Yes |

##### Table S2. Molecular masses of the cleavage products from the cis screen

The ~20 Da mass difference (defined as measured mass - expected mass) consistently observed across all samples is an outcome of chromophore maturation in the C-terminal mScarlet or GFP tag. This table shows the detection of the correct C-terminal product of 24 Zn designs and 36 ZnO designs. The number of designs shown in this table is larger than the number of designs reported in the main text to have functional active sites (14 Zn designs and 35 ZnO designs defined by > 10% cleavage in the cis screen after 12 h incubation at 37 °C). This likely was because the mass spectrometry experiment was performed three days after the initiation of incubation, and because mass spectrometry was sensitive enough to detect < 10% cleavage.

| Design name | Expected mass (Da) | Measured mass (Da) | Mass difference (Da) | Intended cut site | Measured cut site |
| --- | --- | --- | --- | --- | --- |
| Zn5 | 27651 | 27630 | -22 | VIRLR/MRR | VIRLR/MRR |
| Zn44 | 27577 | 27556 | -22 | VLKLT/GRR | VLKLT/GRR |
| Zn45 | 27577 | 27556 | -22 | VLEFT/GRR | VLEFT/GRR |
| Zn48 | 27515 | 27494 | -22 | PLRFR/SGY | PLRFR/SGY |
| Zn52 | 27505 | 27484 | -22 | DMTLTFR/LLA | DMTLTFR/LLA |
| Zn55 | 27532 | 27511 | -22 | SLEGILR/NPI | SLEGILR/NPI |
| Zn63 | 27683 | 27662 | -22 | TRTIRVR/YRR | TRTIRVR/YRR |
| Zn64 | 27628 | 27607 | -22 | SRVIRVR/YTR | SRVIRVR/YTR |
| Zn67 | 27640 | 27619 | -22 | SRVIEVR/YRL | SRVIEVR/YRL |
| Zn71 | 27655 | 27634 | -22 | SRTIRVR/YRK | SRTIRVR/YRK |
| Zn73 | 27683 | 27662 | -22 | SRVIEVR/YRR | SRVIEVR/YRR |
| Zn74 | 27626 | 27605 | -22 | SRVIEVR/YRV | SRVIEVR/YRV |
| Zn75 | 27640 | 27619 | -22 | SRTIRVR/YRL | SRTIRVR/YRL |
| Zn76 | 27683 | 27662 | -22 | SRVIRVR/YRR | SRVIRVR/YRR |
| Zn77 | 27683 | 27662 | -21 | SRVIRVR/YRR | SRVIRVR/YRR |
| Zn78 | 27683 | 27662 | -21 | KRTIEVR/YRR | KRTIEVR/YRR |
| Zn82 | 27683 | 27662 | -22 | SRVITVR/YRR | SRVITVR/YRR |
| Zn83 | 27683 | 27663 | -21 | SRTVEVR/YRR | SRTVEVR/YRR |
| Zn84 | 27683 | 27662 | -22 | SRTITVR/YRR | SRTITVR/YRR |
| Zn85 | 27683 | 27662 | -22 | SRVIRVR/YRR | SRVIRVR/YRR |
| Zn86 | 27683 | 27662 | -22 | SRVIEVR/YRR | SRVIEVR/YRR |
| Zn87 | 27613 | 27592 | -21 | SREIEVR/YLE | SREIEVR/YLE |
| Zn92 | 27479 | 27459 | -21 | SREVVFY/SAL | SREVVFY/SAL |
| Zn93 | 27479 | 27458 | -22 | SRRLIFY/SAL | SRRLIFY/SAL |
| ZnO2 | 27555 | 27534 | -21 | TKTLTFR/IYA | TKTLTFR/IYA |
| ZnO3 | 27507 | 27486 | -21 | TLTLTFR/IVS | TLTLTFR/IVS |
| ZnO4 | 27449 | 27428 | -21 | TLTLTFK/IGA | TLTLTFK/IGA |
| ZnO5 | 27505 | 27484 | -22 | ALTLTFR/LLA | ALTLTFR/LLA |
| ZnO6 | 27505 | 27484 | -22 | ALTLTFR/LLA | ALTLTFR/LLA |
| ZnO7 | 27590 | 27569 | -21 | ALTLTFR/LLR | ALTLTFR/LLR |
| ZnO8 | 27590 | 27569 | -21 | SLTLTFR/LLR | SLTLTFR/LLR |
| ZnO9 | 27590 | 27569 | -21 | ALTLTFR/LLR | ALTLTFR/LLR |
| ZnO10 | 27590 | 27569 | -21 | ALTLTFR/LLR | ALTLTFR/LLR |
| ZnO11 | 27533 | 27512 | -21 | SLTLTFR/LLV | SLTLTFR/LLV |
| ZnO12 | 27590 | 27569 | -22 | ALTLTFR/LLR | ALTLTFR/LLR |
| ZnO13 | 27533 | 27512 | -21 | SLRLTFR/LLV | SLRLTFR/LLV |
| ZnO14 | 27590 | 27569 | -22 | ALTLTFR/LLR | ALTLTFR/LLR |
| ZnO15 | 27533 | 27512 | -21 | ALTLTFR/LLV | ALTLTFR/LLV |
| ZnO16 | 27590 | 27569 | -21 | SLTLTFR/LLR | SLTLTFR/LLR |
| ZnO17 | 27590 | 27569 | -21 | SLTLTFR/LLR | SLTLTFR/LLR |
| ZnO18 | 27590 | 27569 | -21 | SLTLTFR/LLR | SLTLTFR/LLR |
| ZnO19 | 27590 | 27569 | -21 | SLTLTFR/LLR | SLTLTFR/LLR |
| ZnO20 | 27547 | 27526 | -21 | ALTLTFR/LLI | ALTLTFR/LLI |
| ZnO21 | 27547 | 27526 | -21 | ALTLTFR/LLL | ALTLTFR/LLL |
| ZnO22 | 27590 | 27570 | -21 | SLTLTFR/LLR | SLTLTFR/LLR |
| ZnO23 | 27590 | 27569 | -21 | SLTLTFR/LLR | SLTLTFR/LLR |
| ZnO24 | 27590 | 27569 | -21 | ALTLTFR/LLR | ALTLTFR/LLR |
| ZnO25 | 27590 | 27569 | -21 | SLTLTFR/LLR | SLTLTFR/LLR |
| ZnO26 | 27590 | 27569 | -21 | SLTLTFR/LLR | SLTLTFR/LLR |
| ZnO27 | 27547 | 27526 | -21 | ALTLTFR/LLL | ALTLTFR/LLL |
| ZnO28 | 27597 | 27576 | -21 | ALTLTFR/LLY | ALTLTFR/LLY |
| ZnO29 | 27597 | 27576 | -22 | ALTLTFR/LLY | ALTLTFR/LLY |
| ZnO30 | 27597 | 27576 | -22 | SLVLTFR/LLY | SLVLTFR/LLY |
| ZnO31 | 27620 | 27599 | -21 | ALTLTFR/LLW | ALTLTFR/LLW |
| ZnO32 | 27597 | 27576 | -21 | ALRLTFR/LLY | ALRLTFR/LLY |
| ZnO34 | 27597 | 27576 | -21 | SLTLTFR/LLY | SLTLTFR/LLY |
| ZnO35 | 27565 | 27544 | -22 | EVTGVLR/MII | EVTGVLR/MII |
| ZnO36 | 27596 | 27575 | -21 | ARILRVN/FKI | ARILRVN/FKI |
| ZnO38 | 27640 | 27619 | -22 | SRTIRVK/YLR | SRTIRVK/YLR |
| ZnO39 | 27612 | 27591 | -22 | SRVIEVK/YLK | SRVIEVK/YLK |
| TDPr3 | 28015 | 27994,28168 | -21, 153 | ALQSS/WGMMGML | ALQSS/WGMMGML  ALQ/SSWGMMGML |
| TDPn3 | 27528 | 27506 | -22 | ALQSSWG/MMGML | ALQ/SSWG/MMGML |
| TDPn3 | 40187  (C fragment of monomeric TDP-43) | 40169 | -18 | ALQSSWG/MMGML | ALQSSWG/MMGML |
| TDPn3 | 83636  (C fragment of oligomeric full-length TDP-43) | 83620 | -18 | ALQSSWG/MMGML | ALQSSWG/MMGML |

##### Table S3. Screening outcome for two-sided designs

| design name | cis screen  1 h digested fraction | cis screen  12 h digested fraction | trans screen 1 h digestion |
| --- | --- | --- | --- |
| Zn1 | 0 | 0 |  |
| Zn2 | 0 | 0 |  |
| Zn3 | 0 | 0 |  |
| Zn4 | 0 | 0 |  |
| Zn5 | 0.07 | 0.56 | 0.24 |
| Zn6 | 0 | 0 |  |
| Zn7 | 0 | 0 |  |
| Zn8 | 0 | 0 |  |
| Zn9 | wrong size | wrong size |  |
| Zn10 | 0 | 0 |  |
| Zn11 | 0 | 0 |  |
| Zn12 | 0 | 0 |  |
| Zn13 | 0 | 0 |  |
| Zn14 | 0 | 0 |  |
| Zn15 | 0 | 0 |  |
| Zn16 | 0 | 0 |  |
| Zn17 | 0 | 0 |  |
| Zn18 | 0 | 0 |  |
| Zn19 | 0 | 0 |  |
| Zn20 | 0 | 0 |  |
| Zn21 | 0 | 0 |  |
| Zn22 | 0 | 0 |  |
| Zn23 | 0 | 0 |  |
| Zn24 | 0 | 0 |  |
| Zn25 | 0 | 0 |  |
| Zn26 | 0 | 0 |  |
| Zn27 | 0 | 0 |  |
| Zn28 | 0 | 0 |  |
| Zn29 | 0 | 0 |  |
| Zn30 | 0 | 0 |  |
| Zn31 | 0 | 0 |  |
| Zn32 | 0 | 0 |  |
| Zn33 | 0 | 0 |  |
| Zn34 | 0 | 0 |  |
| Zn35 | 0.01 | 0 |  |
| Zn36 | 0.01 | 0 |  |
| Zn37 | 0 | 0 |  |
| Zn38 | 0 | 0 |  |
| Zn39 | 0 | 0 |  |
| Zn40 | 0 | 0 |  |
| Zn41 | 0 | 0 |  |
| Zn42 | 0 | 0.19 |  |
| Zn43 | 0 | 0 |  |
| Zn44 | 0.42 | 0.98 | 0.40 |
| Zn45 | 0.53 | 0.98 | 0.06 |
| Zn46 | 0 | 0 |  |
| Zn47 | 0 | 0.02 |  |
| Zn48 | 0 | 0.18 | 0.00 |
| Zn49 | 0 | 0 |  |
| Zn50 | insoluble | insoluble |  |
| Zn51 | 0 | 0.27 |  |
| Zn52 | 0.91 | 1 | 0.98 |
| Zn53 | 0 | 0 |  |
| Zn54 | 0 | 0 |  |
| Zn55 | 0 | 0 |  |
| Zn56 | 0 | 0 |  |
| Zn57 | 0 | 0 |  |
| Zn58 | 0 | 0 |  |
| Zn59 | 0 | 0 |  |
| Zn60 | 0 | 0 |  |
| Zn61 | 0 | 0 |  |
| Zn62 | insoluble | insoluble |  |
| Zn63 | 0 | 0.06 | 0.02 |
| Zn64 | 0.01 | 0.17 | 0.00 |
| Zn65 | 0 | 0 |  |
| Zn66 | 0 | 0 |  |
| Zn67 | 0 | 0 |  |
| Zn68 | 0 | 0 |  |
| Zn69 | 0 | 0 |  |
| Zn70 | wrong size | wrong size |  |
| Zn71 | 0.05 | 0.41 |  |
| Zn72 | 0 | 0 |  |
| Zn73 | 0.01 | 0.14 | 0.01 |
| Zn74 | 0.01 | 0.09 |  |
| Zn75 | 0 | 0 |  |
| Zn76 | 0 | 0 |  |
| Zn77 | 0 | 0.04 |  |
| Zn78 | 0 | 0.05 | 0.00 |
| Zn79 | 0 | 0 |  |
| Zn80 | 0 | 0 |  |
| Zn81 | 0 | 0 |  |
| Zn82 | 0.07 | 0.56 | 0.02 |
| Zn83 | 0 | 0.02 |  |
| Zn84 | 0 | 0 |  |
| Zn85 | 0 | 0.04 |  |
| Zn86 | 0.1 | 0.72 | 0.01 |
| Zn87 | 0.02 | 0.24 |  |
| Zn88 | 0 | 0.01 |  |
| Zn89 | 0 | 0.06 |  |
| Zn90 | 0 | 0 |  |
| Zn91 | 0 | 0 |  |
| Zn92 | 0 | 0.03 |  |
| Zn93 | 0 | 0.13 |  |
| Zn94 | wrong size | wrong size |  |
| Zn95 | wrong size | wrong size |  |
| ZnO1 | 0 | 0 |  |
| ZnO2 | 0.13 | 0.86 |  |
| ZnO3 | 0.08 | 0.72 |  |
| ZnO4 | -0.01 | 0.1 |  |
| ZnO5 | 0.81 | 0.99 |  |
| ZnO6 | 0.53 | 1 |  |
| ZnO7 | 0.98 | 1 | 0.95 |
| ZnO8 | 0.37 | 1 |  |
| ZnO9 | 0.84 | 1 | 0.88 |
| ZnO10 | 0.32 | 0.99 |  |
| ZnO11 | 0.72 | 1 | 0.73 |
| ZnO12 | 0.11 | 0.78 |  |
| ZnO13 | 0.53 | 1 |  |
| ZnO14 | 0.78 | 1 | 0.80 |
| ZnO15 | 0.18 | 0.75 |  |
| ZnO16 | 0.37 | 0.99 |  |
| ZnO17 | 0.11 | 0.93 |  |
| ZnO18 | 0.34 | 0.98 |  |
| ZnO19 | 0.53 | 0.99 |  |
| ZnO20 | 0.57 | 0.99 |  |
| ZnO21 | 0.84 | 0.99 | 1.00 |
| ZnO22 | 0.26 | 0.97 |  |
| ZnO23 | 0.79 | 1 | 0.95 |
| ZnO24 | 0.1 | 0.75 |  |
| ZnO25 | 0.95 | 0.99 | 0.92 |
| ZnO26 | 0.23 | 0.95 |  |
| ZnO27 | 0.78 | 1 |  |
| ZnO28 | 0.94 | 1 | 1.00 |
| ZnO29 | 0.45 | 0.98 |  |
| ZnO30 | 0.57 | 0.99 |  |
| ZnO31 | 0.9 | 0.96 |  |
| ZnO32 | 0.99 | 0.99 | 0.60 |
| ZnO33 | insoluble | insoluble |  |
| ZnO34 | 0.71 | 0.99 |  |
| ZnO35 | 0 | 0.06 |  |
| ZnO36 | 0.88 | 0.92 | 0.41 |
| ZnO37 | 0 | 0 |  |
| ZnO38 | 0.04 | 0.34 |  |
| ZnO39 | 0.68 | 1 | 0.62 |
| ZnO40 | insoluble | insoluble |  |

##### Table S4. Design sequences

| design name | protease sequence | substrate sequence |
| --- | --- | --- |
| Zn1 | MTAEELAERIGRALARGDWDSVYALGGYAFMTLSEEEKEEMIERLREVVREELARLGVELSEEEVEELVRQAVYEGEASAVVLEERRRRGLPDDLSDEELFELGMLHEAYHVNFGDSYVIADGEKGRVTVLVAETEEELREAERIAEEARREGKEVRRFAKGEREAVIEWLREVAEKYPKIREGLIEGTRRLIEEYRKIE | VIELRMPE |
| Zn2 | MTAEELAERIGRALARGRWDEVYALGGYAFRTLTEEEKEVMIERLREVVRRELAELGVELSDEEVEELVRQAVYEGEASAAVLRYREERGLPDDMTDEQLFELGMLHEAYHVNFGDSYVIADGKSGRVTVLVARTEEELEEARRIAEEREREGLEVRYFKKGETEEVIEWLREVAERYPAIRDGLVRGTRRLIERYREIV | VIELRMPE |
| Zn3 | MTAEELAERIGRALARGDWRSVYALGAYAYLTLTPEEIEEMERRLREVLREELRRRGETWSEEEVDRRVEQAIYEGKAAAVVVEEVERRGLPEDMTDEQLFELGMLHEAYHVNFGDSYVIADGEKGIVTTLIARTEEEKKEAEKIAEEAKKEGKEVKKFKKGEEEEVIEWLREVAEKYPKIREGLIEGARLLLEEYRKIR | VIELRMRR |
| Zn4 | MTAEELAERIGEALARGDWNSVYALGAYAFLTLSPEEIERMRERLREVLRERLRELGERYSEEEVDRLVEAAVMEGEAAAAVVREYEERGLPEDMTDEELFRLGMLHEAYHVRFGDAWVVADGEKGIVTVLVARTPEEEEEARRLAEEYREEGKEVRRFRRGEEEAVIEWLEEVAKKYPKIREGLIEGTRLLLEEYRKIR | VIRLRMPR |
| Zn5 | MTAEELAERIGEALARGRWDEVYALGAYAFLTLTPEEIEEMRRRLREVLREELKKLGKTYSDEEVDRLVEAAVYEGEASAVVVRRYREEGLPEDMTDEQLFELGMLHEAYHVNFGDAYVVADGKEGIVEVLVARTEEELEEARRLAERAREEGKEVRFFKKGEEEAVIEWLREVAEKYPKVREGLIEGTRRLLEEYRKIV | VIRLRMRR |
| Zn6 | MTAEELAERIGRALAEGDWNSVYALGAYAFLTLSPEEQRVMIERLREVVREELRERGERLSEEEVDRLVEQAVMEGEASAVVVREREERGLPDDMTDEELFELGMLHEAEHVNFGDAYVIADGERGIVTVLVARTEEELREAERLAEEARRQGKEVRRFRRGEREAVIEWLREVAERYPKIREGLVRGARLLLEELRKIR | VIELRMRR |
| Zn7 | MTAEELAERIGEAAARGRWDEVYALGALAYLELTPEEIEEMERRFREVLRERLAELGETLSEEEIDRLAEQAFMEGRAAAVVVERYREEGLPEDLTDEQLFELGMLHEAAHVRFGDAYVVADGRTGRVTVLVARTEEEKEEARKLAEEARAEGKEVREFARGEEEAVVEWLREVAERYPAIRDGLVRGARLLLEEYRKYV | VIELRMLP |
| Zn8 | MTPEELGRRLAEALAAGDWREVYILGAYVVLYRSEEEQEEIWEIARERLRELLAERGEEVSEEEVEEIIEIARMEGIANATMVRRARELGLGEEITDEEARELVALHEATHLRFGPGVVVVDPEKGTITVWLARTEEEKEELREQMKRWEAEGKVTRWFERGEVEEAREWILRQMRENPKVAENAAKAGRDFLREFAKLI | VVRLTFPW |
| Zn9 | MTPEELGRRLAEALAAGDWRTVYILGAYVVLYYTPEEQERIWEIARERLRELLREEGREVSEAEVDEIIDIARREGEANATMVREAKRRGLGEEITDEEARELVALHEAYHLRYGRGFIVVDPERGTIEVWLARTEEELEELRERMKEWEAEGKITREFEKGEVEEAREWILEQMEKNPKVAERAAEAGREFLRDFSELI | VVRLTFPR |
| Zn10 | MTPEELGRRIAEALAAGNWEEVYILGAYVELYYSPEEQERIWEIARERLRELLEEQGREVTEEEVDEIIDIARREGIASATMVREAQARGLGEEITDEEARELVALHEATHLEYGEGFVVVDPERGTIEVWLARTEEELEELRERMREWEARGLVTREFRKGEVEEAREWILEQMRENPKVAENAARAGRRFLREYAKLV | VVRLTFPR |
| Zn11 | MTPEELGRRLAEALAAGDWGEVYVLGAYVFLNYTPEEQERIWEIARERLREILAAQGREVTDEEVDRIIEIARMEGEANATMVRENERAGWGTEVTDEQAVELVALHEATHLEYGEGVVVVDPEKGTITVLLAETEEEKEELKEKMKEWEAQGLVTRWFEKGEVEEAREWILEQMRANPKVYENAARQGREFLERYAELR | VIRLTFAF |
| Zn12 | MTPEELGRELARALAAGDWGRVYILGAYVFLNYTPEEQEEIWEIAREELRRLLAEQGREVSEEEVDEIIEIARMEGEANATMVREARRRGWGTEVTDEQARELVALHEAYHLRYGEGVVVVDPERGTITVLLARSEEEKEELLRQMREWEARGLITRWFEKGEVEEARRWILEQMRANPRVAENAARQGRRFLEEYAKLI | IVRLTFAF |
| Zn13 | MTPEELGRELAEALAAGDWGRVYILGAYVELNYTPEEQERIWEIAREELRRLLRERGEEVTEEEVERIIEIARIEGKANATMVRANRERGWGTEVTDEQARELVALHEATHLEEGPGVVVVDPEKGTIRVLLAETEEEEERLREEMKRWEAEGLLTRWFERGEVERAREWILEQMRANPRVAENAARQGKRFLEEYAKLV | VRLLTFSF |
| Zn14 | MTPEELGRRIAEALAAGRWGEVYILGAYVFLNYSPEEQEEIWRIAREELREILRRQGREVTDEEVDEIVEIARMEGEANATMVRENQERGLGTEVTDEEARELVALHEAYHLRYGEGVVVVDPESGTITVWLARTEEEREELERRMREWEAQGKLTRWFERGEVEEARRWILEQMRANPRVAENAARQGKRFLEEYAKLV | VVRLTFAF |
| Zn15 | MTPEELGRRLAEALAAGDWKTVYILGAYVFLNYSPEEQERIWEIARERLRELLAEQGREVSEEEVERIVEIARMEGEANATMVRENERRGLGTEVTDEEAVELVALHEAYHLRYGEGVVVVDPEKGTITVLLAETEEEKEELLELMKEWEAEGKLTRWFKKGEVEEAREWILEQMRANPKVYENAARQGREFLREYAELI | IVLLTFAF |
| Zn16 | MTPEELGRRLAEALAAGDWGTVYILGAYVELYYSPEEQERIWEIAREELRRLLAEQGREVTDEEVDEIIEIARMEGIANATMVRENERRGLGTRVTDEEARELVALHEATHLEFGPGVVVVDPEKGTITVLLARTEEEREELRERMKEWEAEGKITREFERGQVEEAREWILEQMRANPKVAENAARQGKRFLEEYAKLV | VVRLTFAF |
| Zn17 | MTPEELGRQLAEALAAGDWGRVYILGAYVVLNYSEEEQREIWRIARERLRELLRERGEEVTEEEVEEIIRIAEMEGRANATMVRANEERGLGTEVTDEEARELVALHEATHLEEGDGVVVVDPEKGTITVWLARTEEEREELREQMKRWEAEGKLTRWFRRGEVEEAREWILRQMEANPKVAENAARQGKEFLREYAKLV | IVRLTFAF |
| Zn18 | MTPEELGRRLAEALAAGDWGEVYILGAYVFLNYSPEEQERIWEIAREELRRLLAEQGREVTEEEVDRIIEIARREGEANATMVRENERRGLGTEVTDEEAVELVALHEATHLRYGEGVVVVDPERGTIEVLLAETEEERRELEERMREWEAQGLLTRWFERGEVEEAREWILEQMRANPRVYERAARQGREFLRRYAELV | VVRLTFSF |
| Zn19 | MTPEELGRRLAEALAAGRWGEVYILGAYVFLYYSEEEIEEIYEIARRELREILARQGREVTEEEVDRIIEIARMEGEANATMVRENQRRGWGTEVTDEQARELVALHEATHLEYGEGVVVVDPERGTIEVLLAETEEEKEELREEMKEWEARGLVTRWFRKGEVEEAREWILEQMRRYPRVAENAARQGRRFLERFAELV | VIRLTFAF |
| Zn20 | MTPEELGRELAEALAAGDWGRVYILGAYVFLYYSPEEQEEIWRIARERLRELLRERGEEVTEEEVDEIVEIARMEGEANATMVRANRERGLGTEVTDEEARELVALHEATHLRYGRGVVVVDPEKGTIEVRLARTEEEEEELRREMEEWEAEGKITRWFERGEVEEARRWILEQMKANPKVAENAARQGREFLREFAKLI | VVRLTFAF |
| Zn21 | MTPEELGRELARALAAGDWGRVYILGAYVVLYYTPEEQEEIWEIARRELRRLLREQGRKVSEEEVDEIIEIARMEGEANATMVREAERRGWGTEVTDEQARELVALHEAYHLKYGEGVVVVDPERGTIEVRLARTEEEKEELEREMERWRAEGLLTRWFKKGEVEEAREWILEQMRANPRVARNAAEQGRRFLREFAKLV | VIRLTFAF |
| Zn22 | MTPEELGRRLAEALAAGDWGTVYILGAYVVLNYSPEEQERIWEIARERLRELLRERGEEVTEEEVDRIIEIARMEGEANATMVRANRERGWGTEVTDEEARELVALHEATHLEYGEGVVIVDPEKGTITVKLAETEEEKEELLEEMKKWEAEGKVTKWFEKGEVEKAREWILKQMAENPKVAENAAEQGRRFLEEYAKLV | IIELTFAF |
| Zn23 | MTPEELGRELAEAMAAGDWGRVYILGAYVFLNYSEEEIEEIYAIARRRLRELLAEQGREVTEEEVDRIVEIARMEGEADATMVRANRERGWGTEITDEQARELVALHEAYHLRYGEGVVVVDPERGTITVKLAETEEEKEELLREMERWEAEGLVTRYFEKGEVEEAREWILEQMDAYPRVAENAARQGREFMERFAELI | VRVLTFAF |
| Zn24 | MTPEELGRELARALAAGDWGRVYILGAYVFLNYTPEEIEEIYRIAREELRRLLAEEGREVSEEEVDEIIEIARMEGEANATMVRENERRGWGTEVTDEQARELVALHEAYHLKYGEGVVVVDPERGTITVWLARTEEEKEELLEKMKEWEAEGKITRYFEKGEVEEAREWILKQMEENPKVAENAARQGREFLEEYAKLR | VILLTFAF |
| Zn25 | MTPEELGRELARALADGDWGRVYVLGAYVVLYRSPEEQEEIYRIAREELRRILEERGEEVSDEEVDRIIEIARMEGIANATMVREARRRGLGTEMTDEEARELVALHEASHLEVGDGVVVVDPERGTITVLLAETEEEKEELDEKVKEWEAQGLITKRFEKGQTEEAREWILEQMRANPKVAENAARQGREFMREFARLV | IRVLTFAL |
| Zn26 | MTPEELGRRLAEALAAGDWGTVYVLGAYVVLYYSPEEQERIYEIARERLRELLREEGREVTEEEVDRIIEIARMEGKANALMVEEARRRGLGTEITDEEARELVALHEASHLEEGDGVVVVDPEKGTITVLLAETEEEKEELEKQMKEWEAQGLLTRWFAQGETEEARAWILKQMDENPKVAENAARQGREFLREFAKLI | IKVLTFAI |
| Zn27 | MTPEELGRELARALAAGEWGRVYVLGAYVVLNYTPEEQERIYEIARRELERLLREEGREVSEEEVDRIIEIAREEGRANATMVREAQRRGLGTDITDEEARELVALHEATHLEGGDGVVVVDPERGTVTVWLAETEEEKEKLLEKVKEWEAQGLLTRWFEQGETEAARRWILEQMEKNPKVAENAARQGREFLRRYSELV | IKKLTFAL |
| Zn28 | MTPEELGRRLAEAMAAGDWGEVYVLGAYVELYYSEEERERIYEIARERLREILEERGEKVTEEEVEERIEIARMEGIANATMVREAQARGLGTEITDEEARELVALHEATHLLWGDGVVVVDPEKGTITVLLARTEEEKEELMKLVKEWEAQGLITRWFEQGETERAREWILEQMRRNPKVAENAARQGREFLRRFAELV | IVVLTFAL |
| Zn29 | MTPEELGRRLAEALAAGRWGEVYVLGAYIVLNRSEEEQKEIYEIARRELEEILKERGEEVSEEEVEEIIEIARMEGIASAEMVREAQRRGLGTDITDEQAQELVALHEATHLEVGDGVVVVDPEKGTITVKLARTEEEKKELEEEVKRWEAEGKITKWFEQGQTEEAREWILEQMRENPKVAENAARQGKAFLEEYSKLV | VVVLTFAI |
| Zn30 | MTPEELGERLAEALAAGKWGEVYVLGAYVVLYRSPEEQERIYEIARERLRELLEERGEEVSEEEVEEIIEIARMEGEASATMVREAKERGLGTDITDEEARELVALHEATHLERGDGVVVVDPERGTITVLLARTEEEKEELMRRVKEWEAEGKLTRWFRRGETEEAREWILEQMRRNPKVAENAARQGREFLRRFAELV | VVVLTFAI |
| Zn31 | MTPEELGRRLAEALAAGDWGRVYVLGAYVVLYRTPEEQEEIYRIARERLRELLRERGEERTEEEVEEIIEIARIEGEANATMVREAQRRGLGTDITDEEARELVALHEATHLLGGDGVVVIDPEKGTITVELARTEEEWRELEERVKEWEAEGKVTRRFAQGETEEARAWILEQMRAYPKVARNAAEQGRRFLEEFAKLV | ILRLTFAI |
| Zn32 | MTPEELGRRLAEALAAGDWAEVYVLGAYVFLNRSEEEIEEIYRIAREELREILRERGEEVTEEEVEERIEIARIEGEANATMVREAQRRGIGTDITDEQARELVALHEAYHLEYGDGVVVVDPEKGTITVLLARTEEEKKELEEQMKEWEAEGKKTRWFAKGEVEEAREWILEQMRENPKVARNAAEAGRAFLRRYAELI | IVRLTFAW |
| Zn33 | MTPEELGRRLAEALAAGDWGEVYVLGAYVFLNYSPEEIEEIYRIARERLREILRERGVEMSEEEVDEIIEIARMEGEANATMVREAERRGLGKEITPEEARELIALHEATHLEYGEGVVVVDPESGTIRVLLARTEEEKQELLEQMKRWEAEGLLTRWFRKGEVEEAREWILEQMDANPRVAENAARLGKEFLERYAELI | IVVLTFGA |
| Zn34 | MTPEELGERLAEALAAGDWGEVYVLGAYIFLYRSEEEIEEIYEIARERLRELLRERGEEVSDEEVDEIIEIARREGEANATMVREAQRRGLGEEISDEEARELVALHEATHLEYGRGVVVVDPERGTIEVWLARTEEEWEELRRRMEEWEAQGLITREFERGEVERARRWILEQMRANPRVARRAAELGREFLRKFAELV | VVRLTFGA |
| Zn35 | MTPEELGRRLAEALAAGDWGEVYVLGAYVFLNYSEEEREEIYEIARRELRRLLAERGVEVSDEEVDERIEIARIEGEANATMVRKAQEWGLGEEISDEEAVELIALHEATHLRYGDGVVVVDPERGTIEVLLARTEEEKEELRRLMKEWEARGLITRWFEKGEVEEAREWILRQMRENPKVYRNAAELGRGFLRRYAELV | ILRLTFGA |
| Zn36 | MTPEELGRRLAEALAAGDWGEVYVLGAYIFLNYSPEEQEEIYRIARERLRELLRERGEEVSDEEVDEIIEIARMEGRANATMVERARERGLGEEITDEEARELVALHEATHLEFGEGVVVVDPERGTIEVLLARTEEEREELRRLMREWEARGLLTRWFERGEVEEARRWILEQMDANPRVAERAAEAGRGFLRRYAELV | IVRLTFGA |
| Zn37 | MTPEELGRELAEALAAGDWGRVYVLGAYVFLNYSEEEQKEIYEIARRELRRLLAERGEEVSDEEVDERIEIARMEGEANATMVRKAQEWGLGEEITDEEAVELIALHEAYHLRYGDGVVVIDPERGTIEVLLARTEEEKEELLKLMKEWEAQGLITRYFEKGQVEEAREWILEQMDANPKVRENAARLGKEFLRRYAELV | IVRLTFGA |
| Zn38 | MTPEELAEEIARALVEGEWGRVYVLGGLAYQELSEEDIERIEEIVEEKIKELYKEKGREITEEEARDLARRARLEGRAGGAALREAERRGLTADMSTDEGTRLMAVHEAEHLRVGDGAVVVDPETGEVRVLIARTEEEKREVREEGERLREQGYVVGYFDKDEVEKAEEFILRVAEEYPKVREELGRLGREFMRRVEERR | MLELTFRL |
| Zn39 | MTLEELARRIAEAIVAGDWASVYVLGGLAFRDLSEEEIEEVERLIERYIKELYAEEGREITEEEARERARTALIEGRAAAAALEYAERQGLTSEMSTEEGTEVMAVHEATHLEVGDGAVVINPETGEVFVLIARTEEEREEVRRIAEELREKGYVVGFFEKGQVEEAREFILEAAKKYPKVLERLGEQGREFLRRVEEAR | VLRLTFPI |
| Zn40 | MTLEELARRIAEAIVEGRWDDVYVLGGIEFRKLSEEEIERIEELIERYIKELYKEKGEEITEEEARERARIAVIEGIAAAAALEYAEEKGLTSDITVDEGTELMAVHEATHFEVGDGAVVVDPKTGKVVVLIARTEEERREVRERAEELRREGYVVRYFAKGEVEAAREAILEVAEKYPVVRERMGELGREFLRLVEERR | YLVLTFSL |
| Zn41 | FSPEVEEEAERFVERAERDIREMLARNGGRLTEEEQRRVYRYGDEAGARPSRELTYAIVRRLMERAREAFLAGEEELARAMVEAAAGALARIMADTIAEAYNLSEEMTRFVYTAVHELVHIVMLADRGTVIFRSKNTGETYEVPVDVRTMTADEFVELVLELAEKLYEMYKKDPNIEAEAVIEDYDPDRLEEGTERALRLVDEVIERL | PLILRTPV |
| Zn42 | YEELEDEIGDVAREGLERIEELEAEGRVEEAEEVRRETEFRIERLASELALRRARAEYPEELVDGGLRYIRERMERRAAETPGLTLEELRERYRAMWEAAERARPGDPSLEEPGAARIAGLHEEAHVRGIDRGVGIKKITLRLVEVEPEDERTSVEVERRED | VLRFTSPV |
| Zn43 | MGYAIFVPAEEYEEAVRKARELREQGYLARVGVGNGTEEQIEREKREEEELLDYAAERIRRDPRVRAEEVARAIAHEKVHTERLKEIYANRETPEAREKIENILIGQKSPRLKDPEVRRRTYDEERGEELIKELEEEYEKTKDPEYKGEIEVARIEVEAEREAEERTEEYLRRLRERLA | MIELTLRE |
| Zn44 | VPDAERLVEIGREEGRRGAESLAETGVATAEFPLDVPPEKARLYAEAFIEAFRERAIELVKEGRIDAETAAWAIAHEAAHAREAVELTERLARGERKEEVEKLKKYQEKVDPNDEELWRAVYDKSLEEELKKYIEEIKEKYGEEVAEGLRIYAEIEQRAEEEAEREAPRILEEIRAEVA | VLKLTGRR |
| Zn45 | MPDREELVRIGREEGRRGAESLARTGVATARFPLDVPPEQARAYAEAFIEAFRERAIELVREGRIDAETAAWAIAHEAAHARYAVELTERLARGERREEVEQIQKFQKVIDPNDEEMWEAVYDASKKEELEEYIEEIKEKYGEEVAEGLRLYAEIEQAAEREAERRAPEILREIEREVA | VLEFTGRR |
| Zn46 | SELEERVKRVIEERGPYVPLRRIAEELGVSPEEAGEALVRLVEEEGIKEALEAAASDVEYAELAARVARAYWRREGYEPGDEVGVVVVAEEDGTVRTVVTEPEEARRYAEEYGGLVKEVRLFESAEEIIELLVETIRRLKEMRPELTPEELARIGRHEALHFGLSPEEVERLAERVREAV | MIESTTRP |
| Zn47 | SFAEEVERAAERIEELIESDPELRELVERAYKSVEEGIRAAEEAVERYPELSELERRVLKTLLIAEGEAGERARELMAERLAEARRVDADPSVSPEERARVAREAIDAAPEVRPEVREAANRALDELVAQGKPIDAERMIFSAVHESLHADLFDGEIARAEWYEEDGKGFMRVYIEASEEALRRALERAEAL | MLRFRSGV |
| Zn48 | SFREEVEAAFDRILELIDSDPELRRLVDRAYESREAGIEAAREAVERYPELSPLERRVLRAYLIDEGRAGEEAREIMREALERAREVDADPTVSVEERARVAREAIRAEPRVRPEVREAVEEALDRRVAAGEPVDAAAMIFAAVHESLHAEYWDGEVVRARVYEEDGRLRMEVEIDASEEALRRRLAEAEAL | PLRFRSGY |
| Zn49 | MTLEELIELLREARENREEAEELLRRLTEVHEQLHIDRLKEEISDPEVAQEAVENATVDVERGVTTITISDEMYAKLSPEEQERIEKMVRGMEQGLRNPDPEERRLYIDMESGTLKQALEELPLSEEEKAKALVVFFERMLRAGREEEARAYSEGNPRAQRALEAVKEYL | MYTFTGRFPV |
| Zn50 | MTEEEMREALREARRNPERAEELVREFVRLHEEAHVEMIVSLFPDEELREEVRRNATVDVERGETTIRISDEDFARLPPELQERLRLLAQAGAERGRSPDPRARELYARIEGEATVRAIERMDRSPREKALTLAWLAARALEAGKEEELRNYTRLSEPAQVAARAVRAEM | MLTLTFTLVA |
| Zn51 | MTEEEMREALRRALEDPERAERLVREFIRLHEEAHVEQVVALFPDPADQEEVRRNSTIDVEAARTTIRISDELFARLPPEVQERLRLLEEAGAERGRSPDPRARELFADVEGEATVRAIERMDRTPEEKAITLTYLALQALEAGEEEALRYYTRKSEPMQVAVRAVRSML | TLTLTFTLVR |
| Zn52 | MTEEEFREALEEARRNPEEAERLVRELVRLHEEAHVDYIVSLFPDEEIREEIRRNATVDVERAVTTITISDEVFAKLPPELQERLRLLQQAGEERGRSPDPRARELYTDVESDAMVTALARWDRSPREKRIALAWMVARAVEAGRLEELDVYAAKNEELQVARRAVLAEM | DMTLTFRLLA |
| Zn53 | ATMEEIREVLEWAKRNPEEWREWARELLREHEEEHARIIREYLKDPEDRANVTTERELREEDGLLRGETRVTVSPEASPEVRERVEGLRLYAEGRTEEATPEQVAFAEEIEARAAGRVMERNAEERARAMLFFAMVAAAFRRLGLDERGRAYARGAGVTEEQIELVEREL | ATVGRLVSYV |
| Zn54 | MTVEQARELYRWAKENPEEWREKLRRILRRHEEAHTRLLMELIKDEKDRELVTTEVRLREEDGLLTGETVVRVDERASEELKRLVEGMALYSERRFDEMSPEDLEAVLRLEREANRLVVEELKEERDEVLFVAALSAAAWRRLGYDEVGRAYAEGAGVSEEVIEEVERLV | MNEGVLRNVV |
| Zn55 | MTVEEARRLYEWAQANPEEWREVLREILERHEEAHVELINEMIKDEEDRKKVTTRIELTEEDGRLSGRTVVRVDESASEEVKRFVRALKLYAELRFDEMTPEELEEVLRLEREAYRRVVERLADRRDEVQFVAALSAVTWRRLGYPEVGRAYAEGAGVSEEVVEEVERLI | SLEGILRNPI |
| Zn56 | SETYERARRAIAEAAAEATTRTVTVRTKEEMDDVIKAAEEDPRVSREEVIALKHERLHLDYSDEVTLEITADEEGRVRIR | PIRLRFPR |
| Zn57 | SETYEKAKRAIDEAAENATTRYVTVRTKEEMDEVIKAAEEDPRVSREQVIGLRHERRHLDYADEVELAISADENGNVSVSGRSTVSPEGLARIRADPAVQRALEIIRRRLPPELVPKGEEVVIQAYIEAKENNKLKEEYAEEREEIERLIEE | AVKLRFPV |
| Zn58 | SPMYEQAKRAIEEAAENASTRTVVVRTKEEMDEVIRAAEENPRVSREEVIALRHERLHLDYADEVEIEITADENGNIRVSGRSEVSEEGLEKIRKDPIVQEALRIIRERLPEELVPEGEKVVIQSYIEAKENNELIEEYAEEAEEIRELIRR | AIELRFPR |
| Zn59 | SRVYEEAVRAIEEAARDATTRKVRVTTKEEMDEVIEAAEADPRVSEDEVIALRHERLHLDYSDAVEIEISADEEGRVRVRGRSEVSEEGLERIRADPAVQEALAIIRERLPPELVPEGEKVVIQSYIEAKENNQLIREYADEAERIRRLIEE | AITLRFPR |
| Zn60 | SKTYEEARRAIEEAAENATSRTVTVRTKEEMDDVIEAAREDPRVSEEEVIALLHERLHLDYSDKVELTISADAEGRVTIE | PVTLRFPR |
| Zn61 | SEMYEKARRAIEEAARDATTRTVTVRTEEEMDEVIKAAEEDPRVSRDEVIALRHERLHLKYSDEVEITISADEEGRVRVSGRSEVSEKGLEEIRKDPAVQKALAIIRERLPEELVPEGEKVVIRSYIEAKENNELIEEYKEEAEEIRRLIEE | PITLRFPR |
| Zn62 | PAVEVAREMAAELVRLMEEAGRPEEGRRYAERIVANAPKYADNVEGFGTLGGAIAIKRARPELPFGVALGVAAALTREEPEYVEGMANLVGREVVEATMEEGERLAQRMLEEDDREIRLAAARAHEETHARILNGRGYNTVTVVLDNGEGRVIATESEEEAERIEEEERERYGDRAEVRRYRAR | MRILEFRGRF |
| Zn63 | GEEEAFEEMKRFVKEVAGIEVETREEAIEKLLELAYNPETGEAVIELLTRLAERYDLEELRDLALQAALELEAERIEREVLALPDAERAEDVVVEMALEALRAAGEDEAVERLERLWERARSDPELRRALLHEAIHVVIAREGGWRIRVEVDEESGRVRVDIEVPDDTTVDDVREAARRARERLWEEIAK | TRTIRVRYRR |
| Zn64 | SREEAFEEMKRFVKEVAGIEVETEEEAIEELLRLSYDPRTAERVIELLERLAERYELEELKDLALQAALEMEAERIRREILALPDKERAEEVVVEMALEALREAGRDDAVRRLEALWERAKSDPELRISLLHEAIHVVIARRGGWDVDVEVDEESGRVRVSITVPDDTTVEDVRAAAEKAREEIWERIAK | SRVIRVRYTR |
| Zn65 | GEEEAFEEMKEFVREVAGVEVETREEALRELRRLAYDPETADRVIELLERLARRYDLEELREEALAAALEREAERITREILALPDRERAEDVVVEMALEALEREGREEEVEELRELWERAKTDPELRVAILHESVHVVIADEGGWAVDVEVDKESGKIRVDITVPDDTTVDDIYAAADRARERIWERIAK | ARTIRVRYTL |
| Zn66 | SEEEAFEEMRRFVREVSGREVETREEALEELMRLAYNPETAERVIELLERLAKKYDLEELKKLALQAALEYEAVRIEDRILAMPDADRAEDVVVEMAVEALEARGREEEVEKLRRLWERARSDPALRRALLHEAIHVVIAAEGGWDVEVEVDEESGRVRVRIEVPDDTTVEDVREAAERARRYIEEQIAK | SRVIEVRYYL |
| Zn67 | MEEEAFERMREFVRERAGREVRTREEAIEELMELAYDPATADEVIRLLEELAERYDLEELRDLALQAALEAEAMRIEDEILALPDRERAEDVVVEMALRALERAGREDEVERMRRLWERAKTDPELRRALLHEAIHVVIAEEGGWRVRVRVDEENGRVRVEIEVPDDTTVEDIREAARRARERIEREIAR | SRVIEVRYRL |
| Zn68 | SREEAFEEMRRFVREVAGVEVTTWEEAIERLMELAYDPETAERVIKLLEELAKKYNLEELKELALQAALEMEAERIEREILALPDAERAEDVVVEMALEALRRRGREEEVERMRRLWERAKDDPELRRALLHEAIHVVIADEGGWEVRVRVDEEKGRVRVEIEVPDDTTVDDVRAAARRARERIWERIAE | SRVIEVRYYR |
| Zn69 | SREEAFEEMRRFVREVAGVEVRTFEEAVEKLMELAYNPETADRVIELLERLAERYDLEELRDLALQAALEAEAERIERKIRALPDAERAEDVVVEMAIEALERRGREEEVEKIRELWEKAKSDPELRRAILHEAIHVVIADEGGWEVRVEVDEEKGKVRVRIEVPDDTTVDDVREAAERAERRIEERIAE | SRVIEVRYKI |
| Zn70 | SREEAFEEMRRFVLEVAGKEVRTEEEALAELMRLAYDPATADRVIELLERLAEKYDLEELRVLALQAALEAEAEEIERRIRALPDAERAEDVVVEMAVEALRAEGREDEVERIRRLWERARDDPELRRALLHEAIHVVIAREGGWRVRVEVDEESGRVRVRIEVPDDTTVEDVREAARRARRRIEERIAE | SRVVEVRYRR |
| Zn71 | RREEAFERMKEFVLEVAGVEVETEEEAIEELLRLSYDPETADRVIELLERLAEEYDLEELRDLALQAALEREAERIQREIEALPDPERAEDVVVEMALEALRAQGREDAVEKLRRLWEEAKTDPELRRAILHEAIHVVIADEGGWAVDVEVDEESRKIRVRITVPDDTTVADVYAAAARARERILARIAE | SRTIRVRYRK |
| Zn72 | KREEAFEEMKEFVKEVSGIEVETFEEAIEELRKLSYNEETAQDVIDLLQELAEKYDLEELRIEALAAALEREAMKIERQVLALPDPERARDVVVEEAIRALEARGREEAVEALRRLWEEARDDPALARALLHEAIHVVIADEGGWDVDVEVDEESRRVRVRITVPDDTTVEDIERAAERARERLLRRIAE | SRVVEVRYVE |
| Zn73 | SREEAFEEMKEFVRRVAGVEVETEEEAREELMRLAYDPATAEAVIRLLEELARRYDMEELRDLALQAALEMEAERIRDRILALPDADRAEDVVVEMALEALRRRGQDEAVERMERLWEVAKDDPELRRALLHEAIHVVIADEGGWEVEVEIDEESRRVRVRITVPDDTTVEDVRAAAERARERIEAEIAK | SRVIEVRYRR |
| Zn74 | KEEEAFEEMKEFVKEVAGIEVKTREEAIKELMRLSYDPATAERVIRLLEELARKYDLEELRDLALQAALEMEAERIRDEIRALPDAERAEEVVVEMALEVLERRGQEEAVRRLRRLWEEAKDDPELRRALLHEAIHVVIADEGGWRVEVEVDEESRRVRVRIEVPDDTTVEDVRAAAERARERIMARIAE | SRVIEVRYRV |
| Zn75 | GREEAFEEMKRFVLEVSGVEVETEEEAIEELLRLSYDPETADRVIELLERLAEKYDMEELKDLALQAALEAEAERIEDEILALPDAERAIDVVVEMALEALRAAGEDEAVERLERLWERAREDPELRRAILHEAIHVVIAAQGGWRVRVEVDEESGRIRVDIEVPDDTTVADVRAAADRAREDIFARIAA | SRTIRVRYRL |
| Zn76 | NREKAFEEMKRFVKEVAGKEVETYEEAVEELLRLSYDPETAERVIELLEELARKYKMEELKRLALQAALEMEAERIRREIEALPDRERAIDVVVEMALEALRRAGRDDAVRRLERLWERAKSDPELRRAILHEAIHVVIADEGGWEVDVRVDEESGRIEVDITVPDDTTVEDIRAAARRARERLFARIAK | SRVIRVRYRR |
| Zn77 | GEEEAFERMKEFVREVAGKEVETREEAIEWLLELSYDPETADRVIALLEELAEEYDLEELRDLALQAALEREAELIEREIRALPDAERAEDVVVEMALEALRAEGREEEVERLRRLWERARSDPELRRAILHEAIHVVIAREGGWEVEVEVDEESGRIRVRITVPDDTTVEDVREAARRARERIFERIAK | SRVIRVRYRR |
| Zn78 | SEEEAFEEMRRFVLERAGVEVETREEAIEELMRLAYDPETADDVIELLERLAERYDLEELRELALQAALEREAERIEREIRALPDAERAEDVVVEMAIEALERRGREDEVERIRRLWERARDDPELRRALLHEAIHVVIADEGGWDVRVEVDEERGRVRVFIEVPDDTTVEDIERAAERARERIWARIAE | KRTIEVRYRR |
| Zn79 | SREEAFEEMRRFVRERAGVEVETEEEALAELRRLAYDPETADEVIELLERLAEKYDLEELKKEALAAALEAEAERIERRILALPDRERAIDVVVEMAVEALEAAGREEAVRRLRRLWERAKSDPELRRSLLHEAIHVVIAREGGWRVRVRVDEEKGRVRVEIEVPDDTTVEDVREAAERARRYIEERIAE | SRVIEVRYYR |
| Zn80 | SEEEAFEEMREFVLRVAGIEVETREEAIEELMRLAYDPATADEVIRLLEELAEKYNMEELKKLALQAALEAEAERIQREIEALPDRERAIDVVVEMALEALRARGREDEVRRLEELWERARDDPELRRALLHEAIHVVIADEGGWEVRVEVDEEKGRVRVFIEVPDDTTVEDVRAAAERARERIWERIAE | SRTIEVRYYL |
| Zn81 | SREEAFEEMKEFVRRVAGIEVETEEEAIEKLLELAYDPATADEVIELLERLAEKYDLEEYRRLALQAALEAEAERIRDEILALPDRERAIDVVVEMALEALEAAGREDEVERLRRLWERARSDPELRRALLHEAIHVVIAAEGGWRVRVEVDEERGRVRVRIEVPDDTTVDDVRAAAERARARIWERIAR | SRTIEVRYRR |
| Zn82 | PREEAFEEMKEFVKEVAGIEVETEEEAIEELMRLSYDPATAEDVIRLLQELAEKYDLEELRDLALQAALEMEAERIRREILALPDPERAEDVVVEAAVEALERKGREEAVEKLRELWREAQSDPELRIALLHEAIHVVIADEGGWDVEVEVDRESRRVRVRITVPDDTTVEDVREAARRARERLLRRIAE | SRVITVRYRR |
| Zn83 | RREEAFEEMKRFVKEVAGKEVKTEEEAIEELMELSYNPETAERVIELLERLAEEYDLEELRVLALQAALEMEAERIEREIRALPDPERAEDVVVEAAVEALRRRGRDDAVERLEALWEEAQSDPALRREILHEAIHVVIADEGGWDVDVEVDAESRRIRVRITVPDDTTVEDVREAARRARERLLARIAE | SRTVEVRYRR |
| Zn84 | KREEAFEEMKRFVLEVAGKEVETYEEAVEVLLELSYDPETAEEVIRLLQELAEKYQLEELRDLALQAELEREAERIRREILALPDPERAMDVVVEMAVEALRRRGRDDAVERLERLWERARDDPELRIAILHEAIHVVIADEGGWRVDVEVDEESGRVRVRIEVPDDTTVEDVRAAAERARERLLKEIAK | SRTITVRYRR |
| Zn85 | KREEAFKEMKEFVLRVSGREVETYEEAVEELLELSYDPATADEVIELLERLAEKYDLEELRDLALQAALEAEAERIEDEILALPDPERAIDVVVEMAIEALERQGKERAVEELRRLWEEAKSDPELRRAILHEAIHVVIADEGGWEVEVEVDAESRRVRVRITVPDDTTVADVRAAAARARERLLARIAE | SRVIRVRYRR |
| Zn86 | MREEAFERMREFVREVAGIEVETEEEAVEKLLELAYDPETAERVIKLLEELAEKYEMEEYKDLALQAALEMEAERIRREILAMPDAERAEDVVVELALRALEAEGQEEAVERLRRLWERAREDPELRRALLHEAIHVVIAREGGWRVRVEVDEESGRVRVRIEVPDDATVEDVRAAAARARARIEEEIAK | SRVIEVRYRR |
| Zn87 | MREEAFEEMKQFVLETAGVEVETEEEAIEKLLELSYNPETAEDVIRLLEELAEKYNLEELKDLAIQAALEMEAERIRRRILAMPDRERAEEVVVEMAVEALERRGQEEAVEKLRRLWEKAKSDPKLRIALLHEAIHVVIADEGGWEVRVEVDEESGRVTVRIRVPDDATVDDVRAAAARARERIERRIAE | SREIEVRYLE |
| Zn88 | MREEAFEEMREFVRRVAGVEVETEEEAIEELLRLAYDPATADDVIKLLEELAEKYNLEELKKLALQAALEAEAERIEDEILALPDAERAIDVVVEMALEALERRGQTEAVERMRRLWERAKDDPELRRELLHEAIHVVIADEGGWEVEVEVDEESGRVRVRITVPDDTTVEDVRAAARRARERIWEEIAK | SRTIEVRYRR |
| Zn89 | REEEAFERMREYVRRVSGVEVETREEAIAELLRLSYDPATADDVIALLEELAERYDLEELRDLALQAALEREAVEIERWILALPDKERAIDVVVEMALEVLRARGQEEAVRDLEELWEEAKDDEKLRIAILHEAIHVVIADEGGWEVEVEIDRESKRIRVRITVPDDTTVEDVRRAAERARERIERRIAE | SRVIEVRYRV |
| Zn90 | SREKYEEVVLEAAREISGREDLTLEEAREVVREEMYRADERAIRILRKFKKIIDELDAEQAAEVARMALEVVLGIEEREVIAAGELESRTEERVERTVEILEEQGDEEQVETLRRIYEKVKDDPELRRQLYHEVGHILYLRERGVEGRIEIRYDEEKKRISLKIEADLTTEQVRELARQAIEEAEELYRR | KVTFNFSSAS |
| Zn91 | MDADDLVARLVEEARKYVEEVKKLDEETQRELLELFRRLAEEMRRVAGERIADRLLELADEDPLLGMAMARAIYEGDPAVYRRWLREAEEAGNEEVAKVMRMILAEAEANEYAEARRPEYEARFADPAAVRAAVLTLHEAKHVLDETGEIDEFEVETEERPDGTVDVTIKVIGRKDDLTVIGIGRMRVPRERVPAVEEGARWGYEEI | MLLTFYSRL |
| Zn92 | MDADELVERLIKEAEKYIERVKKLDPETQERKLELFFRWAEEMRREAGPEIADRLIEIARENPLLGIAMQEAIYKGDPAVYERWLEEARAAGNERVARVLEMILDEARATRFAEARRPEYEERFADPYAVKAAVLTLHEMDHVLRRTGRVDEFRVRTERLPDGRVRVTIEVIGRRGDLTVRGIGELVVPEEQVPAVKEGARWGYENY | REVVFYSAL |
| Zn93 | MDADDLVDRLVEEARRYLDEVRRLDPETQERLLELFREWAEEMRRVAGPRIADRLLALAEEDPLLGMAMQRAIYEGDPEVYRRWLEEARAAGNEEVARVLEMVLDEAEATRYAEARRPEFEERFEDPWAVRAAVLTLHEAYHVLRRTGKIDEFRVRTRERPDGRVEVTIEVIGRKDDLEVIGIGRLVVPREQVPAVRRGAEWGYENV | RRLIFYSAL |
| Zn94 | MDADDLVAELVRLAEAYRDEVRRLPPERQRELLELFREFAEEVRRVAGPRIADRLLALADEDPVLGMAMMVAIYKGDPELFRRWLRRAREAGNEEVARVMEMILDEAEATKYAEARRPEFEEKFADPAAVEAAVLTLHEAYHVLRRTGKIDEFRVETERLPDGRVKVTIKVVGRKGDLTVIGIGRLVVPEEQVPAVEEGARYGYENI | RLLVFYSGI |
| Zn95 | LLEYGKKKGDELMELLTEPREVQIERTGRILVESYLYATKLRDPERAEELIKRFLEEKREAVREIAEELLEEVEKLKKEDEDKARIYLWFFFASEAAVSPYTTEAIERVERAREERPDLPEEEIKEIKLEGMRKHEEAHIEYAKEEGLEIKKSEIEFDLEKGVGRTTLEVDEESYEKFLEKMKKE | PMFSFFGLI |
| ZnO1 | MEEKEDEIGDIAREGLERIEELEAEGRLEEAEEVRRETELRIERLASELALEEARARYPQELVDGGLRYIRRRMEERAAATPGLTLEELRERYRALWEAAEQAAEGDPALERPGAARINGLHEMAHVRGIDRGVGIKKITLRLNEVEPEDEGTSVEVERRED | TLLLTSAA |
| ZnO2 | ATVEEFEEAYRKAREEPERAEELVRRFTEIHERIHIEQVVDFFSLPEAREEVRRNATIDVEAGRATIKISDEIYAKLPPEEQEKIDLMVEAGATFGKSPDPRARELFIRVENEALARAKEEAPFTPEEKRIVLTRLAERFLRAGEEELLEIYAREDPELRKAAEAVRRLL | TKTLTFRIYA |
| ZnO3 | PTVEEFEEAYRWALENPERAEELVREFTRIHEEIHIEQVVDLFSLPEAAEEVRRNATVDVEAGRATIRISDEIYAKLPPEEQERLRLLEEAGATNGLSPNPRARELFIEVENRALAEALERAPFTPREKRVVLTRLAERLIREGKDELLEIYSAKNEDLRKAAEAVRKLL | TLTLTFRIVS |
| ZnO4 | PTVEEFEEAYREALENPERAEELLREFIRIHEERHVEMVASLFEKPEDRKEIFENSTIDVEAGRTVINISDEVYAKLPEELQRRLDLLQEGGETVGKSPNPEAREIFREEESRAIQEALKELKLPEKEKRIILTTLAKELIERGQEELLEIYSADNPDLQKAVEAVKRLL | TLTLTFKIGA |
| ZnO5 | MTEEEVDEALERARADPARAEELVRELVRLHEQSHVDFIVSLFPDPEVQEEIRRNTTVDVEAARTTITISDEIFSKLPPELQKKLELLKEAGEKLGKSPNPEARELFVEVESQAVVDALARWPRTEEEKRIAIAWMVDRAVERGQVDRLEIYAARNEDLQVAVRTVLSRR | ALTLTFRLLA |
| ZnO6 | MTPEEVDEALAEARADPERAEALVRELVELHEQEHVDYIVSLFPDPEVQEEIRRNTTVDVEASRTTITISDEIFNRLPPELQERLELLKEAGETLGKSPDPRARELFIEVESDAVVRALERWP | ALTLTFRLLA |
| ZnO7 | MTEEEVKEALKEAEKDPARAEELVDQLVELHEQAHVDYVVSLFDDPEVQEEIRRNSEIDVERAITRITISDEIYAKLPPELQERLDLLKEAGETLGRSPDPRARELFVEVESQAVVDALKRWPISEREKNIAIAYLVKKAVEEGKEDLLPIYAAKDERLQVAVKTIKSML | ALTLTFRLLR |
| ZnO8 | MTKEEFYEALKEAEKDPERAEELVDKLIELHEQAHVEYVVSLFPDPEVQEEIRRNTTIDVEKAITTITISDEIYARLPPELQRRLDLLKEAGETEGKSPDPEARELFIEVESTAVVKALERLPISEREKLIAIAYLVKKAVEAGKADLIPIYAAKNEDLQVADEAIRSQL | SLTLTFRLLR |
| ZnO9 | MTKEEVREALRKAREDPERAERLVDQFVELHEQAHVDYVVSLFDDPEVQEEIRRNTTIDVERAITRITISDEIYARLPPELQERLRLLQEAGETRGRSPDPRARELFVEVESQAVVDALERWPISEEEKRIAIAYLVDKAVRAGKRDELPVYAARNPDLQVAVETIESEL | ALTLTFRLLR |
| ZnO10 | MTKEEVLEALKEAEKDPERAEELVRQFIELHEQAHVDFIVSLFDDPAVQEEIRENTTIDVERAVTTITISDEIYARLPPELQRRLRLLQEAGETLGRSPDPEARELFVEVESDAVVRALERWPISEREKNIAIAYLVKKAVEAGKLDLLPIYAAKDPRLQVAVETIKSQL | ALTLTFRLLR |
| ZnO11 | MTKEEVLEALEKAEEDPERAEKLVREFIRLHEQAHVDYIVSLFDDPEVQEEIRKNTTIDVEAGRTTITISDEIYAKLPKDLQERLELMKEAGETLGRSPNPKARELFIEVESTAVVDALARWPRSEEEKRIALAYLIKKAIEEGKEEDIPIYAAKNEDLQVAYETMKSLL | SLTLTFRLLV |
| ZnO12 | MTKEEVREALRKAEEDPERAERLVDEFVRLHEQEHVDFIVSLFDDPEVQEEIRENSTIDVERAITTITISDEIYAKLPPELQERLELLKEAGETLGRSPDPRARELFVEVESEAVVRALERWPRSEEEQLIAIAYLVKKAVEAGKRDLLPVYAAKDPRLQVAVETIEAEL | ALTLTFRLLR |
| ZnO13 | MTEEEFREALERARADPARAERLVDRLVELHEQAHVDYVVSLFDDPEVQEEIRKNSTIDVARGITTIEISDEIYAKLPPELQERLDLMKEAGETRGRSPDPRARELFVEVESDAVVRALERLPVSEEEKRVAIAWLVDRAVRAGRRDDLPVYAAKDEMLQVAVRAVEAEL | SLRLTFRLLV |
| ZnO14 | MTKEEVREALEKAKADPERARELVDRLIELHEQAHVDFVVSLFPDPEVQEEIRKNSVIDVERARTTITISDEIFNKLPPELQEKLELMKEAGETLGKSPNPRARELFVEVESDAVVRALERWPISEEEKLIAIAYLVDEAVKAGKEDLLPIYAAKNEMLQVAVETIREML | ALTLTFRLLR |
| ZnO15 | MTPEEVRAALAEARADPARAEALVREFIRLHEEAHVDYIVSLFDDPEVQEEIRENSTIDVERGITTITISDEIFNTLPPELQERLRLMQEAGETLGRSPDPRARELFVEVESDAVVRALERWPRSEREKNIAIAYLVDRAVRAGKRDLLPIYAAKDERLQVAVRTIEAEL | ALTLTFRLLV |
| ZnO16 | MTKEEFREALEEARRDPERAERLVDELIRLHEEAHVDFVVSLFDDPEVQEEIRKNTTIDVEAARTTITISDEIYAKLPPELQEKLDLMKEAGETLGKSPDPRARELFEEVESDAVVRALERWPRSEREKNIAIAWLVDRAVEAGREDLLPIYAKKDERLQVAVEAVRSEL | SLTLTFRLLR |
| ZnO17 | MTEEEFREALEEARRDPERAERLVRELIRLHEQEHVDYIVSLFPDPEVQEEIRRNSTIDVEAARTTITISDEIFARLPPELQERLELLKEAGETLGKSPDPRARELFVEVESDAVVRALERLPISEREQNIAIAYLVDKAVREGKEDLLPIYAAKDPRLQVAVRTIKAEL | SLTLTFRLLR |
| ZnO18 | MTEEEVDEALARARADPARAERLVDEFIRLHEEAHVDYVVSLFDDPEDQEEIRRNTTIDVEAARTTITISDELFARLPPELQERLRLMQEAGETEGRSPDPRARELFIEVESDAVVRALERWERSEEEKLIAIAYLVDRAVRAGREDLLPVYAAKNETLQVAVRAIRARL | SLTLTFRLLR |
| ZnO19 | MTKEEVLERLEEARRDPARAQALVDRFVELHEQAHVDFVVSLFDDPEIQKEIRENTTIDVERAITTITISDEIYATLPPDLQEKLDLMKEAGETLGKSPDPRARDLFVEVESQAVVDALARWPVSEREKGIAIAWLVDEAVRAGKRDLLPIYAAKDPMLQVAVETIESQL | SLTLTFRLLR |
| ZnO20 | MTEEEFYEALEEARRDPERAERLVRQLVELHEQAHVEFIVSLFDDPEVQEEIRRNSTIDVERGITRITISDEIFAKLPPELQERLRLLQEAGETRGRSPDPRARELFVEVESQAVVDALERLPISEREKDIAIAYLVDQAVRAGKEDELPVYAAKDERLQVAIRAIRSRL | ALTLTFRLLI |
| ZnO21 | MTPEEVREALRKAEEDPERAEELVREFVELHEQAHVDYVVSLFDDPEVQEEIRRNSTIDVERAITTITISDEIYARLPPELQERLDLLKEAGETLGRSPDPRARELYVEVESEAVVRALERWPRSEEEKAIAIAYLVKKAVEAGKRDLLPIYAAKDEMLQVAVRTVEAEL | ALTLTFRLLL |
| ZnO22 | MTEEEVLEALRRAEEDPERAERLVRQFIELHEQAHVDFIVSLFPDPEVQEEIRRNTTIDVERARTTITISDEIYARLPPELQERLRLMQEAGETLGRSPDPRARELFVEVESEAVVRALERLPLSEEEKNIAIAYLVDRAVREGKRDLLPVYAAKDERLQVAVRTIESRL | SLTLTFRLLR |
| ZnO23 | MTEEEFWEALERAKADPERAERLVREFIELHEQAHVDYVVSLFDDPEVQEEIRKNTTIDVEAARTTITISDEIYAKLPPELQERLELLKEAGETLGRSPDPRARDLFVEVESRAVVDALERWPRSEEEKNIAIAYLVDRAVRAGRRDLLPVYAARDERLQVAVRAVESQL | SLTLTFRLLR |
| ZnO24 | MTEEEVREALARAEEDPERAERLVRELIELHEQAHVDFIVSLFDDPEVQEEIRENSTIDVERAITTITISDEIYAKLPPELQERLRLLQEAGETRGRSPDPRARELYVEVESQAVVDALARLPRSREEKLIAIAWLVRRAVEAGKEDELPIYAAKNEDLQVAVRTIRAEL | ALTLTFRLLR |
| ZnO25 | MTKEEFREALEKAREDPERAEALVDELVRLHEQAHVDYIVSLFPDPEVQEEIRKNSTIDVEAARTTITISDEIYAKLPPELQERLNLLKEAGETRGRSPDPRARELFVEVESQAVVDALERWPISEEEKAIAIAYLVDRAVRAGREDELPVYAARNEMLQVAVEAVREEL | SLTLTFRLLR |
| ZnO26 | MTKEEVREYLERARADPARAQELVDQFIELHEQAHVEYVVSLFDDPKVQEEIRKNTRIDVERAITTITISDEIYNTLPPELQEKLDLMKEAGETLGKSPDPRARELYIEVESDAVVRALERLPISEEEKGIALAWLVWEAVEAGKEDLLPIYAAKNEDLQVAVETIREML | SLTLTFRLLR |
| ZnO27 | MTKEEFREALEKAREDPERARALVRELVRLHEQAHVDFIVSLFDDPEVQEEIRKNSTIDVERAVTRITISDEIFAKLPKDLQERLELLKEAGETRGRSPDPRARELYVEVESDAIVRALERWPRSEEEKNIAIAYLVDEAVKAGRRDELPVYAAKDERLQVAVETIESML | ALTLTFRLLL |
| ZnO28 | MTEEEFREALARAREDPARAEELVRELVRLHEQAHVDFVVSLFDDPEVQEEIRANTTIDVERAVTTITISDEVYAKLPPELQEKLRLMQEAGETLGKSPDPRARELYVEVESDAVVRALERWERSREEKAIAIAYLVDRAVRAGKVDLIPIYAAKNEDLQVAQRAVLATQ | ALTLTFRLLY |
| ZnO29 | MTKEEFEEALERARADPERAEELVRELVELHEQEHVDYVVSLFPDPKVQEEIRRNTTIDVERAVTRITISDEIFAKLPPELQERLELMKEAGETLGKSPDPRARELFIEVESDAVVRALERWERSEEEKNIAIAYLVDRAIDAGKEDLLPIYAAKDERLQVAVETVLARR | ALTLTFRLLY |
| ZnO30 | ATKEEIREALERARADPARAEELVRELVRLHEQAHVDFVVSLFPDPEVQKEIRENTTIDVERAVTRIEISDEIFAKLPPELQEKLELMKEAGETLGKSPDPRARELFIEVESDAVVRALERWERSEEEKLIAIAYMVDRAVEAGKVDLIDIYAAKNETLQVAKETILEMQ | SLVLTFRLLY |
| ZnO31 | MTEEEFREALEEARRDPERAERLVREFVRLHEQAHVDYVVSLFPDPEVQAEIRENTTIDVERAVTRITISDEVYAKLPPELQERLELMKEAGETLGRSPDPRARELYVEVESDAVVRALERWPVSEREKNIAIAYMVDRAVERGQVDLIPVYAAKDERLQVAERTVLAER | ALTLTFRLLW |
| ZnO32 | MTKEEFDEALAEARADPERAEALVRELVELHEQAHVDYIVSLFPDEEVQKEIRENTTIDVERAVTTIEISDEIYNKLPPELQKKLELLQEAGETLGRSPDPEARRLFVEVESDAVVRALERWDRSEREKAIAIAYLVDRAVEAGREDLLPIYAAKNEDLQVAVEAILARR | ALRLTFRLLY |
| ZnO33 | MTKEEFYRALEEARRDPARAEELVRRLVELHEQAHVEYVVSLFPDPEVQEEIRENTTIDVERAVTTITISDEIFERLPPELQERLRLMKEAGETRGRSPDPRARELFVEVESDAVVRALERWPVSEEEKAIAIAYMVDRAVEAGKTEDIPIYAAKNEDLQVAQEAVLARQ | ALTLTFRLLW |
| ZnO34 | VTEEEMLEALEKAKADPERAERLVRELVELHEQAHVEFVVSLFDDPEVQEEIRRNTTIDVEAARTTITISDEIFNKLPKELQERLELMKRAGEERGKSPDPRARELFIEVESDAVVRALERWP | SLTLTFRLLY |
| ZnO35 | PSLEELKELIEWAKENPEEWEERLRELALDHERNHERLFRERIKDPELREKLRFERRVRERDGKTVYETVITVPDSFPQEIKDRLRGAKLVAEGRAEEATPEQLEFVRREEREVMRELLRENEDERAEVVLFAIISYFNYRELGLEDIGRAYLEGVGVSVELAEEVEKYL | EVTGVLRMII |
| ZnO36 | SFEEEVERLVEVLRRKLLEMGLSESLAEQVGEALRRAARDDSLLESFAMLAAIARLAEEGNDEAVYVALALLYIASQTRPEYSRALEEGLSPENLAELRAFLKEVYEKYKDKITTEKVREVARAHEEAHIRWFAGRGYWGVGIIRSD | ARILRVNFKI |
| ZnO37 | MIKEKAFKLEEKAIEIWNEAKEKYEKEEEQLKFIEEELEKYLKEEYGLEEAKVRIERDGDRLYVTVRWKKDGVEFELRSYSSENTAKALGHEMAHVGLMLGLAEELSPERFKVLADALLEVTRRYLEGKLPEEVKELIIKSYSLSFDELEKMVEENRDKLTLEEILFMYVEAGLLNREEFDRFLEKMFEVLEELLKK | TFRTGAIYVP |
| ZnO38 | RREEAFEDMRRFIREVAGVEVETWEEAKEKLLELAYDPATADRVIEFLEELARKYELPELRDLALQAALEAEAMRIEREINAMPNAERAREVVIDRAIELLRERGREEDVRVLEELKERAKTDPRLQDALLHEAIHVVIADEGGWEVRVRVDEETGEVTVELTVPDDTTVADVREAARRARERLWREIEK | SRTIRVKYLR |
| ZnO39 | RREEAFEDMKRFIKEVAGKEVETWEEAVEVLIELAYDPETADRVIKFLEEAAEKYELPELRDLALRAAFEAEAMRIEREIEAMPDAERAREVVIDYAIELLRRRGREESVEVLERLREESKTNPELRTALLHEAIHVVIADEGGWDVEVRVDEETGEVTVRLEVPDDTTVEDVRRAAERARERLLERIAK | SRVIEVKYLK |
| ZnO40 | AAEEGGRLGAELARRIFEEPDMPIREVAERIFEMYTLTDNPEVRELLEREYRERPEEVMRVLEKVARELIEEWRRETGGDEELFVRLGGLFGAIEGITMQYGGEVLIEELRKIEAENPNDEEARLEAKRRGAEVHEERHEEVLRELGARVRVEERRIEEEDGVVLGKTVLEFETEEDFERMLARV | MRLLFFGII |
| TDPr1 | SEKEEFKEALERAEADPARAEELVRRFIELHEQAHVDFIVSLFPDPEIQKEIRENSVVDVAAARTTITISDEVYERLPPELQERLDLLVVAGRERGRSPDPRARELFTEVESDAVVRALERLEISEEEKKIALAWIVKRAIEAGEEEDLAIYAAKVPELQIALEAVKSTL | ALQSSWGMMGML |
| TDPr2 | SFEEEVERLAEVLREWLLRQGLSEELAEEVAEALRVAAKDPSLLESFAVLAAIARLAQEGNEKAVYVALALLYLLSQTRPEYTKALREGLSPKSQAELEKYLKEVYEKYKDDITLENIKVVTRAHEEAHIRRFASLGYWAVGVVLSDGSGLAVAAETREELERLLEELRREYPNIVEERRREPD | ALQSSWGMMGML |
| TDPr3 | SFEEEVEELAETLREYLRRLGMSERLAEEVAEALRRAARDPDLLESFAVLAAIARLAREGDELAVYVALALLYLLSQTRPEYTKALDEGLSPESKEELRRFLEEVYRRYRDDITLENIIRVTRAHEEAHIRRFASLGYYAVGIVRSDGSGLAVAAETREELDRLVAELREKYPDIIEERRREPD | ALQSSWGMMGML |
| TDPr4 | SFRERVERLAEVLRDRLREMGLSERLAEEVAEALRRAAEDPSLLESYAVLAAIAELAEEGDELTVYVALALLYLLSQTRPEYTRALREGLSPESQEELRAFLERVYEEYKDKITLENVIKVTRAHERAHIERYASLGYWAVGVVLADGSGLAVAAETREELERLLEQLRAEYPNIIEEERREPD | ALQSSWGMMGML |
| TDP_c1 | SFEERVERLAEVLRRYLLEQGLSERLAEEVAQALIDAAKDPALEETYAVLAAIAELAEEGKLEAVYVALALLYIYSKSRPEYTKALDEGLSPESLKELREYLKEVYEKYKDKITLENKKEVSWAHERAHIERYASQGYWAVGVVLADGSGLAVAAETREELDRLLEELRRRYPNIIREERREPD | ALQSSWGMMGML |
| TDP_c2 | SYEERVERLAEVLRDYLLRQGLSEKLAEEVAEALRVAAKDDSLLESYAVLAAIAELAEEGDEKAVNVALSILYLKMKTEPEYTKALNEGLSPESMKELEEYLKKVYEEYKKDITLENIKVVTRAHERAHIEYFASKGYWGVGVVFADGSGEAVAAETEEELERLLEELRRRRPDIVEEFRRRPL | ALQSSWGMMGML |
| TDP_c3 | SFEERVEELAEVLREKLREMGMSEELAEQVAEALRRAARDPDLLESYAVLAAIAELAKEGDELAVYVALALLYELSKTRPEYTRALDEGLSPEAREELEKFLEEVYEKYKDKITLENIKVVTRAHERAHIERFASLGYWAVGIVRADGSGLAVAARTEEELEELLAQLRAEYPDIIEEERRRPD | ALQSSWGMMGML |
| TDPn3 | MSLSEIFDRLVEPTLRVPRLARTVDELVAAGYDRRDAARLVAAIEALAFAGVALVAPEAYREIAEAIRELDPTAYEVYVEGAEALLAAARAGDPVAQELLERARRLGEEALTNERGKELRERIEEIVDELETRASREAVIEAATIHEYLHAALFDGEILALRVEGPWIVSEVAVPRERLDELLEKVK | ALQSSWGMMGML |
| SAAc8 | SEEEERWIENMRKTIKELFPDVDVSDALKAAREIAERLGYELVVFPSREAAREKLGFTGVTLPDDKVIYAADEELTLLHETMHALMVERPQEPPPEEVLAFIRSLEGIEATIAALALVEGNVNRGILVEIPQEVFDALSRENVEEVRRNLSPEALRIIYTATRENAVEAARRAIEDYF | SSRSFFSFLG |
| AbetaF3 | DDREELAERMRRIAEAATEEELEEGERETARLLAYLREATDTPEYRRAVEKIREALKDEDRFTQTHEVNHLYLTAALARRAGAEVYAEEVEVETETLPDGRRISRITIPVSDAEAVRRAAAELGRVMGLTGSEEALFEAVMEGLVTAMTAVEEGIPLRTLRDGPNPLYGAGGVVAALRAREVEAFLK | KGAIIGLMVG |

##### Table S5. Crystallography data collection and refinement statistics

|  | **Zn(II) complex of Zn5**  PDB Code: 11CE | **E32Q -Zn(II) - substrate complex of ZnO7**  PDB Code: 11CM | **TDPr3 E45Q apo**  PDB Code: 11CP |
| --- | --- | --- | --- |
| **Data collection** |  |  |  |
| Space group | R 3 :H | P 3_1_ 21 | P 4_1_ 2_1_ 2 |
| Cell dimensions |  |  |  |
| *a*, *b*, *c* (Å) | 108.80, 108.80, 89.07 | 66.51, 66.51, 80.47 | 80.43, 80.43, 167.03 |
| α, β, γ (˚) | 90, 90, 120 | 90, 90, 120 | 90, 90, 90 |
| Resolution (Å) | 64.73 - 3.41 (3.68 - 3.41) | 33.26 - 2.37 (2.50 - 2.37) | 33.66 - 2.70 (2.83 - 2.70) |
| *R*_sym_ or *R*_merge_ | 0.501 (1.043) | 0.655 (3.364) | 0.140 (1.144) |
| *I* / σ*I* | 4.2 (1.9) | 4.3 (1.1) | 12.6 (2.6) |
| Completeness (%) | 99.7 (99.4) | 100 (100) | 95.5 (100) |
| Redundancy | 10.1 (9.5) | 19.9 (20.0) | 12.0 (12.3) |
| **Refinement** |  |  |  |
| Resolution (Å) | 64.73 - 3.41 (4.29 - 3.41) | 33.26 - 2.37 (2.52 - 2.37) | 33.66 - 2.70 (2.91 - 2.70) |
| No. reflections | 5355 (2674) | 8718 (1414) | 15039 (3067) |
| *R*_work_ / *R*_free_ | 0.2116 (0.2315) / 0.2447 (0.2798) | 0.2301 (0.3067) / 0.2867 (0.3507) | 0.2291 (0.2688)/ 0.2733 (0.2976) |
| No. atoms |  |  |  |
| Protein | 1656 | 1454 | 3004 |
| Ligand/ion | 1 | 1 | n/a |
| Water | 7 | 44 | 17 |
| *B*-factors |  |  |  |
| Protein | 84 | 37 | 65 |
| Ligand/ion | 95 | 27 | n/a |
| Water | 49 | 33 | 57 |
| R.m.s. deviations |  |  |  |
| Bond lengths (Å) | 0.002 | 0.002 | 0.004 |
| Bond angles (°) | 0.36 | 0.46 | 0.68 |

*Single xtal used for each data/structure.

*Values in parentheses are for highest-resolution shell.

|  | **Zn5 apo**  PDB Code: 11CR | **E146Q-Zn(II)-substrate complex of Zn48**  PDB Code: 11CU | **E75Q-Zn(II) complex of Zn5**  PDB Code: 11CY | **E45Q-substrate complex of TDPr3**  PDB Code: 11DU |
| --- | --- | --- | --- | --- |
| **Data collection** |  |  |  |  |
| Space group | R 3 :H | P 2_1_ 2_1_ 2_1_ | R 3 :H | P 2_1_ |
| Cell dimensions |  |  |  |  |
| *a*, *b*, *c* (Å) | 107.33, 107.33, 89.35 | 76.94, 81.09, 105.89 | 107.86, 107.86, 88.88 | 31.76, 108.15, 93.75 |
| α, β, γ (˚) | 90, 90, 120 | 90, 90, 90 | 90, 90, 120 | 90, 91.28, 90 |
| Resolution (Å) | 64.42 - 3.57 (3.91 - 3.57) | 43.62 - 2.74 (2.89 - 2.74) | 64.39 - 2.77 (2.92 - 2.77) | 108.15 - 3.61 (3.95 - 3.61) |
| *R*_sym_ or *R*_merge_ | 0.381 (0.924) | 0.197 (3.031) | 0.097 (1.058) | 0.344 (0.990) |
| *I* / σ*I* | 3.3 (1.8) | 9.2 (1.3) | 15.0 (1.8) | 5.4 (1.9) |
| Completeness (%) | 99.8 (99.5) | 100 (100) | 99.9 (100) | 98.4 (99.5) |
| Redundancy | 10.8 (10.2) | 13.8 (14.4) | 10.5 (10.9) | 6.8 (7.0) |
| **Refinement** |  |  |  |  |
| Resolution (Å) | 64.42 - 3.57 (4.09 - 3.57) | 43.62 - 2.74 (2.82 - 2.74) | 64.39 - 2.77 (3.05 - 2.77) | 70.83 - 3.61 (3.89 - 3.61) |
| No. reflections | 4541 (1502) | 17933 (1346) | 9781 (2425) | 7209 (1453) |
| *R*_work_ / *R*_free_ | 0.2319 (0.2996) / 0.2546 (0.3122) | 0.2185 (0.3884) / 0.2716 (0.4614) | 0.2165 (0.2847) / 0.2541 (0.3708) | 0.2566 (0.2963)/ 0.2969 (0.3504) |
| No. atoms |  |  |  |  |
| Protein | 1675 | 4881 | 1662 | 6215 |
| Ligand/ion | n/a | 1 | 1 | n/a |
| Water | n/a | 7 | 3 | n/a |
| *B*-factors |  |  |  |  |
| Protein | 130 | 93 | 90 | 109 |
| Ligand/ion | n/a | 65 | 87 | n/a |
| Water | n/a | 68 | 65 | n/a |
| R.m.s. deviations |  |  |  |  |
| Bond lengths (Å) | 0.002 | 0.003 | 0.003 | 0.003 |
| Bond angles (°) | 0.42 | 0.51 | 0.45 | 0.61 |

##### Table S6. Sequence similarity between active metalloprotease designs and proteins in nature.

Design sequences were searched against the NCBI non-redundant database using BLASTP. The accession number of the best matches and the corresponding E-values are listed below.

| **Design** | **Accession number for best match** | **E-value** |
| --- | --- | --- |
| Zn5 | KAJ3117608.1 | 2.6 |
| ZnO7 | HOO77639.1 | 0.53 |
| ZnO25 | HOO77639.1 | 0.34 |
| ZnO36 | MCP4583916.1 | 2.9 |
| ZnO36R79A | MCP4583916.1 | 2.9 |
| ZnO39 | MDN5744340.1 | 1.0 |
| Zn45 | MEP7171328.1 | 24 |
| Zn45_v2 | HET9569644.1 | 33 |
| TDPr3 | WP_292068575.1 | 0.34 |
| TDPn3 | MEZ5720962.1 | 0.67 |

##### Table S7. Kinetic parameters of native zinc proteases arranged in ascending order of *k*_cat_

| **Enzyme** | **Substrate** | ***k*_cat_ (s^-1^)** | ***K*_M_ (µM)** | ***k*_cat_*/K*_M_ (M^-1^s^-1^)** | **Reference** |
| --- | --- | --- | --- | --- | --- |
| A disintegrin and metalloproteinase 10 (ADAM10) | Fluorogenic peptide | 0.04 | 17 | 2400 | 39 |
| membrane type-I metalloproteinase  (MT1-MMP) | Triple-helical substrate | 0.14 | 15.1 | 9315 | 40 |
| Matrix metalloprotease I (MMP1) | Type I collagen | 0.47 | 0.9 | 5.2x10^5^ | 41 |
| Botulinum neurotoxin A light chain   (BoNT/A LC) | SNAP-25 | 2.3 | 41 | 5.6x10^4^ | 42 |
| Angiotensin converting enzyme (ACE) | angiotensin-I | 7 | 17 | 4.1x10^5^ | 43 |
| Astacin | Bradykinin | 78.4 | 29 | 2.74x10^6^ | main text 37 |
| Thermolysin | Leucine-enkephalin | 180 | 140 | 1.29x10^6^ | main text  48 |
| Aminopeptidase N | Arg-Phe-[^3^H]anillide | 294 | 0.2 | 2.45x10^7^ | main text 38 |

##### Table S8. Sequences of full-length human TDP43

ALQSSWG/MMGML is the designed epitope, and the “/” stands for the designed cleavage site.

**Purified Oligomer** (peak1：**TDP43-linker-monoGFP-linker-MBP-Histag**)

MAAAMSEYIRVTEDENDEPIEIPSEDDGTVLLSTVTAQFPGACGLRYRNPVSQCMRGVRLVEGILHAPDAGWGNLVYVVNYPKDNKRKMDETDASSAVKVKRAVQKTSDLIVLGLPWKTTEQDLKEYFSTFGEVLMVQVKKDLKTGHSKGFGFVRFTEYETQVKVMSQRHMIDGRWCDCKLPNSKQSQDEPLRSRKVFVGRCTEDMTEDELREFFSQYGDVMDVFIPKPFRAFAFVTFADDQIAQSLCGEDLIIKGISVHISNAEPKHNSNRQLERSGRFGGNPGGFGNQGGFGNSRGGGAGLGNNQGSNMGGGMNFGAFSINPAMMAAAQAALQSSWG/MMGMLASQQNQSGPSGNNQNQGNMQREPNQAFGSGNNSYSGSNSGAAIGWGSASNAGSGSGFNGGFGSSMDSKSSGWGMGAPENLYFQGGSAAKFKETAAAKFERQHMDSGGGGSSGPSGSSASMVSKGEELFTGVVPILVELDGDVNGHKFSVSGEGEGDATYGKLTLKFICTTGKLPVPWPTLVTTLTYGVQCFSRYPDHMKQHDFFKSAMPEGYVQERTIFFKDDGNYKTRAEVKFEGDTLVNRIELKGIDFKEDGNILGHKLEYNYNSHNVYIMADKQKNGIKVNFKIRHNIEDGSVQLADHYQQNTPIGDGPVLLPDNHYLSTQSKLSKDPNEKRDHMVLLEFVTAAGITLGMDELYKTRLEVLFQGPGSSMKIEEGKLVIWINGDKGYNGLAEVGKKFEKDTGIKVTVEHPDKLEEKFPQVAATGDGPDIIFWAHDRFGGYAQSGLLAEITPDKAFQDKLYPFTWDAVRYNGKLIAYPIAVEALSLIYNKDLLPNPPKTWEEIPALDKELKAKGKSALMFNLQEPYFTWPLIAADGGYAFKYENGKYDIKDVGVDNAGAKAGLTFLVDLIKNKHMNADTDYSIAEAAFNKGETAMTINGPWAWSNIDTSKVNYGVTVLPTFKGQPSKPFVGVLSAGINAASPNKELAKEFLENYLLTDEGLEAVNKDKPLGAVALKSYEEELVKDPRIAATMENAQKGEIMPNIPQMSAFWYAVRTAVINAASGRQTVDEALKDAQTNSSSNNNNNNNNNNGSSHHHHHHSG

**Purified Monomer** (peak2：**TDP43-linker-monoGFP**)

MAAAMSEYIRVTEDENDEPIEIPSEDDGTVLLSTVTAQFPGACGLRYRNPVSQCMRGVRLVEGILHAPDAGWGNLVYVVNYPKDNKRKMDETDASSAVKVKRAVQKTSDLIVLGLPWKTTEQDLKEYFSTFGEVLMVQVKKDLKTGHSKGFGFVRFTEYETQVKVMSQRHMIDGRWCDCKLPNSKQSQDEPLRSRKVFVGRCTEDMTEDELREFFSQYGDVMDVFIPKPFRAFAFVTFADDQIAQSLCGEDLIIKGISVHISNAEPKHNSNRQLERSGRFGGNPGGFGNQGGFGNSRGGGAGLGNNQGSNMGGGMNFGAFSINPAMMAAAQAALQSSWG/MMGMLASQQNQSGPSGNNQNQGNMQREPNQAFGSGNNSYSGSNSGAAIGWGSASNAGSGSGFNGGFGSSMDSKSSGWGMGAPENLYFQGGSAAKFKETAAAKFERQHMDSGGGGSSGPSGSSASMVSKGEELFTGVVPILVELDGDVNGHKFSVSGEGEGDATYGKLTLKFICTTGKLPVPWPTLVTTLTYGVQCFSRYPDHMKQHDFFKSAMPEGYVQERTIFFKDDGNYKTRAEVKFEGDTLVNRIELKGIDFKEDGNILGHKLEYNYNSHNVYIMADKQKNGIKVNFKIRHNIEDGSVQLADHYQQNTPIGDGPVLLPDNHYLSTQSKLSKDPNEKRDHMVLLEFVTAAGITLGMDELYKTRLEVLFQ

##### Table S9. Designs sequences of masked biologics systems.

**Neo2**

PKKKIQLHAEHALYDALMILNIVKTNSPPAEEKLEDYAFNFELILEEIARLFESGDQKDEAEKAKRMKEWMKRIKTTASEDEQEEMANAIITILQSWIFS

**Neo2 mask**

AAEAAKAAIEAAKRAAERAEELGKESTPEVAKLYKKAAEASKKLAELTAKWAEDPAYADEAAEANKESTKLLLEAMEADK

**EGFRmb**

GSGDHWEEVFRWALEHLQEATQQNDPQKAKKILEEAHKWLRRELSEEEARAVVRWLKQLVDREL

**EGFRmb mask**

LEEWRRRAEEYLERLRERLR
